# Interplay of actin nematodynamics and anisotropic tension controls endothelial mechanics

**DOI:** 10.1101/2024.03.10.584287

**Authors:** Claire A. Dessalles, Nicolas Cuny, Arthur Boutillon, Paul F. Salipante, Avin Babataheri, Abdul I. Barakat, Guillaume Salbreux

## Abstract

Blood vessels expand and contract actively, while continuously experiencing dynamic external stresses from the blood flow. The mechanical response of the vessel wall is that of a composite material: its mechanical properties depend on a diverse set of cellular mechanical components, which change dynamically as cells respond to external stress. Mapping the relationship between these underlying cellular processes and emergent tissue mechanics is an on-going challenge, in particular in endothelial cells. Here we use a microstretcher mimicking the native environment of blood vessels to assess both the mechanics and cellular dynamics of an endothelial tube in response to a physiological increase in luminal pressure. The characterization of the instantaneous monolayer elasticity reveals a strain-stiffening, actin-dependent and substrate-responsive behavior. In response to a maintained pressure increase, the tissue displays a fluid-like expansion, accompanied by the reorientation of cell shape and of actin fibers. This actin-driven reorientation depends on focal adhesions and adherens junctions, two key mechanosensors. We introduce a mechanical model coupling actin fiber nematodynamics with active and elastic tension generation by actin fibers in the endothelium, which recapitulates the response to pressure of endothelial tubes.

## Introduction

The hierarchical structure of the cardiovascular system matures after the onset of the blood flow, starting from an initial microvascular meshwork in the embryo Udan et al. (2013); Hoefer et al. (2013); Lindsey et al. (2014). In the adult, small vessels regulate the blood flow actively through changes in their diameter to optimize tissue oxygenation. The deformation of these microvessels depends on the mechanical properties of their walls Segal (2005). The endothelium, the main constituent of the thin wall of microvessels, is a composite material: a tubular assembly of connected cells, wherein each cell itself is an assembly of various biological components. Consequently, the overall mechanics of this living material emerge from both the properties and the interactions of its constituents. But neither are constant. Subcellular processes respond to external stresses such as changes in wall tension, and alter the individual cells, thereby inducing dynamic adaptation in tissues Dessalles et al. (2021a). To maintain endothelial function throughout the organisms’ life, these adjustments occur over a wide range of time and length scales in both physiological and pathological cases Dessalles et al. (2021a).

While these changes initiate at the smallest scales, they propagate to the largest structures within the vasculature. In this adaptation process as well as in tissue mechanics, the primary actors are the cytoskeleton and adhesions complexes, physically connecting cells to the substrate and to neighboring cells Alert and Trepat (2020); Xi et al. (2018); Campas et al. (2023). Tissues exhibit viscoelastic behavior. This viscoelasticity stems from the cytoskeletal elements, such as supra-cellular actin networks and intermediate filaments Bonakdar et al. (2016); Latorre et al. (2018); Needleman and Dogic (2017); Harris et al. (2014), and from intercellular junctions Lång et al. (2018); Czirok et al. (2013); Campas et al. (2023) and adhesions to the substrate Vazquez et al. (2022); Pallarès et al. (2022). Besides passively resisting deformations, the actomyosin network creates contractile active stresses. Force transmission at adhesions propagate these subcellular stresses to the tissue level. On long time scales, cellular rearrangements such as intercalation, division and apoptosis influence the rheology of tissues and notably, their viscosity Hallatschek et al. (2023); Trubuil et al. (2021); Xi et al. (2018).

For tissues to adjust to variations in their mechanical environment, cells require a sensing mechanism Yap et al. (2017); Charras and Yap (2018); Dessalles et al. (2021a). For instance, when subjected to changes in experienced forces, adhesions trigger the recruitment of additional proteins, such as vinculin, forming mechanosensitive hubs. Subsequently, these proteins modulate the anchoring and force transduction of the actin filaments.

Deciphering how subcellular processes and their regulation by force sensing mechanisms are coupled across scales to give rise to active tissue mechanics is one of the current challenges in the field. To that end, *in vitro* systems have become versatile tools. To date, two main families of *in vitro* systems have proven instrumental: stretchers and mechanical testing platforms. Stretchers subject the substrate to controlled changes in length over time. Concomitant monitoring of the cellular response led to the discovery of a host of adaptation mechanisms triggered by mechanical stimulation, such as remodeling of actin fibers and junctions or cell stiffening Constantinou and Bastounis (2023); Dessalles et al. (2021a); Krishnan et al. (2012); Pourati et al. (1998); Hatami et al. (2013). In comparison, mechanical testing platforms, including systems such as micropipette aspiration or indentation, provide quantitative measurements of the material properties of various cells and tissues Efremov et al. (2021); Narasimhan et al. (2020); Trubuil et al. (2021); Xi et al. (2018). Despite the remarkable advancements achieved with these systems, addressing the coupling between the tissue mechanics and the multiscale dynamics of the mechanoresponsive components remains elusive, due to the difficulty of observing living tissue over the required spatial and temporal scales.

Here, we use our previously developed microstretcher, mimicking the native environment of blood vessels Dessalles et al. (2021b), to impose tension on a tubular endothelium through a physiological increase in luminal pressure. Over the course of multiple hours, this experimental setup allows to simultaneously monitor the response of an endothelial tube on the cell and tissue levels, under dynamically changing conditions. We show that an increase in luminal pressure induces anisotropic tension. While the tissue resists this tension elastically at short time scales, it displays a viscoelastic behavior at long time scales. In response to tension, endothelial cells dynamically align their cytoskeleton and their axis of elongation in the direction of the tension. This collective reorientation depends on the presence of intercellular junctions and focal adhesions. We show that an active nematic surface model of the endothelial tube, incorporating a two-way coupling between actin stress fibers orientation and tissue tension, recapitulates the dynamics of the tube radius and ordering under pressure.

## Results

### Endothelial tubes exhibit actin-dependent elasticity under luminal pressure

#### Anisotropic tension induced by luminal pressure

To study endothelium mechanics over several days, we used our hydrogel-based microstretcher Dessalles et al. (2021b). A cylindrical channel is templated within a soft collagen hydrogel and lined with ECs. An increase in luminal pressure leads to vessel dilation and monolayer stretching along the circumferential direction within seconds (Fig. 1A, Movie SV1). This deformation is reversible at short time scales (minutes), characteristic of an elastic behavior Dessalles et al. (2021b). To assess tissue tension, we performed laser ablation in monolayers of cells expressing the actin reporter lifeact, along the longitudinal (L) and circumferential (C) directions (Fig. 1Bi, Movie SV2), at the low pressure used for monolayer culture (150Pa) and in the minutes following the pressure increase (650Pa). The recoil velocity of the tissue post-ablation is thought to increase with tissue tension and to decrease with increasing tissue viscosity or elasticity Bonnet et al. (2012); Davis et al. (2022). The recoil velocity is doubled when the pressure is increased to *∼*650Pa, but only in the circumferential direction (Fig. 1Bii, S1A), indicating an increase in circumferential tension. In addition, in monolayers on low concentration hydrogels the recoil velocities are higher (Fig. S1A). As they are subjected to the same imposed tension, the viscoelastic properties of endothelia differ between the soft gels and the stiffer gels. Together, these results demonstrate a switch from an isotropic to an anisotropic tension upon the pressure increase and a substrate-dependent monolayer viscoelasticity.

**Figure 1.**
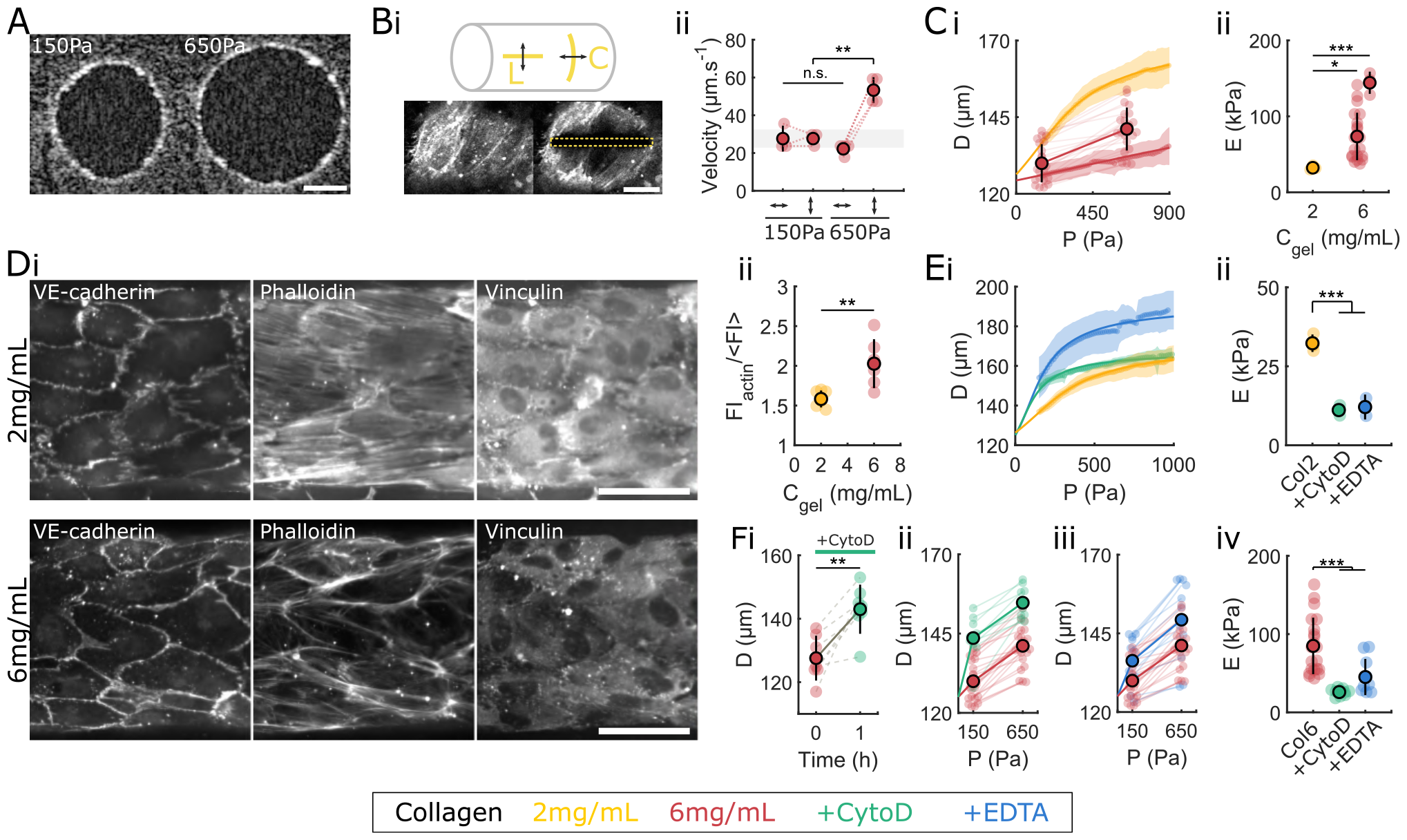
Endothelial tubes exhibit actin-dependent elasticity under luminal pressure. **A** OCT images of the vessel cross section showing the increase in radius during the increase of pressure. Scale bar, 50 µm. **Bi** Schematics of the laser ablation showing the two directions of ablation: longitudinal (L) and circumferential (C). Fluorescence images of LifeAct-ECs showing the endothelial actin network prior and post longitudinal ablation, with the area of ablation denoted in yellow, showing a rapid opening of the wound, characteristic of high tissue tension in the circumferential direction. Scale bar, 20 µm. **Bii** Initial recoil velocity post ablation for monolayers cultured on a 6 mg mL^−1^ collagen gel, showing an increase between the control (150Pa) and stretched (650Pa) channels, but only in the circumferential direction. Ablations were performed in the minutes following the pressure increase for the stretched condition. **Ci** Channel diameter as a function of the luminal pressure (points) for monolayers cultured on a 2 mg mL^−1^ (yellow) and 6 mg mL^−1^ (red) collagen gel, obtained either continuously with live imaging (chain of dots), or at the beginning and the end of the pressure application (paired dots), with the fitted analytical curves obtained from the strain-stiffening model (solid lines). **Cii** Inferred Young’s moduli of the endothelial tissue for the two collagen concentrations. For the 6 mg mL^−1^ concentration (red), data from the continuous measurement (right) and the discrete two-points measurement (left), matching the curves of panel ii, are separated for clarity. **Di** Endothelium stained for VE-cadherin, phalloidin and vinculin for the two collagen concentrations: 2 mg mL^−1^ (top) and 6 mg mL^−1^ (bottom). **Dii** Fluorescence intensity of the actin stress fibers (normalized by the mean cell intensity) as a function of collagen concentration. **Ei** Channel diameter as a function of the luminal pressure for control monolayers (yellow) and monolayers treated with cytochalasinD (green) and EDTA (blue), cultured on a 2 mg mL^−1^ collagen gel. **Eii** Inferred Young’s moduli of control, cytochalasinD-treated and EDTA-treated endothelia, cultured on a 2 mg mL^−1^ collagen gel. **Fi** Channel diameter as a function of time just after treatment with cytochalasinD (at t=0), for monolayers cultured on a 6 mg mL^−1^ collagen gel. **Fii**,**iii** Channel diameter as a function of the luminal pressure for control monolayers (red) and monolayers treated with cytochalasinD (green) and EDTA (blue), cultured on a 6 mg mL^−1^ collagen gel. **Fiv** Inferred Young’s moduli of control, cytochalasinD-treated and EDTA-treated endothelia, cultured on a 6 mg mL^−1^ collagen gel.

#### Substrate-dependent stiffness of endothelial tubes

To measure the tissue stiffness quantitatively, we recorded strain-stress curves in the physiological range. We applied an external tension on the tissue by increasing the pressure continuously from 150 Pa to 1000 Pa in one minute, and measured the deformation of the channel, either continuously with live imaging (Fig. 1C,E, Movie SV3), or at the beginning and the end of the pressure application (Fig. 1F). The maximum pressure was chosen to approximate the pressure in native capillaries, where the vessel wall is composed of a single cell layer, around 1 kPa Shore (2000), to measure the endothelium rigidity in the physiological range of mechanical load. We find the tissue stiffnesses to be around 0.13 N/m on the softer gel and 0.26-0.4 N/m on the stiffer gel, corresponding to Young’s moduli of 30 kPa and 50-120 kPa respectively (Fig. 1Cii), confirming the substrate-dependent tissue stiffening observed with the laser ablations. These Young’s modulus values are similar to those reported for suspended epithelial tissues Harris et al. (2012). In addition, the deformation-pressure curves show a strain-stiffening behavior, with a threshold strain of approximately 20% separating a linear regime at low pressures and a saturating regime at large pressures, which can be captured using a Gent model (Fig. 1Ci) Salipante et al. (2022); Pourati et al. (1998).

#### Subcellular determinants of endothelium elasticity

To understand the biological origin of the substrate-dependent mechanics, we imaged the actin network and its anchoring points. Immunostaining of the tissues show more prominent actin filaments and larger focal adhesions on stiffer gels, compared to those on the soft gels (Fig. 1D). Actin stress fibers are therefore sensitive to substrate density or mechanical stiffness and their reinforcement could underlie the substrate-dependent stiffening, in line with previous reports Harris et al. (2012); Janmey et al. (2020).

To further probe the contribution of the subcellular processes to the elastic and strain-stiffening properties, we treated the monolayers for an hour prior to mechanical testing with either cytochalasin D, an inhibitor of actin polymerization, or EDTA, a calcium chelator that perturbs adherens junctions (Fig. S1B). Both treatments lead to a significant increase in monolayer deformation for both collagen concentrations (Fig. 1Ei,Fi-ii), characteristic of tissue softening. The Young’s modulus drops to 10-20 kPa upon actin depolymerization (Fig. 1E,F), consistent with previous reports Harris et al. (2012); Pourati et al. (1998). This value is similar for both gel concentrations, confirming that actin underlies the response to substrate properties. In addition, the maximal strains of actin-depleted monolayers are identical to untreated monolayers for the low collagen concentration (Fig. 1Ei), suggesting that another cytoskeletal element controls the large deformation regime. We speculate that it could be intermediate filaments, intact in cells with depolymerized actin (Fig. S1B), as was reported previously Latorre et al. (2018); Singh et al. (2018). Perturbing adherens junctions with EDTA decreases the effective Young’s modulus of the endothelium to 15kPa and 50 kPa on the 2 mg mL^−1^ and 6 mg mL^−1^ collagen respectively (Fig. 1Eii,Fiv). The tissue likely still has some mechanical contribution despite being morcelated (Fig. S1B), as these effective moduli are much higher than that of a bare gel (50-100 Pa for the 2 mg mL^−1^ collagen and 600-1000 Pa for the 6 mg mL^−1^ collagen) Jansen et al. (2018). Individual cells may induce a local stiffening, due to their adhesions to the underlying matrix. The final strain is increased, by a value decreasing with increased collagen concentration (Fig. 1Ei,Fiii), likely due to stretching of the bare collagen between cells.

#### Cells and actin stress fibers align dynamically in the tension direction

We then sought to study endothelial tissue mechanics and long term adaptation to a change in luminal pressure, a phenomenon occurring in the native vasculature, for instance at the onset of blood flow in the embryo or due to pathologies in the adult Lindsey et al. (2014); Fegan et al. (2003); Williams et al. (1990). We therefore subjected the endothelial tube, formed at *≃*150Pa, to a fixed luminal pressure of *≃*650Pa for several days and monitored tissue and cellular response. The order of magnitude of this pressure increase mimics the initial pressurisation of the naive vascular network in embryos Lindsey et al. (2014) and the 400-900 Pa increase in capillary pressure found in hypertensive patients Fegan et al. (2003); Williams et al. (1990).

First, the diameter increases due to the rapid pressure increase, corresponding to the instantaneous elastic response described above. Over the next 56 hours, despite the fixed pressure, the diameter increases continuously, showing a fluid-like creeping behavior (Fig. 2A). When the pressure is removed after 7 hours, the diameter decreases instantaneously (Fig. 2Bi), validating the presence of tension in the tissue throughout the assay. After the application of cytochalasinD at the 7-hour time-point, the diameter increases abruptly (Fig. 2Bii), likely due to softening of the endothelial tube induced by depolymerization of the actin network (Fig. 1E,F). This further supports the notion that tension in the tissue is resisting the applied pressure during the entire duration of the experiment.

**Figure 2.**
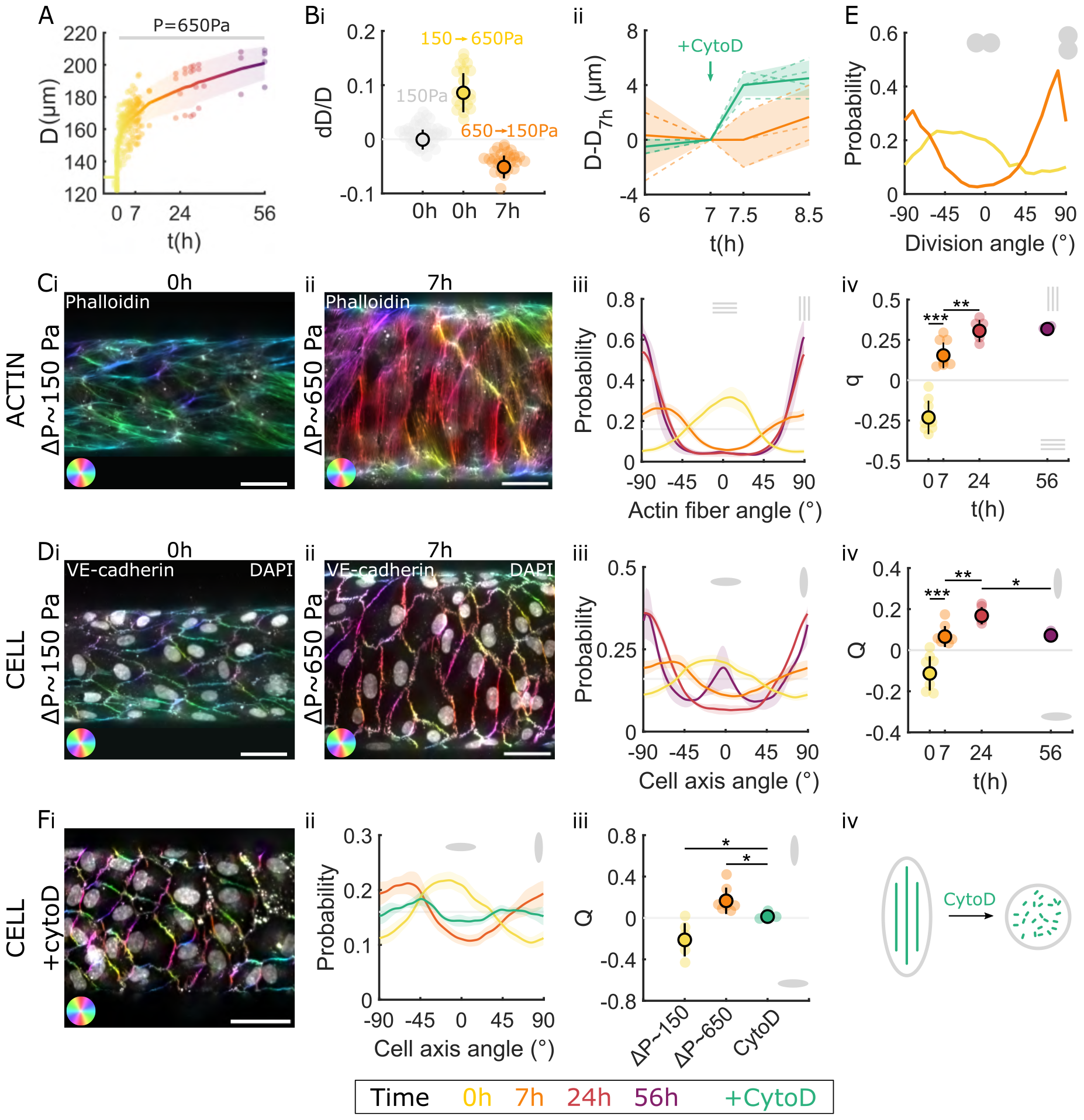
Cells dynamically align in the tension direction via an active actin-dependent process. **A** Channel diameter as a function of time after the pressure increase (t=0), color coded for time. **Bi** Relative diameter change when increasing pressure from 150Pa to 650Pa (yellow) and when decreasing pressure back to 150Pa 7 hours later (orange). The diameter fluctuations at 150Pa without any pressure change are shown in grey as a reference. **Bii** Evolution of the channel diameter between 6 and 8.5 hours for control monolayers (orange) and for monolayers treated with cytochalasin at *t* = 7h (green) under a pressure of *≃* 650Pa, showing a sudden diameter increase due to actin depolymerization. **Ci**,**ii** Endothelium stained for phalloidin at *t* = 0h under *≃* 150Pa (**i**) and after *t* = 7h under *≃* 650Pa (**ii**), with the orientation of the actin stress fibers color coded. **Ciii**,**iv** Evolution of the probability distribution of the actin stress fibers orientation (**iii**) and the associated nematic order parameter q (**iv**) at 0 (yellow), 7 (orange), 24 (red) and 56 (purple) hours. The sign of q denotes the direction of actin fibers mean orientation and its absolute value the strength of the alignment. **Di**,**ii** Endothelium stained for VE-cadherin at *t* = 0h under *≃* 150Pa (**i**) and after *t* = 7h under *≃* 650Pa (**ii**), with the orientation of the junctions color coded. Nuclei are overlaid in white. **Diii**,**iv** Evolution of the probability distribution of the cells orientation (**iii**) and the associated nematic order parameter Q (**iv**) at 0 (yellow), 7 (orange), 24 (red) and 56 (purple) hours. The sign of Q denotes the direction of cell mean elongation and the absolute value its magnitude. **E** Probability distribution of the division orientation for monolayers, measured at *t* = 7h, under low pressure Δ*P ≃* 150Pa (yellow) and high pressure Δ*P ≃* 650Pa (orange). **Fi** Cytochalasin-treated monolayer stained for VE-cadherin after 7h of pressure showing round cells. **Fii**,**iii** Evolution of the probability distribution of the cells orientation (**ii**) and the associated nematic order parameter Q (**iii**) before the pressure increase (*≃* 150Pa), and after 7h of high pressure for the control (*≃* 650Pa) and cytochalasin-treated (CytoD) monolayers. **Fiv** Schematics showing round cells after actin depolymerization by the cytochalasinD treatment, despite the circumferential stretching force. Scale bar, 50 µm.

To link the dynamics of the subcellular elements to the observed tissue flow, we imaged the actin network, cell-cell junctions and nuclei at various time points. During the assay, the actin cytoskeleton reorganizes from a longitudinal orientation to prominent stress fibers oriented in the circumferential direction (Fig. 2Ci-iii, Movies SV4, SV6). ECs and nuclei elongations follow the same dynamic pattern, from longitudinal to circumferential orientation (Fig. 2Di-iii, S2Ai, Movies SV5, SV7). In addition, the orientation of cell divisions switches from longitudinal at Δ*P ≃* 150Pa to circumferential at Δ*P ≃* 650Pa, aligning with the cell elongation axis (Fig. 2E).

To quantify these changes of orientation, we introduce the tissue nematic tensors q, Q and Q_**n**_ characterizing the collective order and orientation of actin stress fibers, cell shapes and nuclei, respectively (Methods, SI). The sign of their components along the circumferential direction, denoted *q, Q* and *Q*_*n*_, indicates whether actin stress fibers (respectively cell elongation and nuclei) are preferentially circumferential (*q >* 0, *Q >* 0, *Q*_*n*_ *>* 0) or longitudinal (*q <* 0, *Q <* 0, *Q*_*n*_ *<* 0). The magnitude of *q, Q* and *Q*_*n*_ indicates the strength of this alignment. We find that both *q, Q* and *Q*_*n*_ show significant differences at 0, 7 and 24 hours, as the actin stress fibers, the cells and the nuclei reorient progressively from the longitudinal to circumferential direction (Fig. 2Civ,Div, S2Aii). The nuclei elongation follows the cell elongation, with their order parameters showing a linear correlation (S2Aiii), and the nuclei aspect ratio showing a higher value at 24 hours, matching the peak in the cell order parameter *Q* (Fig. S2B).

#### Cell elongation and alignment is an actin-dependent active process

To disentangle whether the cell elongation is due to passive deformation by the anisotropic stretch or an active process, we treated the cells prior to the pressure increase with CytochalasinD to depolymerize actin. Adherens junctions are still present right after the treatment and after seven hours of pressure, ensuring tissue cohesion (Fig. 2Ei, S2C). CytoD-treated monolayers display round and randomly oriented cells, as quantified with the actin and nuclei orientations (Fig. 2Fi-iv, S2D), showing that the cell elongation observed in this experiment is an active process, requiring an intact actin cytoskeleton.

#### Cell-cell junctions and focal adhesions are necessary for actin alignment

Stress fibers have to be anchored to transmit the tension generated through contractility. Two main forms of stress fiber anchoring have been observed in ECs: at focal adhesions (FAs) and at adherens junctions (AJs) Han and de Rooij (2016). Here, in tissues under tension, two types of AJs are observed: classical linear adherens junctions and focal adherens junctions (Fig. 3A, S3A), known to form under tension Millán et al. (2010); Huveneers et al. (2012), that enable actin anchoring and long transcellular actin cables (Fig. S3B). Here, focal AJs are found mostly at longitudinal cell-cell interfaces (Fig. 3A), while linear AJs with parallel stress fibers are found mostly at circumferential interfaces (Fig. S3A), confirming that the tension needs to be orthogonal to the interface to trigger focal AJs formation. Interestingly, the stress fibers in focal AJs are spaced regularly (Fig. 3A), suggestive of an optimization of the distribution of the mechanical load. A similar distribution is seen in anchoring at FAs, with FAs clustered together along a line, away from the cell periphery (Fig. S3C). This line of FAs seems to organize the stress fibers as regularly spaced bundles perpendicular to its axis (Fig. 3B), a peculiar structure that has not been characterized in depth.

**Figure 3.**
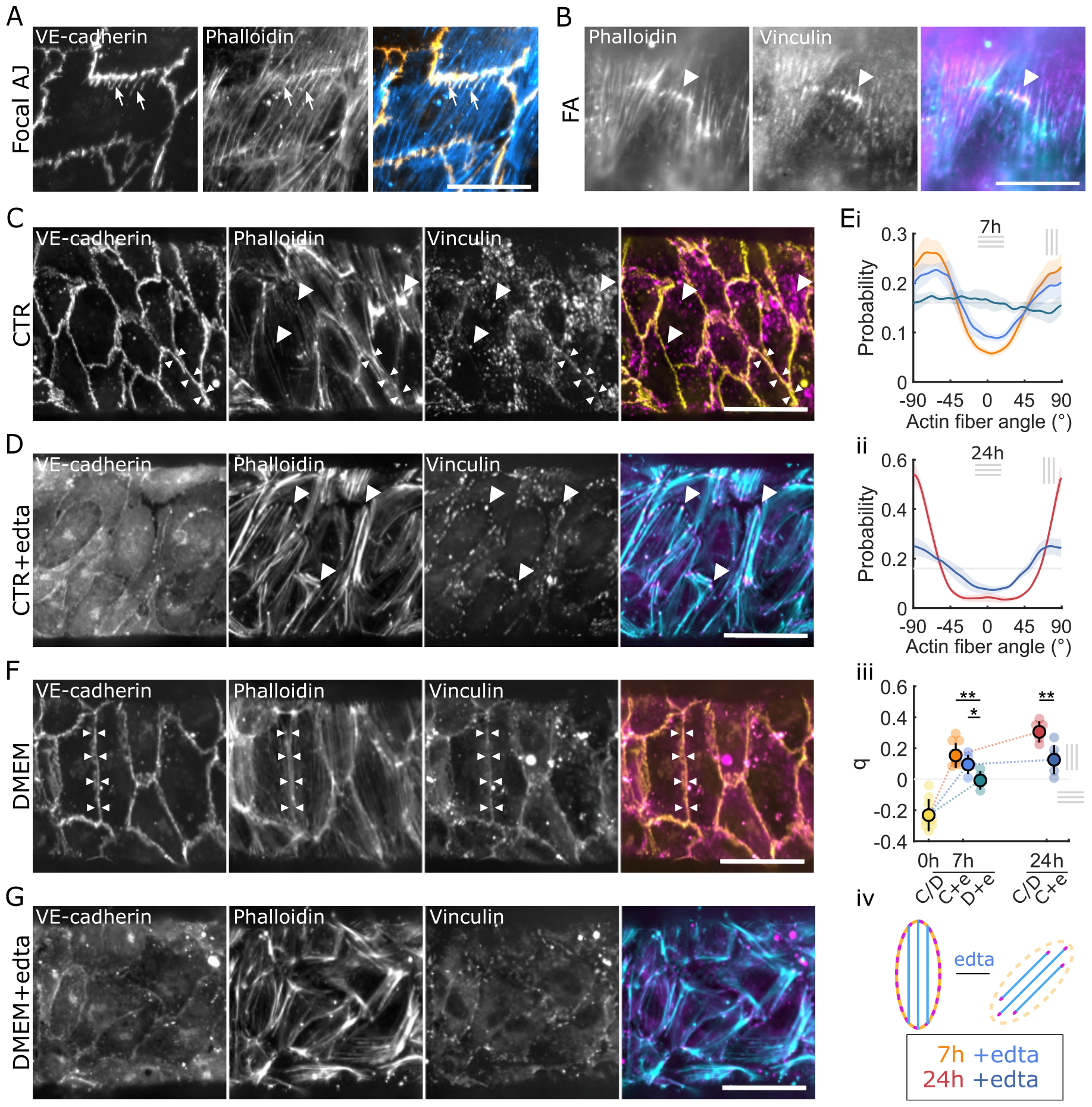
Cell-cell junctions and focal adhesions are necessary for actin alignment. **A** Endothelium stained for VE-cadherin (yellow) and phalloidin (cyan) after 7h of stretch, showing a focal adherens junction with transendothelial actin fibers association (arrows). **B** Endothelium stained for phalloidin (cyan) and vinculin (magenta) after 7h of stretch at Δ*P ≃* 650Pa, showing a line of clustered focal adhesion with actin fibers anchoring (arrowhead). Scale bar A-B, 20 µm. **C** Control endothelium stained for VE-cadherin (yellow), phalloidin and vinculin (magenta) after 7h of stretch, showing vinculin association to focal adhesions at the end of actin stress fibers (arrowheads) and to adherens junctions with parallel actin stress fibers (double arrowheads). **D** EDTA-treated endothelium stained for VE-cadherin, phalloidin (cyan) and vinculin (magenta) after 7h of stretch, showing vinculin association to focal adhesions at the ends of actin stress fibers (arrowheads). **E** Probability distribution of the actin stress fibers at 7 hours (**i**) and 24 hours (**ii**), and the associated nematic order parameter q (**iii**) for control (orange-red) and EDTA-treated (blue) endothelia. **Eiv** Schematic of a cell before and after treatment with EDTA. **F** DMEM-cultured endothelium stained for VE-cadherin (yellow), phalloidin and vinculin (magenta) after 7h of stretch, showing vinculin association to adherens junctions with parallel actin stress fibers (double arrowheads). **G** DMEM-cultured and EDTA-treated endothelium stained for VE-cadherin, phalloidin (cyan) and vinculin (magenta) after 7h of stretch. Scale bar C,D,F,G, 50 µm.

To investigate possible tension sensing by AJs, we stained for vinculin, a known mechanosensor and actin regulator, that has been previously shown to be recruited to focal AJs under tension. Vinculin colocalizes with focal AJs but also with linear AJs (Fig. 3C,F), contrary to what has been previously reported Huveneers et al. (2012). In our system, linear AJs can be subjected to tension parallel to their axis. As a result, circumferential linear AJ are also under tension, this time along their direction, and show vinculin co-localization. This is consistent with the hypothesis that vinculin is recruited by high tension in junctions Charras and Yap (2018); Huveneers et al. (2012), but suggests that this effect can happen without the remodeling into focal AJs.

To validate the putative role of junctions and FAs as mechanosensory hubs regulating actin, we first treated the monolayers prior to and during the pressure increase with EDTA to perturb AJs, while maintaining the presence of FAs (Fig. 3D). The actin network in EDTA-treated tissues after stretch showed a weaker realignment (Fig. 3D,E), indicating that AJs are involved in actin reorientation and/or tension sensing. EDTA-treated tissues continue to exhibit FAs, as shown by the vinculin dots (Fig. 3D), that may be responsible for the weaker sensing. To probe the role of FAs, we used endothelial tissues cultured with DMEM (instead of the EC specific medium EGM2 used in controls), which have fewer and smaller FAs (Fig. 3F). These tissues show intact response (Fig. 3F, S3F), but a complete loss of actin fibers circumferential orientation at 7 hours when further treated with EDTA to perturb AJs (Fig. 3G,E). Individual cells still possess ordered stress fibers which shows some remodeling, with the formation of thick bundles (Fig. 3G), consistent with a possible role for the remaining small FAs in tension sensing. Cells are elongated in the direction of their internal stress fibers (Fig. S3D), confirming that cell elongation is an active mechanism driven by actin; but they are not collectively aligned in the tension direction (Fig. 3G,E), consistent with the random orientation of nuclei (Fig. S3E). Taken together these results suggest that AJs are sufficient for tension sensing and actin reorientation, and that FAs can partially rescue the mechanosensing when AJs are perturbed with EDTA.

#### A model for tissue mechanics and actin nematodynamics recapitulates the response of endothelial tubes

To test our hypothesis linking actin dynamics to tissue mechanics, we developed an active surface model to describe the dynamics of the endothelial tube (Fig. 4A). The model is based on a description of actin dynamics encoding mechanosensitive effects observed in experiments, the force balance relating the luminal pressure and the tissue tension, and the relationship between the tissue tension, tissue deformation and actin stress fibers orientation.

**Figure 4.**
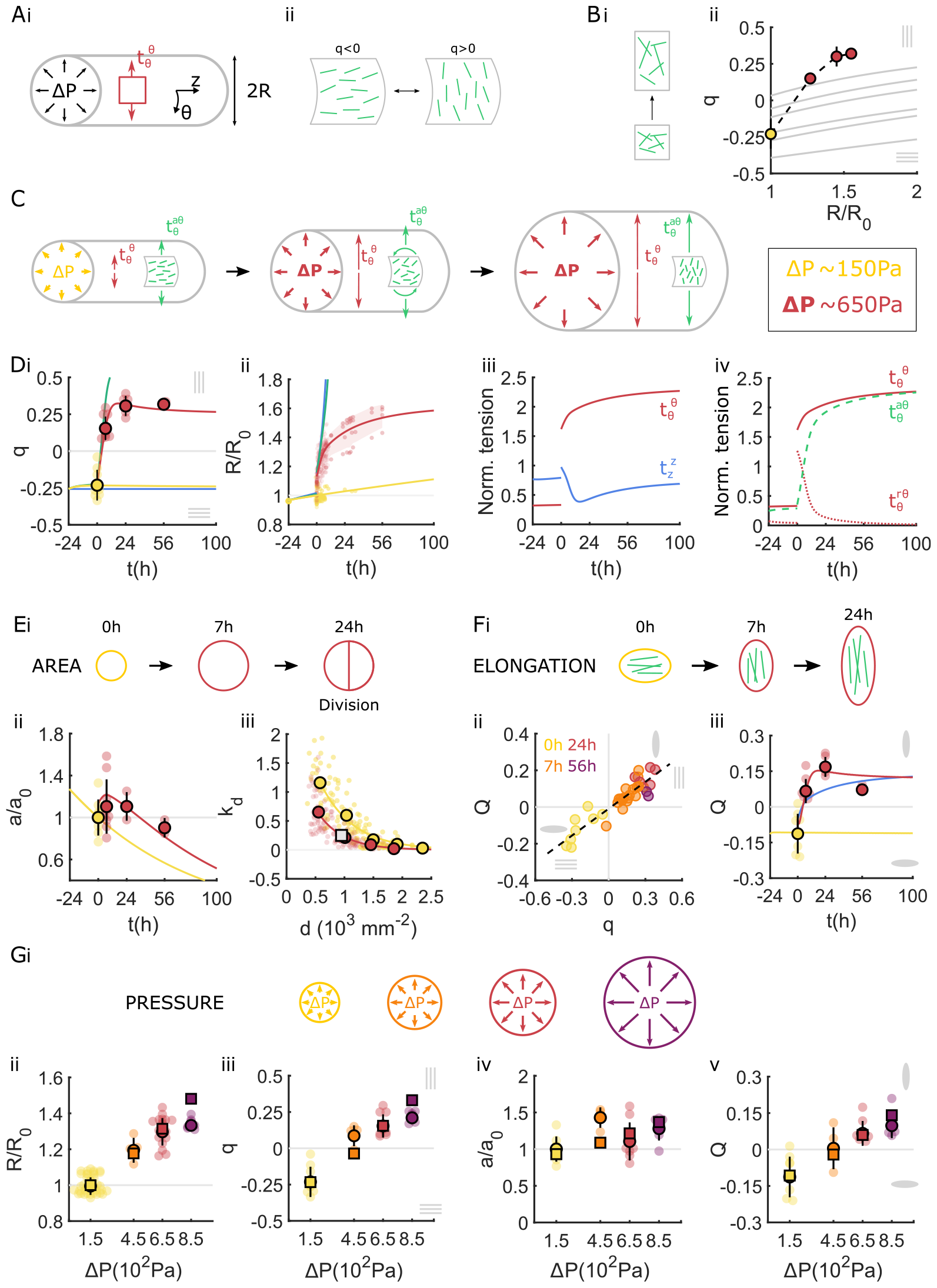
A model for tissue mechanics and actin nematodynamics recapitulates the response of endothelial tubes. **Ai** Schematic of cylindrical tube or radius *R* subjected to the pressure difference Δ*P*, balanced by the circumferential tension 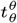. **Aii** The change of orientation of actin fibers from longitudinal to circumferential corresponds to a change of sign of the order parameter *q*. **Bi** Schematic of the change of nematic order through fiber reorientation induced by tissue shear (grey lines in panel ii). **Bii** Circumferential actin nematic order *q*, as a function of the normalized tube radius *R/R*_0_, comparing experimental data (dots) and the numerically computed contribution of deformation by the tissue shear (grey lines). **C** Schematic of tube expansion dynamics and nematic reorientation induced by the tube expansion. A sudden increase in the luminal pressure from Δ*P ≃* 150Pa to Δ*P ≃* 650Pa results in an instantaneous deformation, followed by a reorientation of actin fibers and an increase of the tension generated in actin stress fibers, 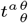, that slows down tube expansion. **D** Actin order parameter *q* (**i**) and normalized tube radius *R/R*_0_ (**ii**) as a function of time, comparing the experimental data (dots) and the model prediction (solid lines), for a constant pressure Δ*P ≃* 150Pa (yellow) and with an pressure increase Δ*P ≃* 650Pa (red). The model prediction without the elastic component of the actin tension (green line, *K*_*a*_ = 0) and without the tension coupling inducing actin reorientation (blue line, *β* = 0) are also shown. **Diii** Normalized total circumferential tension 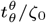 (solid red line), and total longitudinal tension 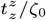 (solid blue line), as a function of time. **Div** Normalized total circumferential tension 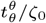 (solid red line), circumferential tension in the actin stress fiber network 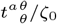 (dashed green line), tissue circumferential tension 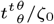 (dotted red line). **Ei** Schematic of the mean cell area dynamics. **Eii** Normalised cell area as a function of time, comparing the experimental data (dots) and the model prediction (solid lines), for a constant pressure Δ*P ≃* 150Pa (yellow) and after the pressure increase Δ*P ≃* 650Pa (red). **Eiii** Proliferation rate *k*_*d*_ as a function of cell density, measured between *t* = 0 and *t* = 7h, for monolayers under low pressure Δ*P*_0_ *≃* 150Pa (yellow dots) and high pressure Δ*P*_*m*_ *≃* 650Pa (red dots). Lines: exponential fit. Grey square: prediction from isotropic shear decomposition. **Fi** Schematic of the cell elongation dynamics. **Fii** Cell circumferential elongation *Q* as a function of the actin nematic order parameter *q*, showing a linear empirical correlation, color coded for time (0h yellow, 7h orange, 24h red and 56h purple). **Fiii** Cell circumferential elongation *Q* as a function of time, comparing the experimental data (dots) and the model prediction (solid lines), for a constant pressure Δ*P ≃* 150Pa (yellow) and with a pressure increase Δ*P ≃* 650Pa (red). Blue line: model prediction for the case where the cell elongation follows tissue deformation. **Gi** Schematic of the different pressures applied to the endothelial tube. **Gii**,**iii** Normalized tube radius *R/R*_0_ (**ii**), actin nematic order parameter *q* (**iii**), normalized cell area *a/a*_0_ (**iv**) and cell elongation *Q* (**v**) as a function of pressure, measured 7 hours after pressure step application, comparing the experimental data (circles) and the model prediction (squares).

#### Actin nematodynamics

We first investigated actin reorientation dynamics during tube expansion. We asked if the reorientation of actin stress fibers was a direct consequence of the anisotropic deformation induced by the circumferential elongation of the tube (Fig. 4B). We therefore tested whether the dependency of the circumferential actin nematic order *q* on the increase of tube radius Δ*R/R* was compatible with a simple geometric reorientation of actin fibers as a response to the tube anisotropic shear (SI). However, this predicted a lower circumferential orientation than observed experimentally (Fig. 4Bii, S4A), indicating that actin stress fibers are not simply following the tube deformation.

We then asked if the dynamics of the actin fibers order *q* could instead be explained by a generic nematodynamics description (Fig. 4C). Upon the application of additional pressure, the order *q* switches from a negative to a positive value but maintains a similar magnitude, suggesting that actin stress fibers maintain a similar level of organisation over time, but reorient strongly. Therefore, we assumed that the actin fibers form a network in the nematic phase, with a spontaneous tendency to order. We then write the following dynamic equation for the actin nematic tensor *q*_*ij*_:

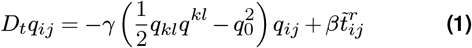

Here, *D*_*t*_ is the corotational time derivative, 1*/γ* a timescale of relaxation of the order parameter *q*_*ij*_, *q*_0_ a magnitude of nematic order, and *β* a mechanosensitive coupling term between the actin order and 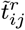 the traceless part of the residual tension 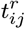, defined as the tension in the tissue that does not come from actin stress fibers (Eqs. 3-5). *β >* 0 describes the tendency of filaments to align along the direction of anisotropic residual tension. Overall, Eq. 1 implies that actin stress fibers tend to reach an ordered state with a preferred strength *q*_0_ in a stress-free configuration, and reorient dynamically according to cues given by the orientation of stresses within the tissue.

#### Active viscoelastic model of tissue tension

Force balance implies that the luminal pressure in the tube Δ*P*, its radius *R* and circumferential tension 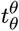 are related by the law of Laplace:

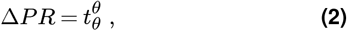

such that at constant pressure and for an increasing tube radius, the circumferential tension must increase. To account for this increase, we assumed that the tissue tension *t*_*ij*_ stems from an elastic tension from the actomyosin network 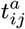, acting along actin stress fibers, and a residual viscoelastic tension from the other tissue components, 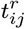. The tension tensor in the tube then follows a rheology combining anisotropic elastic and Maxwell elements:

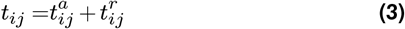

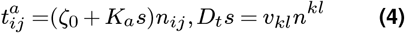

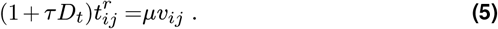

The tension in the actin network is taken to be proportional to the mean orientation tensor *n*_*ij*_ = *g*_*ij*_ */*2 +*q*_*ij*_, with *g*_*ij*_ the surface metric tensor. Our rationale is that the tension in the actin network is generated by a set of actin fibers under tension, with mean orientation *n*_*ij*_ (Fig. 4C). The tension magnitude has a constant active contribution, *ζ*_0_, and an additional contribution proportional to the elongational strain of actin stress fibers *s*. The dynamics of elongational strain depends on the tissue shear *v*_*ij*_, which results in actin stress fiber elongation. *K*_*a*_ is a two dimensional elastic modulus: here actin fibers are assumed to be effectively elastic and to sustain stresses on long time scales when deformed. Laser ablation experiments indicate that the tension is not simply acting along actin stress fibers, as the circumferential tension is larger than the longitudinal tension following pressure application, before actin fibers have reoriented (Figs. 1Bii, S1A, 2C). Therefore we introduce an additional residual tension contribution 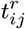 which evolves according to a simple viscoelastic description, with *µ* a viscosity and *τ* a viscoelastic relaxation timescale (Eq. 5). At short time scales, the response of the material is that of a linear elastic material, in line with the linear deformation observed at high collagen concentration in the range of pressure considered here (Fig. 1Ci).

#### Dynamics of tube expansion and fiber reorientation

We found that Eqs. 1-4 account well for the dynamics of the actin nematic order (Fig. 4Di). The increase of circumferential tension after application of additional tube pressure leads to reorientation of actin stress fibers from longitudinal to circumferential on a time scale of *∼* 1 hour. This timescale of reorientation is somewhat larger but comparable in magnitude to previously reported values of actin reorientation in ECs under cyclic stretch Krishnan et al. (2012); Takemasa et al. (1997); Iba and Sumpio (1991); Hayakawa et al. (2001). In line with the model prediction, we did not observe significant deviation of the fiber orientations from the longitudinal or circumferential directions (Fig. S4B, C).

The model also accounts quantitatively well for the dynamics of tube expansion (Fig. 4Dii). In the model, the initial jump in tube radius is limited by the initial elastic response of the tissue (Fig.4C, Diii, S4D). The order of magnitude of fitted values of the elastic coefficients, *K*_*a*_ = 0.22N/m and *K* = *µ/τ* = 0.29N/m, match well with values extracted from pressure ramp application, of the order of 0.26 *−* 0.4N/m (Fig. 1). Elastic residual tension in the tissue then relaxes, leading to further tube expansion (Fig.4C, Dii,iv). The reorientation of fibers however allows limiting and eventually fully opposing expansion of the tube (Fig. 4Di, ii, blue and green curves, S4E). Alternative models we have considered did not account as well for experimental data, except for a model with mechanosensitive coupling of the nematic order to the full tension tensor *t*_*ij*_ (SI, Fig. S5). In the model, setting *β* = 0 (no tension coupling) leads to a lack of circumferential reorientation, such that actin fibers are unable to resist the expansion of the tube (Fig. 4Di, ii, blue curves). Setting *K*_*a*_ = 0 also lead to fast tube expansion, since in that case elastic stresses do not oppose the expansion (Fig. 4Di, ii, green curves). The total tension values (Fig. 4Diii) are anisotropic with a larger circumferential tension after pressure increase, in qualitative agreement with laser ablation experiments (Fig. 1Bii, S1A). The model also predicts a larger longitudinal tension prior to pressure application (Fig. 4Diii), that is detected by laser ablation only in monolayers grown on the soft collagen (Fig. 1Bii, S1A), possibly due to the weak longitudinal anisotropy of the high-density monolayers used for these experiments.

Overall, we propose that actin stress fibers reorient along the direction of highest residual tissue tension, allowing the tissue to resist anisotropic deformation of the tissue (Fig. 4Di-ii).

#### Dynamics of cell area

We then investigated the dynamics of the average cell area in the tissue, which increases transiently after pressure application before decreasing over 56 hours(Fig. 4Eii). The nuclei area mirrors this trend, showing a linear correlation with the cell area maintained throughout the experiment, suggesting that the nuclei are also stretched transiently (Fig. S4F). To understand this dynamics, we note that in the absence of cell apoptosis, the average cell area *a* changes due to tissue expansion and cell division, following an isotropic shear decomposition Etournay et al. (2015); Popović et al. (2017) (Fig. 4Ei):

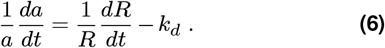

This relationship indeed accounts well for the observed dynamics of the average cell area, for a constant cell division rate *k*_*d*_ = 0.26*d*^*−*1^ (Fig. 4Eii). Upon application of the larger pressure, the average cell area increases due to fast tissue expansion, and it then relaxes due to continued cell division (Fig. 4Ei,ii).

To verify that the corresponding predicted cell division rate value is reasonable, we obtained direct experimental measurements of the cell division rate with EdU labelling measured between 0 and 7 hours at the reference pressure of 150Pa, as well as following application of the larger pressure value of 650Pa (Methods, Fig. 4Eiii). To avoid confounding effects of changes in cell density brought by tissue expansion, we measured cell division rates in endothelial tubes seeded with different initial cell densities. The cell division rate sharply decreases with cell density, with a dependency well fitted by an exponential function (Fig. 4Eiii). One may then expect that increasing tissue tension leads to larger cell area and therefore higher proliferation. Surprisingly however, the cell division rate is lower for larger pressure, implying that increased tissue tension slows down proliferation (Fig. 4Eiii). Therefore, tissue tension influences cell proliferation beyond changing cell area. The predicted cell division rate obtained from isotropic shear analysis (Fig. 4Eii) matches very well with its measured value under high pressure, at the average density of the experiment (Fig. 4Eiii), consistent with cell proliferation being responsible for the decrease in cell area after 7 hours and the absence of apoptosis.

#### Dynamics of cell elongation

We then asked if, similar to the average cell area, the dynamics of cell elongation could be understood with an anisotropic shear decomposition, where the cell elongation changes due to tissue anisotropic shear and cellular rearrangements Etournay et al. (2015); Popović et al. (2017). We postulated that cellular rearrangements are driven by the difference between the actin nematic order and the cell elongation, based on the observed empirical linear correlation between *Q* and *q* (Fig. 4Fi,ii) and on the absence of cell elongation when actin is depolymerized (Fig. 2F), leading to the following dynamics:

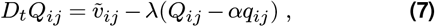

where 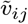.is the traceless part of the tissue shear *v*_*ij*_, 1*/λ* a relaxation timescale of the cell elongation *Q*_*ij*_ to a preferred value *αq*_*ij*_. This hypothesis accounts well for the dynamics of cell elongation (Fig. 4Fiii) for a time scale 1*/λ ∼* 45min, fast compared to the duration of the experiment. Therefore, we propose that cell elongation is largely slaved to the orientation of actin fibers. The final decrease of cell elongation at 56 hours however is not predicted by the theory and could arise from slow effects linked to cell division, which are oriented circumferentially (Fig. 2E), indicating a possible decoupling between the actin stress fibers and cell elongation nematics on longer time scales.

#### Effect of pressure magnitude

To further verify that the proposed physical description indeed accounts for endothelial tube dynamics under pressure, we obtained the predicted tube radius *R*, actin order parameter *q*, cell elongation order *Q* and the cell area *a*, 7 hours after application of a range of applied pressures. The model predicts that the tube radius and reorientation of actin stress fibers and cell elongation axis are strongly dependent on the value of the increasing pressure (Fig. 4G, S4G): for larger pressures, the tube radius expands more and fibers reorient more strongly along the circumferential direction. To test these predictions, we measured the response of the tissue at 7 hours for different imposed pressure differences (Δ*P ≃* 450Pa and Δ*P ≃* 850 Pa). The tube radius, actin orientation and cell orientation at 7 hours all scale roughly linearly with the applied pressure, in good agreement with the theoretical predictions (Fig. 4G, S4G). Overall, we conclude that actin fibers orientation is sensitive to the physiologically relevant range of pressure variations 0 *−* 850 Pa, thanks to the mechanosensitivity of the nematic order of actin stress fibers, encoded by the parameter *β* in Eq. 1.

## Discussion

Here we demonstrate that pressure application on a reconstituted endothelial tube leads to actin stress fiber circumferential alignment and supracellular actin formation. Actin fiber reorientation is parallel to the direction of applied stretch and maximum tension. This response is consistent with alignment of fibers along the direction of maximal tension reported in epithelia *in vivo* López-Gay et al. (2020), but contrasts with the observation that actin fibers in ECs reorient perpendicularly to the direction of stretch under imposed cyclic deformation of the substrate Dessalles et al. (2021a). This shows that the exact nature of the mechanical cue is crucial to set the cell orientation Dessalles et al. (2021a).

Tubes under tension famously tend to be unstable Plateau (1873); de Gennes et al. (2004). Indeed, the law of Laplace (Eq. 2) states that in a pressurised expanding tube, the circumferential tension must increase. The fact that the endothelial tube under pressure appears to resist expansion therefore suggests that it responds elastically. However, the average cell area decreases over long time scales, while the circumferential cell elongation plateaus or decreases (Fig. 4E,F). This implies that the elastic stresses associated with deviation from a preferred value of cell area or cell elongation can not account for the increased tension, in contrast with results in other tissues Etournay et al. (2015). This led us to propose that the long time scale behavior of the endothelial tube is largely driven by actin fibers reorientation and elongational strain. In the model, the actin fibers generate active and elastic stresses along their direction, while a residual tissue tension triggers actin fiber alignment, creating a two-way coupling which allows the system to oppose the applied anisotropic tension. In addition to the actin stress fiber nematic, we have examined simultaneously the dynamics of cell elongation. Although these two nematic fields have been proposed to be decoupled in epithelial layers Nejad et al. (2023), here we find that they follow each other for most of the tube expansion (Fig. 4Fii).

Our microstretcher allows to set a fixed value of pressure magnitude, thus preventing tension relaxation in the tissue, in contrast to length-controlling assays. It mimics a key *in vivo* scenario, modeling pressure increases due to the onset of the heart beat or hypertension, with physiologically relevant pressure differences Lindsey et al. (2014); Fegan et al. (2003); Williams et al. (1990). The setup also allows to study the role played by subcellular components, such as actin and junctions, in the mechanical response of the endothelial tube. As such, the cellular dynamics we report here could underlie vessel remodeling and pathological mechanoadaptation of vessels.

Tension-induced remodeling of adherens junctions is evident by shape change and actin and vinculin association, a mechanism previously linked to the protection of tissue integrity and barrier function Millán et al. (2010); Huveneers et al. (2012); Stoel et al. (2023); Oldenburg and Rooij (2014). *In vivo*, an increase in luminal pressure has been reported to trigger vessel radial growth and junction remodeling along the tension direction in the zebrafish, mirroring our observations Kotini et al. (2022). More broadly, tension anisotropy has been shown to guide many morphogenic events through remodeling of actin cytoskeleton and adhesionsPriya et al. (2020); Shyer et al. (2013); Trubuil et al. (2021); Torres-Sánchez et al. (2021). Application of a controlled pressure in an *in vitro* microstretcher, as performed in this study but with other cell types, would allow to decipher the underlying mechanosensitive couplings. Finally, pressure induced remodeling is believed to underlie other fluid transporting tubular networks, such as the mammary glands, the bronchial tree or the lymphatic network, potentially presenting a mechanism extending beyond the cardiovascular network Torres-Sánchez et al. (2021); Breslin (2014). Thereby, luminal pressure induced wall tension may serve as a universal regulator in vessels in vivo, governing the adaptation of their properties and shape and optimizing them for their specific function. By linking the behavior of biological components to the emergent tissue mechanics, we present a key step towards understanding the process of mechanoadaptation in vessels.

## Methods

### Microvessel-on-chip fabrication

The microvessel-on-chip system consists of a chamber that houses a 120 µm-diameter endothelium-lined channel embedded in a soft collagen hydrogel. The detailed methods for the different components of the system have been described previously Dessalles et al. (2021b) and will be shortly summarized here.

After fabricating the PDMS housing with its inlet and outlet ports, a 120 µm diameter acupuncture needle (Seirin) was introduced into the chamber and the housing was bound to a coverslip through plasma activation. Two PDMS reservoirs were sealed with liquid PDMS to the inlet and outlet ports, their inner diameter matching the diameter of standard plastic straws. The chamber was sterilized with 70% ethanol and 20 min of UV light. To improve collagen adhesion to the PDMS walls, the chamber was coated with 1% polyethylenimine (PEI,an attachment promoter; Sigma-Aldrich) for 10 min followed by 0.1 % glutaraldehyde (GTA, a collagen crosslinker; Polysciences, Inc.) for 20 min. Collagen I was isolated from rat tail tendon as described previously Antoine et al. (2015), to obtain a stock solution of 12 mg mL^−1^. Type I collagen solution was then prepared by diluting the acid collagen solution in a neutralizing buffer at a 1-to-1 ratio, pipetted into the housing chamber, and allowed to polymerize in a tissue culture incubator for 15 min for the baseline 6 mg mL^−1^ collagen concentration and for up to 4 h for lower collagen concentrations. The acupuncture needle was then carefully removed, and the needle holes were sealed with vacuum grease (Bluestar Silicones) to avoid leakage.

### Cell culture, seeding and inhibition

Human umbilical vein ECs (Lonza) were cultured using standard protocols in Endothelial Growth Medium (EGM2; Lonza) and used up to passage 7. Upon confluence, ECs were detached from the flask using trypsin (Gibco, Life Technologies) and concentrated to 10^7^ cells.ml^-1^. 1 µL of the concentrated cell suspension was pipetted through the inlet port of the device. After a 5 min incubation, non-adhering cells were gently flushed out. After 1 h, a flow rate of 2 µL min^−1^ was applied via a syringe pump (PhD Ultra, Harvard apparatus). A confluent monolayer was obtained in 24 hours. For the pharmacological experiments, cells were cultured for one hour prior to the experiment in the presence of 100 nM cytochalasinD in EGM2 for actin disruption, and 5 mM EDTA in EGM2 or 2 mM EDTA in DMEM for cell-cell junction perturbation.

### Monolayer stretch

A hydrostatic pressure head was used to impose both the luminal pressure and the flow rate. A PDMS reservoir was created by puncturing a 5 mm-diameter hole in a PDMS cubic block (1 cm on a side) and gluing the block to the inlet. The reservoir was continuously replenished using a syringe pump at a fixed flow rate. The pressure within the channel is therefore set by the height of the outlet reservoir while the luminal flow rate (and the ensuing pressure gradient) is set by the syringe pump flow rate. The height of medium in the inlet reservoir is equal to that in the outlet reservoir height plus the pressure gradient due to the luminal flow and the channel hydraulic resistance.

During the monolayer growth the flow rate was set to 2 µL min^−1^ and the outlet pressure to 100 Pa, which was maintained for the control channels. A pressure gradient of 100 Pa is established between the inlet and the outlet. For the stretch experiments, at t=0, both inlet and outlet reservoirs were filled with a syringe in less than a minute to impose a hydrostatic pressure at the outlet of 400, 600 or 800 Pa, defined by the height of the reservoir. The outlet pressure was maintained constant for the duration of the experiments (7, 24 or 56 hours), while the inlet pressure was maintained by the flow rate imposed with a syringe pump. The flow rate was increased during the course of the experiments, to 3 µL min^−1^ at t=0h and to 4 µL min^−1^ after 24 hours, to account for the increased diameter and to maintain the pressure gradient roughly constant, 100*P a*. We neglected the effect of the pressure gradient, in both experimental quantifications and theoretical modeling, and considered the average pressure of the channel to be around 150, 450, 650 and 850 Pa for the outlet pressures of 100, 400, 600 and 800 Pa respectively.

### Measurement of the channel deformation

An increase in the luminal pressure leads to a pressure difference across the vessel wall and subsequent channel dilation and circumferential stretch (SV1). The circumferential strain, defined as the ratio of the increase in perimeter of the cross section of the channel to the initial perimeter, was obtained from the vessel diameter. Channel diameters were automatically measured, as previously described Salipante et al. (2022). Briefly, channels edges are detected by identifying the position of the peaks in the intensity gradient along the vertical direction. The diameter is then the mean distance between the two peaks.

### Laser ablation

Laser ablation experiments were performed as described in Boutillon et al. (2021, 2022). The chip was placed on a TriM Scope II microscope (La Vision Biotech) equipped with a femtosecond Mai Tai HP DeepSee laser (Spectra Physics), an Insight DeepSee (Spectra Physics) laser and a XLPLN25XWMP2 (Olympus) 25x water immersion objective. Lifeact mCherry was excited through 2-photon excitation using the Inshight laser set to 1160 nm and ablation was performed using the Mai Tai laser set to 820 nm and exit power at 0.45 mW. Using an electro-optic modulator, the region to be ablated was defined as an XY ROI of 4.5x76 µm located at the level of actin cytoskeleton and oriented either longitudinally or circumferentially. Endothelial cells were imaged with a frame every 130 ms, for 5 time-frames prior to ablation, then ablated for two frames and, finally, imaged for 53 time-frames (SV2). The same channel served for six ablations, three in each directions and alternating, starting from 1.5 mm away from the channel border and separated by 1.5 to 2 mm in order to avoid the influence of one cut on the adjacent cuts. To compute the initial recoil velocity, data were analyzed by manually measuring the distance traveled by the edge of the cut in the first frame relative to its initial position.

### Measurement of the monolayer stiffness

For instantaneous deformations of linear elastic materials, the relationship between tension and strain is dictated by the Young’s modulus. We therefore used the present system to measure the Young’s modulus of the endothelium, as previously described Salipante et al. (2022) (see SI).

#### Measurement of the stress-strain relationship

Briefly, channels were subjected to a one minute long pressure ramp, from 150 Pa to 1000 Pa and the circumferential strain was recorded (SV3). The Young’s modulus is then extracted from the slope of the stress-strain curve, assuming that dissipative stresses are negligible. The pressure was increased at a constant speed by filling simultaneously the inlet and outlet reservoirs with a syringe pump at a 2 mL min^−1^ rate, which corresponds to 1000 Pa min^−1^. Channels were imaged every second in phase contrast with a 10x objective. The increase in diameter was then measured automatically with the same method as for still snapshots (see above). A second method was used, where a similar one minute long pressure ramp was applied, starting at 150 Pa but stopping at 650 Pa. Here there was no continuous imaging, a brightfield image was taken only at the beginning (150Pa) and the end of the pressure application (650Pa), from which the diameters were measured.

#### Fitting procedures for parameter inference

The stress-strain curves of control and pharmacologically-perturbed monolayers on 2 mg mL^−1^ collagen, measured continuously, displayed a typical strain-stiffening behavior (Fig 1C,E). The cell monolayer was then modeled as a 3.6 *±* 0.5µ*m* thick shell, composed of a nonlinear elastic Gent material; while the hydrogel was modeled as a linear elastic material that decreases the pressure drop across the cell layer. The monolayer thickness was measured from fluorescent images of the actin cytoskeleton obtained at the tube mid-plane. By fitting the model to the experimental curves, we inferred the Young’s modulus of the endothelium (see SI).

The stress-strain curves of control monolayers on 6 mg mL^−1^ collagen, measured continuously, displayed a linear relationship (Fig. 1C) up to 900 Pa. The paired-values of the control monolayers on 6 mg mL^−1^ collagen also showed a linear behavior when connected to the reference state *D*_0_ = 125*µm* at Δ*P* = 0 Pa. This reference state was determined with the Gent model (see SI) and consistent with previous measurement of bare channel diameter Dessalles et al. (2021b). We therefore used the value of the radial strain at 650 Pa to estimate the Young’s modulus of these control monolayers (Fig. 1C, F). The paired-values of the pharmacologically-treated monolayers showed a strain-stiffening behavior when connected to the reference state (Fig. 1F), consistent with the diameter at 650 Pa being above the threshold diameter for stiffening, estimated to be around 150 *µ*m (Fig. 1Ei). We therefore used the value of the radial strain at 150 Pa, still in the linear regime, to estimate the Young’s modulus of the cytoD and EDTA treated monolayers (Fig. 1F).

### Immunostaining

Cell-cell junctions were stained using a rabbit anti-VE-cadherin primary antibody (Abcam). Actin filaments and nuclei were stained using Alexa Fluor phalloidin (Invitrogen, Thermo Fisher Scientific) and DAPI (Sigma-Aldrich), respectively. In addition, a mouse anti-vinculin (Abcam) and a mouse anti-vimentin (Abcam) primary antibodies were used to stain for focal adhesions and intermediate filaments. Immunostaining was performed by slow infusion of reagents into the microchannel. Cells were fixed in 4% paraformaldehyde (PFA; Thermo Fisher Scientific) for 15 min, rinsed with phosphatebufered saline (PBS), and then permeabilized with 0.1% Triton in PBS for another 15 min. The channel was then perfused with a 3% bovine serum albumin (BSA) solution in PBS for 1 h to block non-specific binding. Cells were incubated with the primary antibodies (1:400) in PBS for 1 h at room temperature and then rinsed with PBS for an additional 1 h. The channel was then perfused with the secondary antibodies (1:400), phalloidin (1:200), and DAPI (1:1,000,000) in PBS. Finally, the cells were incubated overnight in PBS at 4°C. Samples were imaged using the NIS-Elements software on an epifluorescence inverted microscope (Nikon Eclipse Ti) and/or a Crest X-Light confocal system mounted on an inverted microscope (Nikon Eclipse Ti).

### Analysis of orientation

For statistical analysis of orientations, at least 15 images along the bottom half and the top half of the channel were acquired with a 10x objective. A region of interest was then selected to match the area in focus. Angles were defined relative to the channel longitudinal axis, aligned to the horizontal axis. To be coherent with the tangential basis we chose in the model (Fig. 4Ai), angles were defined positive in the bottom half of trigonometric circle and negative in its bottom half.

#### Actin and cell orientation

Actin fiber orientation was obtained from images of phalloidin stainings using the plugin OrientationJ in ImageJ Rezakhaniha et al. (2011). The window size was set to 5 pixels and the method to cubic spline. The probability distributions of the angles generated by the plugin for each longitudinal position within one channel were then averaged together. The same pipeline was applied to the VE-cadherin immunostaining and to the brightfield images to estimate the cell elongation and orientation.

#### Nuclei orientation

Nuclei were segmented using a custom-made Matlab code. Each nucleus was fitted with an ellipse and the angle of the long axis of the ellipse was used as the nucleus angle. Angles were then binned to create a probability distribution.

#### Division orientation

Mitotic angles were manually measured from nuclear (DAPI) immunostainings. Dividing cells were identified by their condensed chromosomes and the angle between the segment connecting the two daughter nuclei and the longitudinal axis was measured using ImageJ. Angles were grouped and then binned to create a probability distribution.

#### Nematic order parameter calculation

The tensors *q*_*ij*_, *Q*_*ij*_ and *Q*_*nij*_ are defined as:

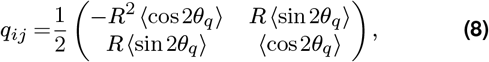

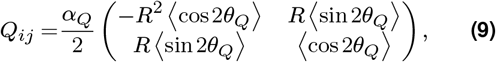

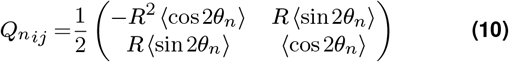

where *θ*_*q*_, *θ*_*Q*_ and *θ*_*n*_ are a set of angles obtained by the image analysis, as described above. In the case of symmetric distribution of angles *θ*_*q*_, *θ*_*Q*_ and *θ*_*n*_around 0, off-diagonal terms of the tensors *q*_*ij*_, *Q*_*ij*_ and *Q*_*nij*_ vanish and actin orientation, cell elongation and nucleus elongation can be described by the order parameters 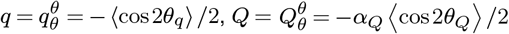 and 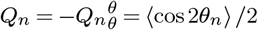. We chose the convention where a positive value of *q, Q* or *Q*_*n*_ indicates that the orientation is preferentially circumferential. Conversely a negative value of *q, Q* or *Q*_*n*_ indicates a preferentially longitudinal orientation. The magnitude of *q, Q* or *Q*_*n*_ corresponds to the strength of the alignment along the circumferential or longitudinal direction.

#### Calculation of the α_Q_

Because the orientation of junctions does not necessarily exactly reflect cell elongation, we introduce a correction factor *α*_*Q*_ as follows. For a uniform shear flow 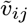 (traceless part of the gradient of flow *v*_*ij*_) and in the absence of cellular rearrangements, we expect 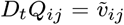. In the context of a cylindrical tube, this implies that the variation of cell elongation following a change of radius and for a homogeneous material deformation should be 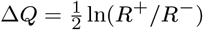, with *R*^*−*^ (resp. *R*^+^) the radius of the tube before (resp. after) the deformation. We tested this relation by imposing a fast change of pressure on tubes of different radii, and by measuring the resulting change of tube radius and change in measured average cell elongation Δ*Q*. For measurement of Δ*Q* with a fast change of pressure the junction angle distribution was obtained from brightfield images, instead of VE-cadherin stainings in the general case. Using images with both VE-cadherin staining and brightfield pictures, we find that the measured value of *Q* using VE-cadherin staining corresponds to approximately half its measured value using brightfield picture (Fig. S6A). Converting the value of *Q* compute from brightfield to its VE-cadherin staining based value, we finally found that the linear relation 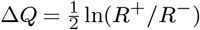 was satisfied for a correction factor *α*_*Q*_ = 0.8 (Fig. S6B).

### Nuclei metrics and density measurements

Nuclei were segmented using the StarDist ImageJ pluggin Schmidt et al. (2018). Nuclei area and aspect ratio were extracted from each segmented object in ImageJ. Cell density was calculated by dividing the number of cells in a field of view by the area (accounting for channel curvature), for each position along the channel length. The mean cell area was calculated as the inverse of cell density.

### Proliferation assay

To assess EC proliferation, EdU was added to the cell culture medium at a concentration of 10 µM. Cells were maintained in EdU-containing culture medium for 8h, either at 150Pa or at 650Pa, after which they were fixed and stained for DAPI and EdU-positive nuclei. The fraction of EdU-positive to EdU-negative nuclei provides a measure of the proliferation rate.

### Statistical analysis

For Figures 1-4, all data are plotted as mean ± standard deviation, except for the probability distribution plots where the shadowed area corresponds to the standard error of the mean. An unpaired Student t-test was used for significance testing between two conditions. Statistical tests were performed using Matlab. *** denotes p < 0.001, ** denotes p < 0.01, and * denotes p < 0.05.

## Acknowledgments

We thank Pierre Mahou and the Polytechnique Bioimaging Facility for assistance with live imaging on their equipment partly supported by Région Ile-de-France (interDIM) and Agence Nationale de la Recherche (ANR-11-EQPX-0029 Morphoscope2, ANR-10-INBS-04 France BioImaging). We also thank Michael Riedl for constructive inputs on the manuscript. This work was funded in part by an endowment in Cardiovascular Bioengineering from the AXA Research Fund to AIB, an AMX doctoral fellowship from Ecole Polytechnique and an EMBO fellowship ALT 886-2022 to CAD. NC was supported by a SNSF project grant 200021_197068 to GS.

## Supplementary Information

**Figure S1.**
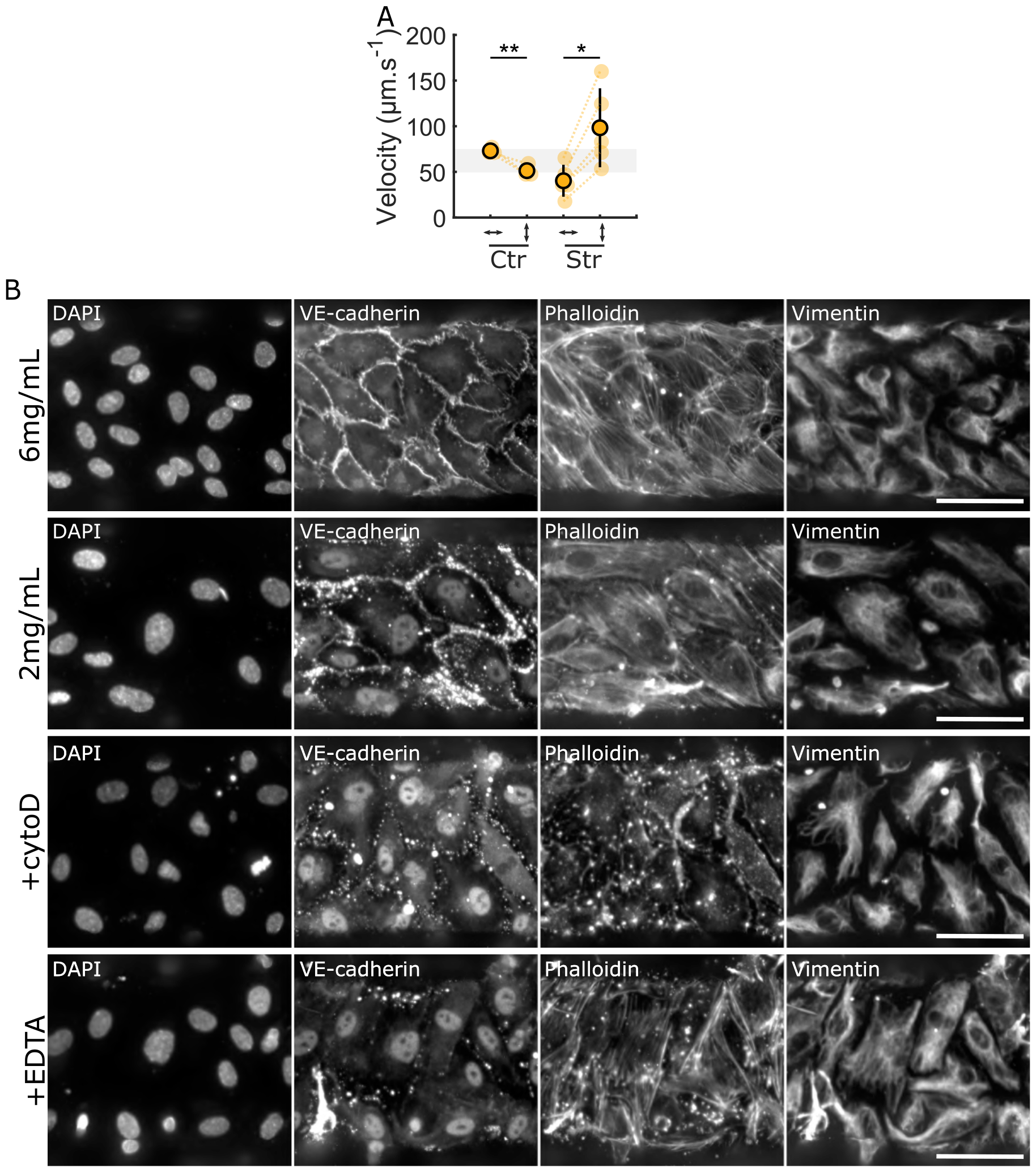
Endothelial tubes exhibit actin-dependent elasticity under luminal pressure. **A** Initial recoil velocity post ablation for monolayers cultured on a 2 mg mL^−1^ collagen gel, showing an increase between the control (150Pa) and stretched (650Pa) channels, but only in the circumferential direction. **B** Endothelium stained for DAPI, VE-cadherin, phalloidin and vimentin one minute after the application of the 1000 Pa pressure, for the two collagen concentrations: 2 mg mL^−1^ and 6 mg mL^−1^, and with the cytochalasinD or EDTA treatment, on the soft gel. Scale bar 50 µm.

**Figure S2.**
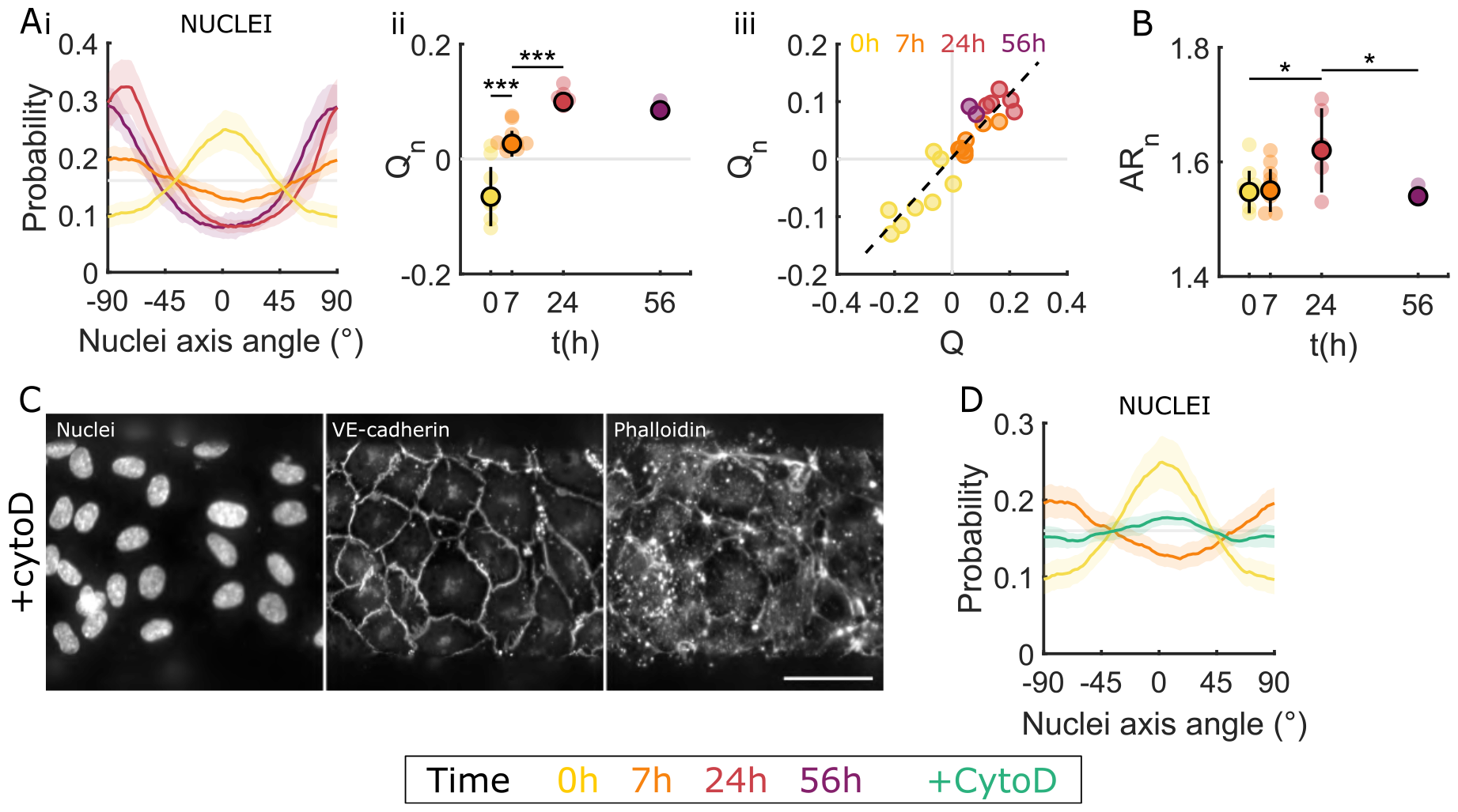
Cells dynamically align in the tension direction via an active actin-dependent process. **A** Evolution of the probability distribution of the nuclei orientation (**i**) and the associated nematic order parameter *Qn* (**ii**) at 0 (yellow), 7 (orange), 24 (red) and 56 (purple) hours. The sign of *Qn* denotes the direction of the orientation and the absolute value its strength. **Aii** *Qn* as a function of the cell elongation order parameter *Q*, showing a linear empirical correlation, color coded for time (0h yellow, 7h orange, 24h red and 56h purple). **B** Mean nucleus aspect ratio as a function of time. **C** Cytochalasin-treated monolayer stained for nuclei, VE-cadherin and phalloidin after 7h of pressure showing round cells, the presence of cell-cell junctions and the absence of actin fibers. Scale bar 50 µm. **D** Evolution of the probability distribution of the nuclei orientation before the pressure increase (Δ*P ≃* 150Pa, yellow), and after 7h of high pressure for the control (Δ*P ≃* 650Pa, orange) and cytochalasin-treated (green) monolayers.

**Figure S3.**
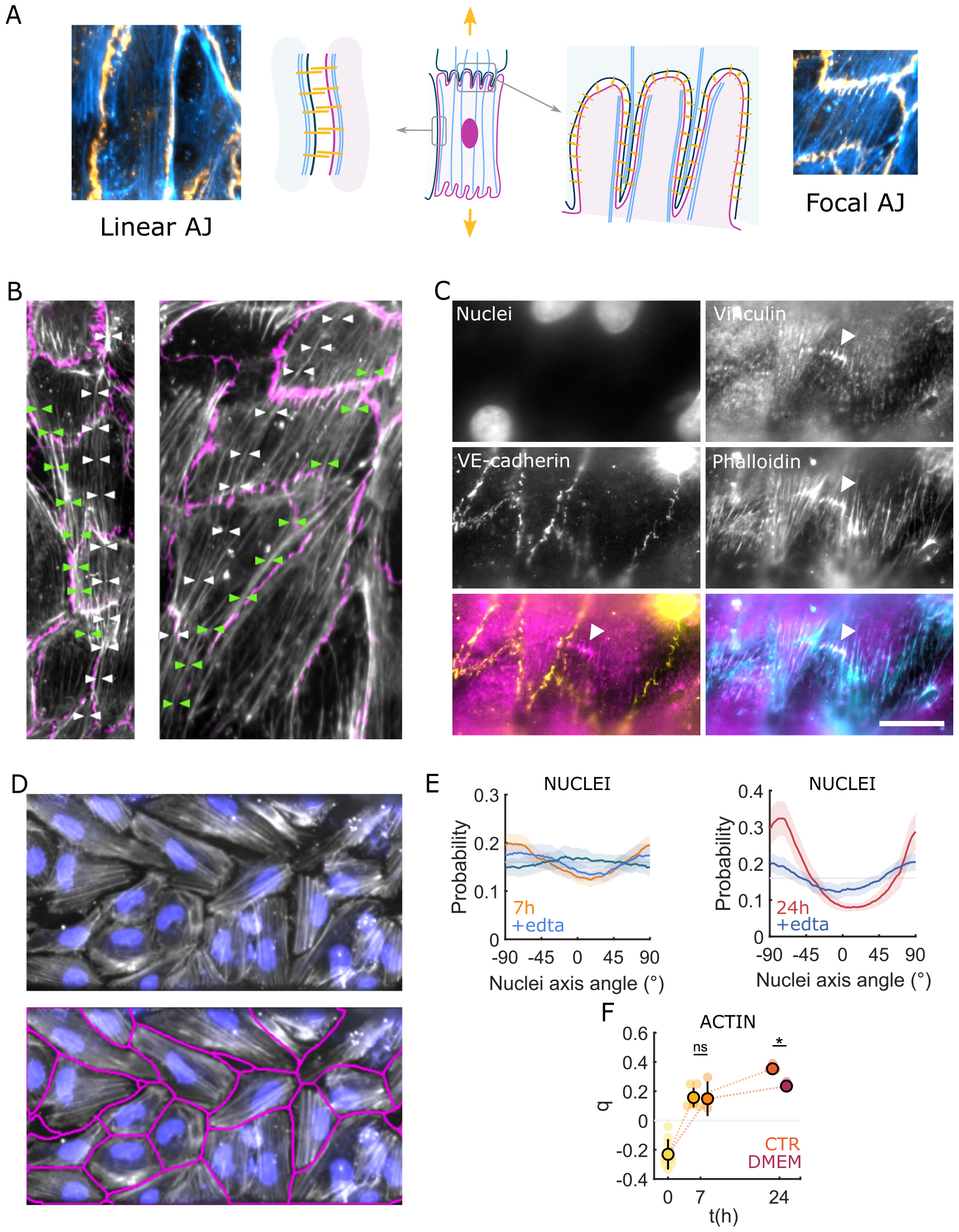
Cell-cell junctions and focal adhesions are necessary for actin alignment. **A** Schematics of the cell-cell junctions and associated actin fibers, showing a linear AJ (left) and a focal AJ (right), observed in monolayers under Δ*P ≃* 150Pa. **B** Endothelium stained for VE-cadherin (pink) and phalloidin (white) after 7h of stretch at Δ*P ≃* 650Pa, with long transcellular actin cables manually traced (white and green triangles). **C** Endothelium stained for nuclei, vinculin, VE-cadherin and phalloidin after 7h of stretch at Δ*P ≃* 650Pa, showing a line of clustered focal adhesion with actin fibers anchoring (white arrowhead), away from cell-cell junctions (yellow). Scale bar 20 µm. **D** EDTA-treated endothelium stained for nuclei (blue) and phalloidin (white) after 7h of stretch at Δ*P ≃* 650Pa, with cell border automated detection (red, bottom), showing elongated cells whose orientation follows stress fiber orientation. **E** Probability distribution of the nuclei long axis at 7 hours (**i**) and 24 hours (**ii**) for control (orange-red) and EDTA-treated (blue) endothelia. **F** Nematic order parameter q as a function of time for EGM2-cultured (light orange) and DMEM-cultured monolayers (dark orange).

**Figure S4.**
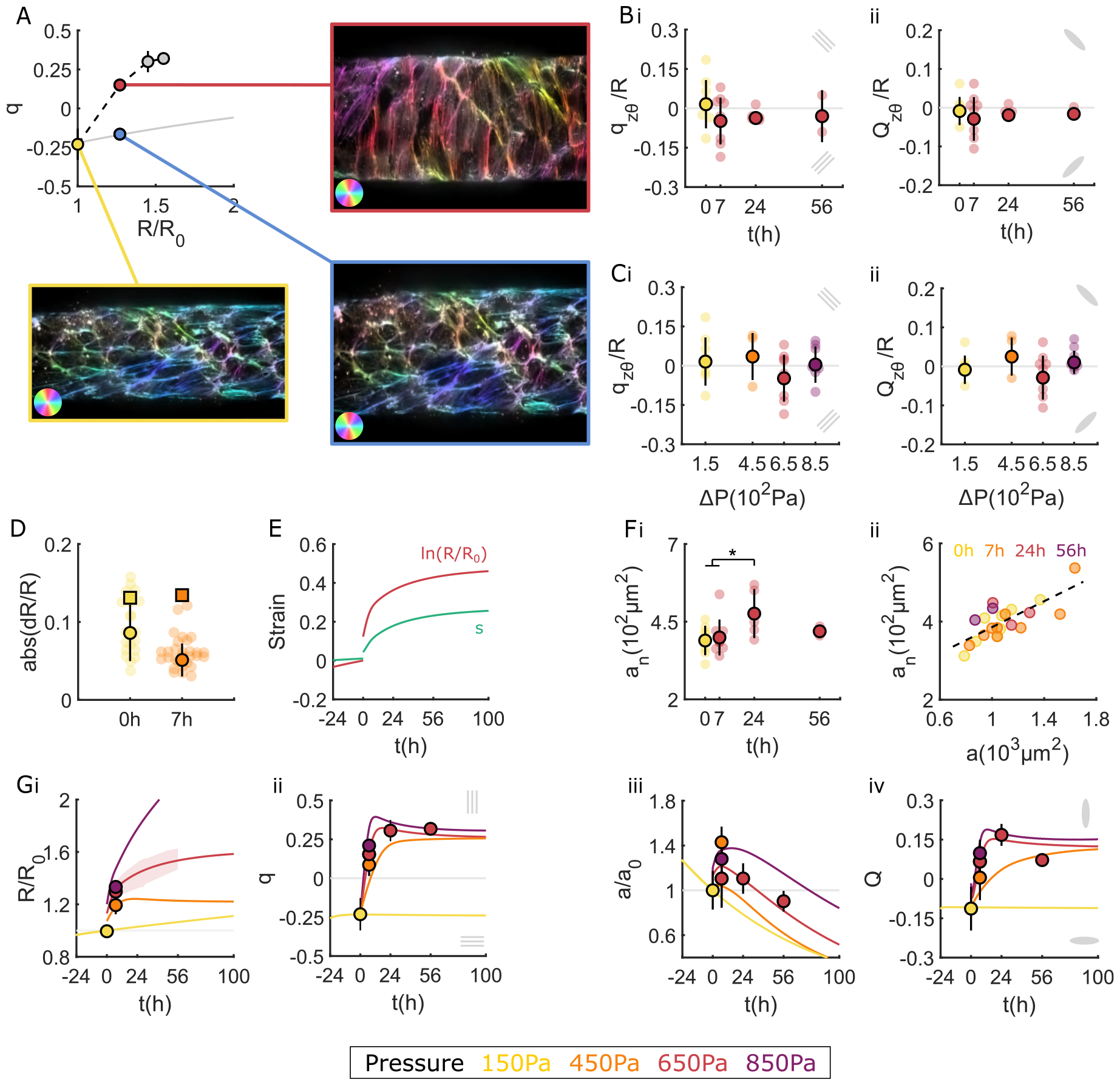
A model for tissue mechanics and nematodynamics recapitulates the response of endothelial tubes. **A** Circumferential actin nematic order *q*, as a function of the normalized tube radius *R/R*_0_, comparing experimental data (dots) and the numerically computed contribution of deformation by the tissue shear, starting with a similar value for the reference state *R/R*_0_ = 1(grey line). Insets show actin fibers color-coded for their orientation, before the stretch (yellow), after 7 hours of 650 Pa (red) and for an artificial deformation of the initial image by an amount corresponding to the *R/R*_0_ at 7 hours (blue). The reorientation due to the shear (blue) is visibly smaller than the reorientation observed in experiments for the same deformation (red). **B**,**C** Value of normalized off-diagonal component *q*_*zθ*_ */R* of actin orientation tensor (**Bi**,**Ci**) and *Q*_*zθ*_ */R* of cell elongation tensor (**Bii**,**Cii**) at different time after pressure increase from 150Pa (yellow) to 650Pa (red) (**B**) and 7 hours after different pressure increases (**C**). **D** Absolute relative radius change when increasing pressure from 150Pa to 650Pa (yellow) and when decreasing pressure back to 150Pa 7 hours later (orange), comparing the experimental data (dots) to the theoretical prediction (squares, SI, section 2.5). **E** Tissue strain ln(*R/R*_0_) (red) and actin strain *s* (green) as a function of time, in the model. **Fi** Mean nucleus area as a function of time (experimental data). **Fii** Nucleus area as a function of cell area after 0 (yellow), 7 (orange), 24 (red) and 56 (purple) hours, showing a linear correlation. **G** Time evolution of normalized radius *R/R*_0_ (**i**), actin order parameter *q* (**ii**), mean cell area (**iii**) and cell elongation (**iv**) for Δ*P* = 150Pa (yellow), 450Pa (orange), 650Pa (red) and 850Pa (violet) (lines: model, dots: experiment)

**Figure S5.**
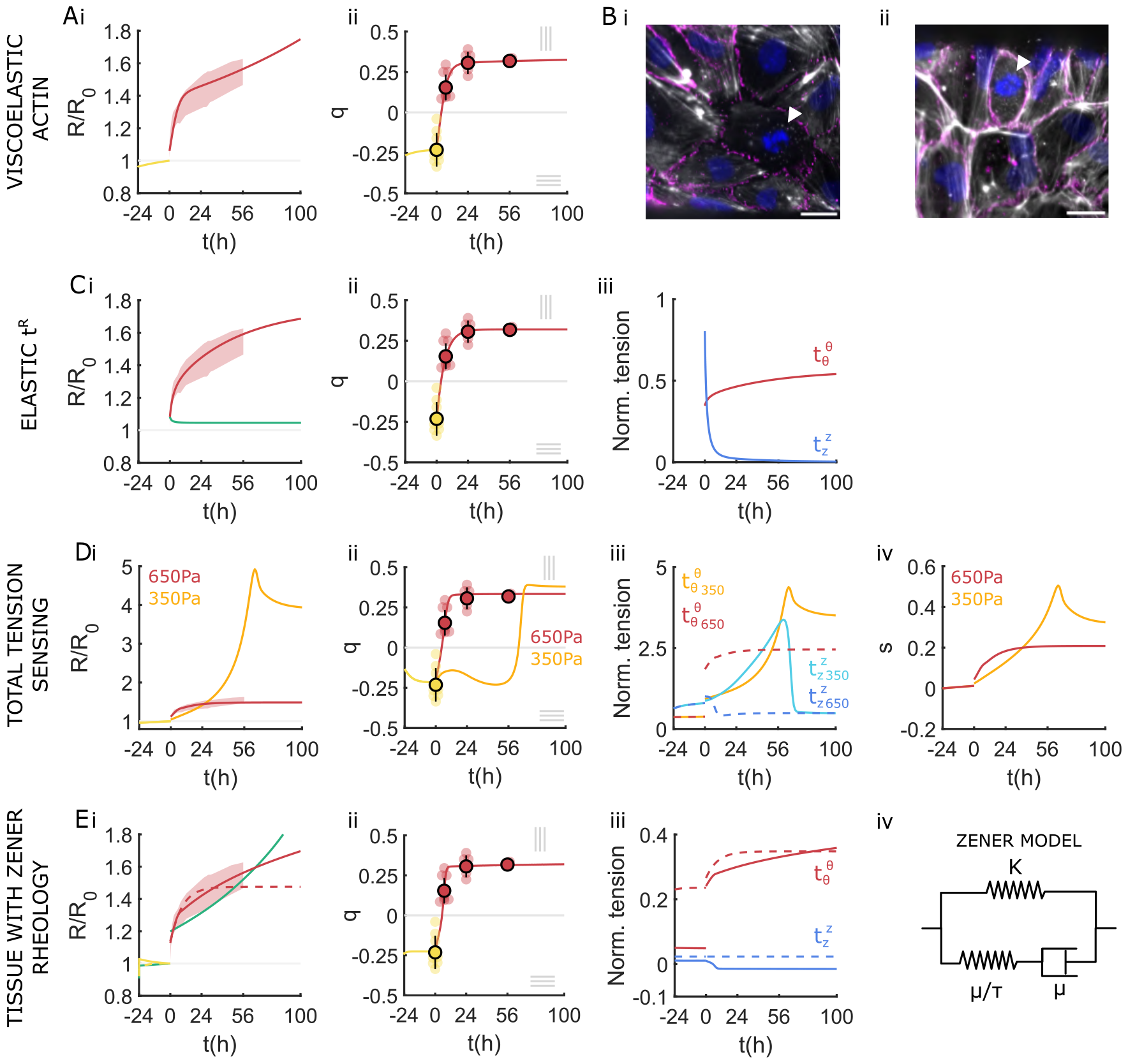
Exploring alternative models of endothelial tube mechanics. **A** Time evolution of normalized radius *R/R*_0_ (**i**) and actin order parameter *q* (**ii**), in a model with viscoelastic actin accounting for actin strain memory loss (solid line, SI Sec. 3.2) compared to experimental data (shadow, dots). **B** Endothelium stained for DAPI (blue), VE-cadherin (magenta) and phalloidin (white) under low pressure (**i**) and high pressure (**ii**), showing a mitotic cell (arrowhead) with condensed chromatin and no stress fibers. Scale bar 20 µm. **C** Time evolution of normalized radius *R/R*_0_ (**i**) obtained for a model where the total tension is the sum of a purely elastic tension 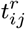 and a tension 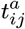 oriented along actin stress fibers, either purely elastic (green line, SI Sec. 3.3) or viscoelastic (red line, SI Sec. 3.3), fitted to experimental data. Red shadow indicates experimental data. Here the actin stress fibers order parameter has been fitted by *q*(*t*) = *−*0.25 + 0.57 *×* (1 *−* exp(*−*0.17*t*)) (**ii**, dots show experimental data). **Ciii** Time evolution of normalized circumferential tension 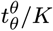 (red) and longitudinal tension 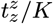 (blue), computed from the model in (i), with viscoelastic actin oriented tension. **D** Time evolution of normalized radius *R/R*_0_ (**i**) and actin order parameter *q* (**ii**) after a pressure increase from 150Pa to 350Pa (orange) and 650Pa (red), obtained from a model where actin fibers mean orientation dynamics is coupled to total tension (solid lines, SI Sec. 3.4), fitted to experimental data. Shadow and dots indicate experimental data. **Diii** Time evolution of normalized circumferential tension 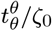 (red and orange) and longitudinal tension 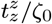 (blue and cyan) for a pressure increase from 150Pa to 350Pa (red and blue) and 650Pa (orange and cyan), computed from the model in (i). **Div** Time evolution of actin strain *s* after a pressure increase from 150Pa to 350Pa (orange) and 650Pa (red), computed from this model. **Ei** Time evolution of normalized radius *R/R*_0_ obtained from the following models: tissue described as an isotropic material following a Zener rheology, without (green, SI Sec. 3.6), or with isotropic active tension (red dashed line, SI Sec. 3.6), and tissue described as an elastic material with Zener rheology, together with an active tension contribution oriented along actin stress fibers (red solid line, SI Sec. 3.7). Shadow indicates experimental data. **Eii** Time evolution of actin order parameter *q* obtained from the model describing the tissue as an elastic material with Zener rheology together with actin oriented active tension (red), fitted to experimental data. Dots indicate experimental data (dots). **Eiii** Time evolution of normalized circumferential tension 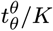 (red) and longitudinal tension 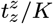 (blue), computed from either a model describing the tissue as an elastic material with Zener rheology and subjected to actin oriented active tension (solid line) or a model describing the tissue as an isotropic material following a Zener rheology, together with isotropic active tension (dashed lines). **Eiv** Schematics of the Zener rheology, consisting of a spring of elastic modulus *K*, in parallel with a serial association of a dashpot of viscosity *µ* and a spring of elastic modulus *µ/τ*, with *τ* a characteristic time.

**Figure S6.**
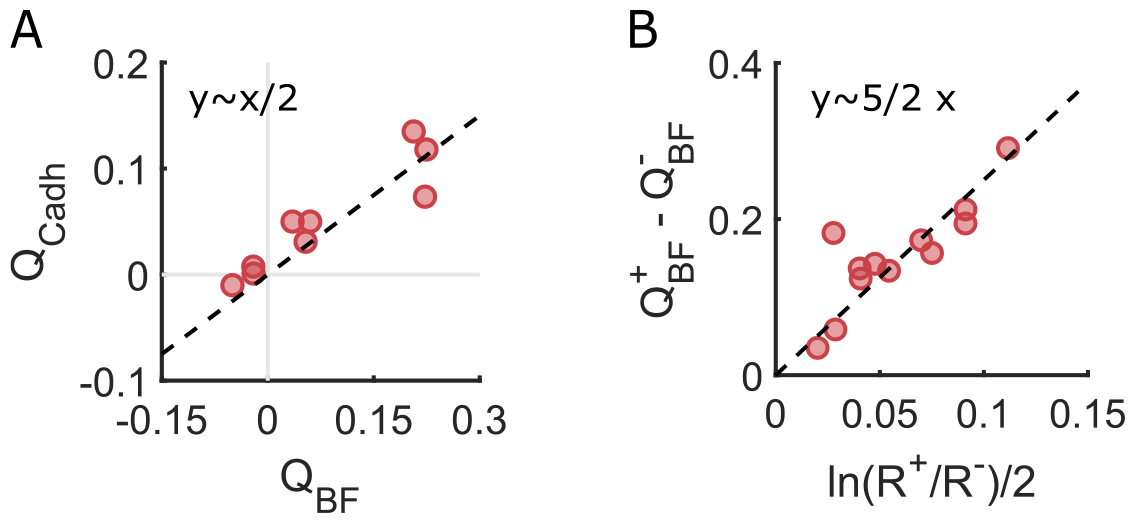
Determination of the coefficient in the cell elongation nematic parameter Q. **A** Cell order parameter extracted from immunostaining of VE-cadherin *Q*_*Cadh*_ as a function of the cell order parameter extracted from phase contrast brightfield *Q*_*BF*_ showing a linear correlation with a 0.5 coefficient. **B** Jump in the cell order parameter, extracted from phase contrast brightfield *Q*_*BF*_ as a function of the logarithm of the jump in radius, showing a linear correlation with a 5/2 coefficient. 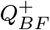 and 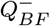 (resp. *R*^+^ and *R*^*−*^) are respectively the value of *Q*_*BF*_ (resp. *R*) just before and just after the pressure increase.

## Supplementary Videos

**Figure SV1.**
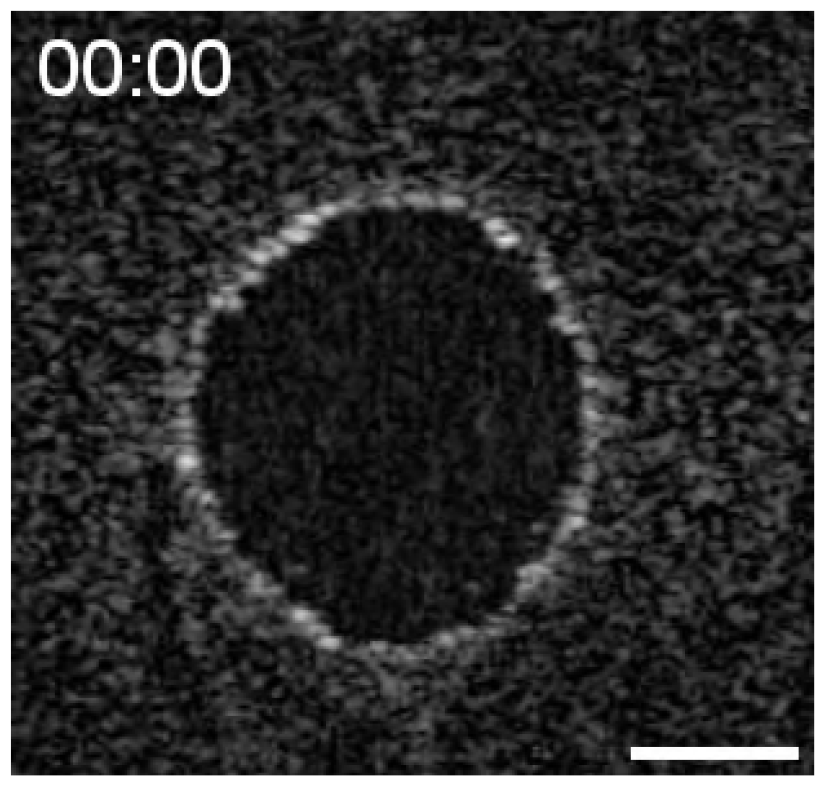
Timelapse of the endothelial tube pressurization. Timelapse of the channel cross-section under a rapid pressure increase from 150Pa to 650Pa, showing a elastic expansion followed by a stable radius. Time, seconds. Scale bar, 50 µm.

**Figure SV2.**
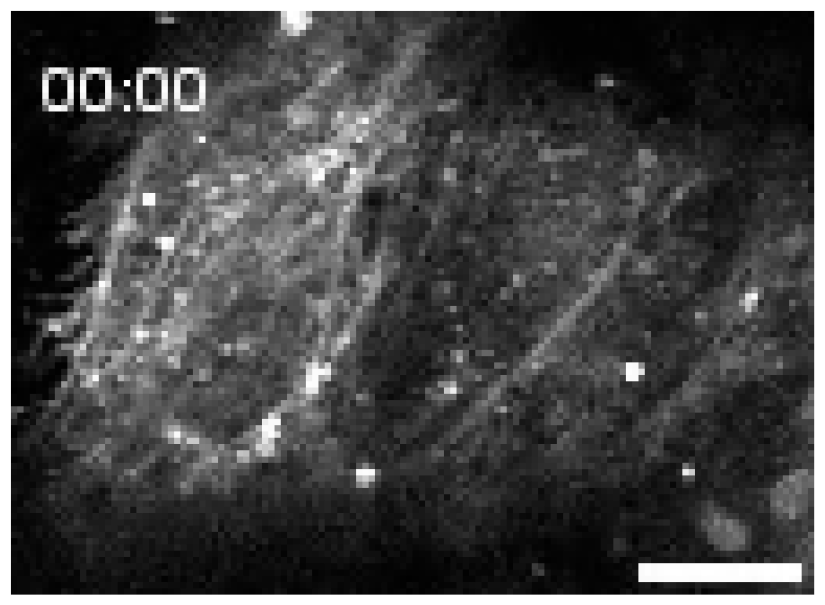
Timelapse of a laser ablation experiment. Fluorescence timelapse of LifeAct-ECs on a soft collagen gel showing the endothelial actin network prior and post longitudinal ablation, with a rapid opening of the wound, characteristic of high tissue tension in the circumferential direction. Time, seconds. Scale bar, 20 µm.

**Figure SV3.**
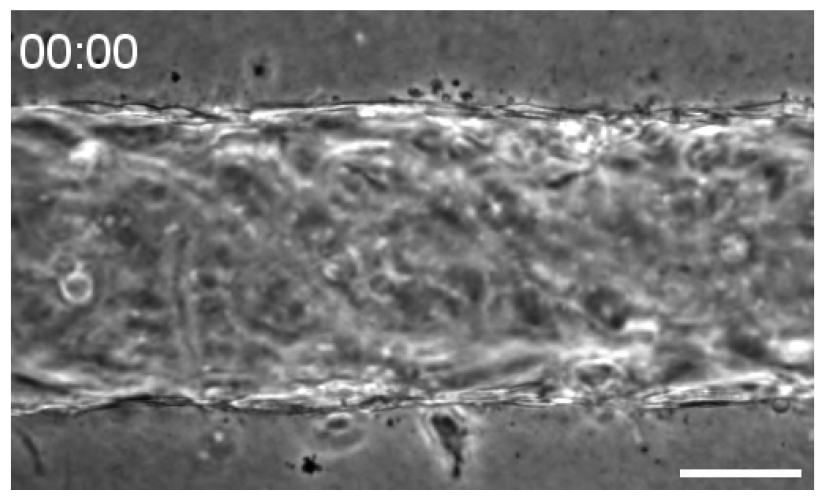
Timelapse used for monolayer stiffness measurement. Brightfield timelapse of an endothelial tube on a soft gel subjected to a linear increase in pressure from 150Pa to 1000Pa, undergoing circumferential expansion with a visible slow down caracteristic of a strain-stiffening behavior. Time, seconds. Scale bar, 50 µm.

**Figure SV4.**
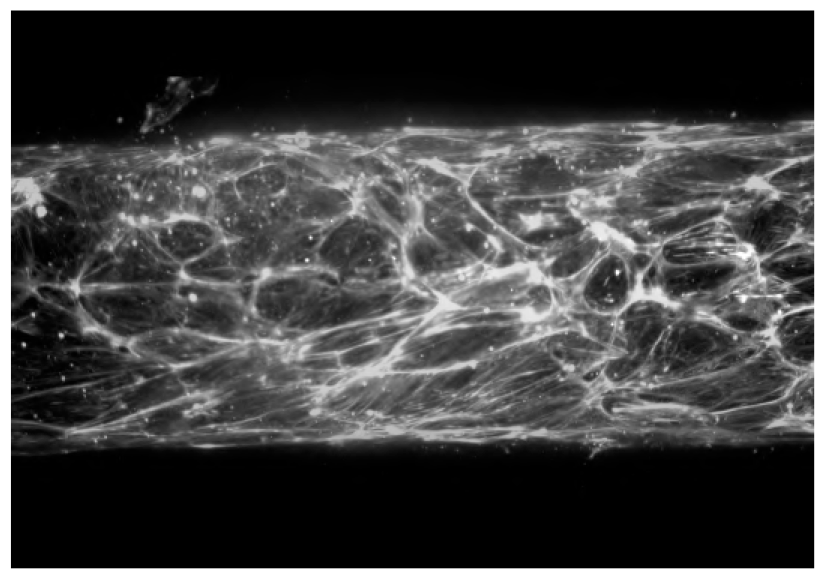
3D reconstruction of the actin network of an endothelial tube at 150Pa. Obtained from a Z-stack of fluorescent images of a monolayer stained with phalloidin, showing longitudinal alignment of the actin stress fibers.

**Figure SV5.**
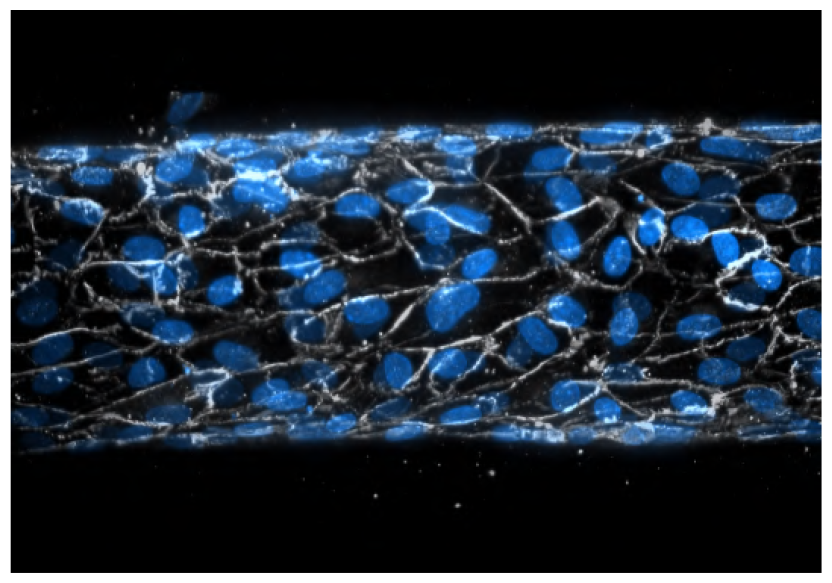
3D reconstruction of the cells and nuclei of an endothelial tube at 150Pa. Obtained from a Z-stack of fluorescent images of a monolayer stained with VE-cadherin (white) and DAPI (cyan), 7 hours after the pressure increase, showing longitudinal alignment of the cells and nuclei.

**Figure SV6.**
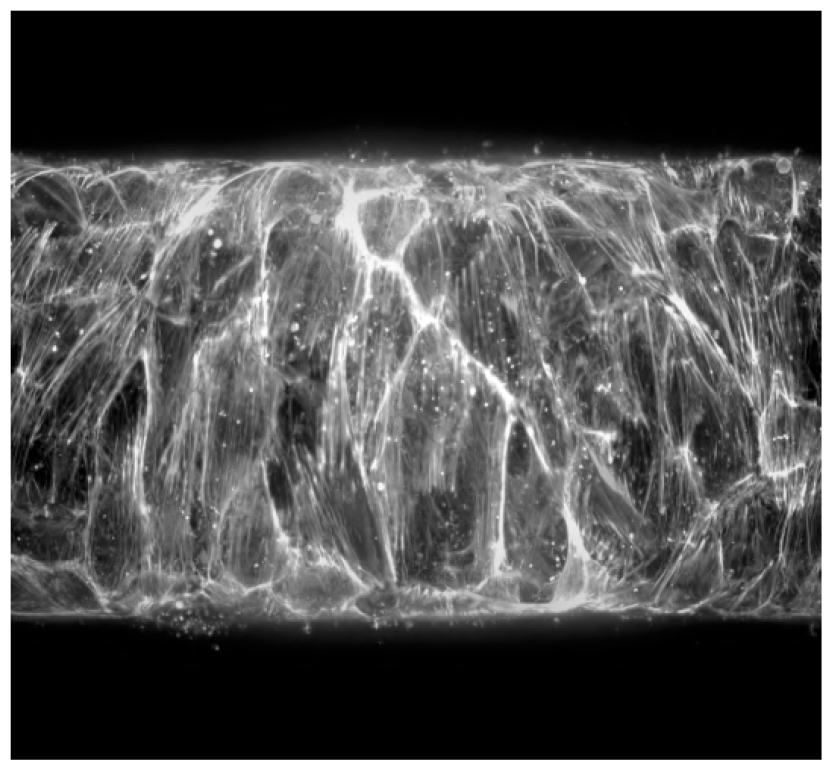
3D reconstruction of the actin network of an endothelial tube at 650Pa. Obtained from a Z-stack of fluorescent images of a monolayer stained with phalloidin, 7 hours after the pressure increase, showing circumferential alignment of the actin stress fibers.

**Figure SV7.**
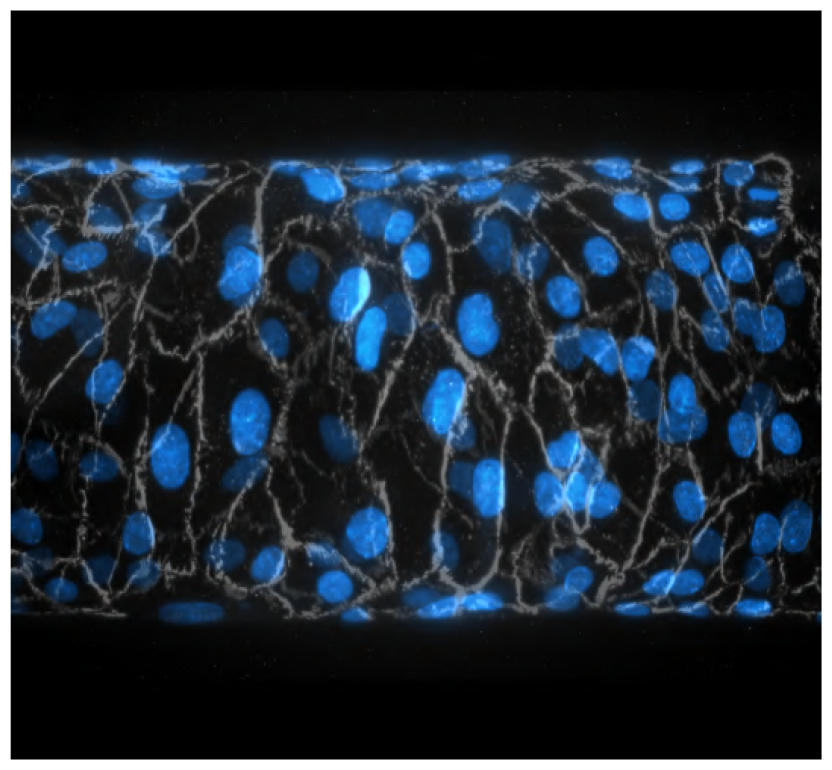
3D reconstruction of the cells and nuclei of an endothelial tube at 650Pa. Obtained from a Z-stack of fluorescent images of a monolayer stained with VE-cadherin (white) and DAPI (cyan), showing circumferential alignment of the cells and nuclei.

## Bibliography

Alert, R. and Trepat, X. (2020). Physical models of collective cell migration. 10.1146/annurev-conmatphys-031218-013516, 11:77–101.

Antoine, E. E., Vlachos, P. P., and Rylander, M. N. (2015). Tunable collagen i hydrogels for engineered physiological tissue micro-environments. PLOS ONE, 10(3).

Bonakdar, N., Gerum, R., Kuhn, M., Spörrer, M., Lippert, A., Schneider, W., Aifantis, K. E., and Fabry, B. (2016). Mechanical plasticity of cells. Nature Materials, 15(10):1090–1094.

Bonnet, I., Marcq, P., Bosveld, F., Fetler, L., Bellaïche, Y., and Graner, F. (2012). Mechanical state, material properties and continuous description of an epithelial tissue. Journal of The Royal Society Interface, 9:2614–2623.

Boutillon, A., Escot, S., and David, N. B. (2021). Deep and spatially controlled volume ablations using a two-photon microscope in the zebrafish gastrula. Journal of Visualized Experiments, (173).

Boutillon, A., Escot, S., Elouin, A., Jahn, D., González-Tirado, S., Starruß, J., Brusch, L., and David, N. B. (2022). Guidance by followers ensures long-range coordination of cell migration through α-catenin mechanoperception. Developmental Cell, 57(12).

Breslin, J. W. (2014). Mechanical forces and lymphatic transport. Microvascular research, 96:46–54.

Campas, O., Noordstra, I., and Yap, A. S. (2023). Adherens junctions as molecular regulators of emergent tissue mechanics. Nature Reviews Molecular Cell Biology 2023, pages 1–18.

Charras, G. and Yap, A. S. (2018). Tensile forces and mechanotransduction at cell–cell junctions. Current Biology, 28(8).

Constantinou, I. and Bastounis, E. E. (2023). Cell-stretching devices: advances and challenges in biomedical research and live-cell imaging. Trends in Biotechnology, 41:939–950.

Czirok, A., Varga, K., Elod, Mehes, and Szabo, A. (2013). Collective cell streams in epithelial monolayers depend on cell adhesion. New Journal of Physics, 15:075006.

Davis, J. R., Ainslie, A. P., Williamson, J. J., Ferreira, A., Torres-Sánchez, A., Hoppe, A., Mangione, F., Smith, M. B., Martin-Blanco, E., Salbreux, G., and Tapon, N. (2022). Ecm degradation in the drosophila abdominal epidermis initiates tissue growth that ceases with rapid cell-cycle exit. Current Biology, 32:1285–1300.e4.

de Gennes, P.-G., Brochard-Wyart, F., and Quéré, D. (2004). Capillarity and wetting phenomena. Capillarity and Wetting Phenomena.

Dessalles, C. A., Leclech, C., Castagnino, A., and Barakat, A. I. (2021a). Integration of substrate- and flow-derived stresses in endothelial cell mechanobiology. Communications Biology, 4(1).

Dessalles, C. A., Ramon-Lozano, C., Babataheri, A., and Barakat, A. I. (2021b). Luminal flow actuation generates coupled shear and strain in a microvessel-on-chip. Biofabrication, 14(1):p015003.

Efremov, Y. M., Zurina, I. M., Presniakova, V. S., Kosheleva, N. V., Butnaru, D. V., Svistunov, A. A., Rochev, Y. A., and Timashev, P. S. (2021). Mechanical properties of cell sheets and spheroids: the link between single cells and complex tissues. Biophysical Reviews 2021 13:4, 13:541–561.

Etournay, R., Popović, M., Merkel, M., Nandi, A., Blasse, C., Aigouy, B., Brandl, H., Myers, G., Salbreux, G., Jülicher, F., et al. (2015). Interplay of cell dynamics and epithelial tension during morphogenesis of the drosophila pupal wing. Elife, 4:e07090.

Fegan, P. G., Tooke, J. E., Gooding, K. M., Tullett, J. M., MacLeod, K. M., and Shore, A. C. (2003). Capillary pressure in subjects with type 2 diabetes and hypertension and the effect of antihypertensive therapy. Hypertension, 41:1111–1117.

Hallatschek, O., Datta, S. S., Drescher, K., Dunkel, J., Elgeti, J., Waclaw, B., and Wingreen, N. S. (2023). Proliferating active matter. Nature Reviews Physics, 5(7):407–419.

Han, M. K. and de Rooij, J. (2016). Converging and unique mechanisms of mechanotransduction at adhesion sites. Trends in Cell Biology, 26(8):612–623.

Harris, A. R., Daeden, A., and Charras, G. T. (2014). Formation of adherens junctions leads to the emergence of a tissue-level tension in epithelial monolayers. Journal of Cell Science, 127:2507–2517.

Harris, A. R., Peter, L., Bellis, J., Baum, B., Kabla, A. J., and Charras, G. T. (2012). Characterizing the mechanics of cultured cell monolayers. Proceedings of the National Academy of Sciences, 109(41):16449–16454.

Hatami, J., Tafazzoli-Shadpour, M., Haghighipour, N., Shokrgozar, M. A., and Janmaleki, M. (2013). Influence of cyclic stretch on mechanical properties of endothelial cells. Experimental Mechanics, 53:1291–1298.

Hayakawa, K., Sato, N., and Obinata, T. (2001). Dynamic reorientation of cultured cells and stress fibers under mechanical stress from periodic stretching. Experimental Cell Research, 268:104–114.

Hoefer, I. E., Adel, B. D., and Daemen, M. J. (2013). Biomechanical factors as triggers of vascular growth. Cardiovascular Research, 99:276–283.

Huveneers, S., Oldenburg, J., Spanjaard, E., van der Krogt, G., Grigoriev, I., Akhmanova, A., Rehmann, H., and de Rooij, J. (2012). Vinculin associates with endothelial VE-cadherin junctions to control force-dependent remodeling. Journal of Cell Biology, 196(5):641–652.

Iba, T. and Sumpio, B. E. (1991). Morphological response of human endothelial cells subjected to cyclic strain in vitro. Microvascular Research, 42:245–254.

Janmey, P. A., Fletcher, D. A., and Reinhart-King, C. A. (2020). Stiffness sensing by cells. Physiological Reviews, 100(2):695–724.

Jansen, K. A., Licup, A. J., Sharma, A., Rens, R., MacKintosh, F. C., and Koenderink, G. H. (2018). The role of network architecture in collagen mechanics. Biophysical Journal, 114:2665.

Kotini, M. P., van der Stoel, M. M., Yin, J., Han, M. K., Kirchmaier, B., de Rooij, J., Affolter, M., Huveneers, S., and Belting, H.-G. (2022). Vinculin controls endothelial cell junction dynamics during vascular lumen formation. Cell Reports, 39(2):110658.

Krishnan, R., Canović, E. P., Iordan, A. L., Rajendran, K., Manomohan, G., Pirentis, A. P., Smith, M. L., Butler, J. P., Fredberg, J. J., and Stamenović, D. (2012). Fluidization, resolidification, and reorientation of the endothelial cell in response to slow tidal stretches. American Journal of Physiology-Cell Physiology, 303(4):C368–C375. PMID: 22700796.

Latorre, E., Kale, S., Casares, L., Gómez-González, M., Uroz, M., Valon, L., Nair, R. V., Garreta, E., Montserrat, N., del Campo, A., and et al. (2018). Active superelasticity in three-dimensional epithelia of controlled shape. Nature, 563(7730):203–208.

Lindsey, S. E., Butcher, J. T., and Yalcin, H. C. (2014). Mechanical regulation of cardiac development. Frontiers in Physiology, 5.

Lång, E., Połeć, A., Lång, A., Valk, M., Blicher, P., Rowe, A. D., Tønseth, K. A., Jackson, C. J., Utheim, T. P., Janssen, L. M., Eriksson, J., and Bøe, S. O. (2018). Coordinated collective migration and asymmetric cell division in confluent human keratinocytes without wounding. Nature Communications 2018 9:1, 9:1–15.

López-Gay, J. M., Nunley, H., Spencer, M. A., di Pietro, F., Guirao, B., Bosveld, F., Markova, O., Gaugué, I., Pelletier, S., Lubensky, D. K., and Bellaıche, Y. (2020). Apical stress fibers enable a scaling between cell mechanical response and area in epithelial tissue. Science, 370(6514):eabb2169–eabb2169.

Millán, J., Cain, R. J., Reglero-Real, N., Bigarella, C., Marcos-Ramiro, B., Fernández-Martín, L., Correas, I., and Ridley, A. J. (2010). Adherens junctions connect stress fibres between adjacent endothelial cells. BMC Biology, 8(1).

Narasimhan, B. N., Ting, M. S., Kollmetz, T., Horrocks, M. S., Chalard, A. E., and Malmström, J. (2020). Mechanical characterization for cellular mechanobiology: Current trends and future prospects. Frontiers in Bioengineering and Biotechnology, 8:595978.

Needleman, D. and Dogic, Z. (2017). Active matter at the interface between materials science and cell biology. Nature Reviews Materials, 2(9).

Nejad, M. R., Ruske, L. J., McCord, M., Zhang, J., Zhang, G., Notbohm, J., and Yeomans, J. M. (2023). Stress-shape misalignment in confluent cell layers.

Oldenburg, J. and Rooij, J. D. (2014). Mechanical control of the endothelial barrier. Cell and Tissue Research, 355:545–555.

Pallarès, M. E., Pi-Jaumà, I., Fortunato, I. C., Grazu, V., Gómez-González, M., Roca-Cusachs, P., de la Fuente, J. M., Alert, R., Sunyer, R., Casademunt, J., and Trepat, X. (2022). Stiffness-dependent active wetting enables optimal collective cell durotaxis. Nature Physics 2022 19:2, 19:279–289.

Plateau, J. (1873). Statique expérimentale et théorique des liquides soumis aux seules forces moléculaires. Number v. 1 in Statique expérimentale et théorique des liquides soumis aux seules forces moléculaires. Gauthier-Villars.

Popović, M., Nandi, A., Merkel, M., Etournay, R., Eaton, S., Jülicher, F., and Salbreux, G. (2017). Active dynamics of tissue shear flow. New Journal of Physics, 19(3):033006.

Pourati, J., Maniotis, A., Spiegel, D., Schaffer, J. L., Butler, J. P., Fredberg, J. J., Ingber, D. E., Stamenovic, D., and Wang, N. (1998). Is cytoskeletal tension a major determinant of cell deformability in adherent endothelial cells? American Journal of Physiology - Cell Physiology, 274.

Priya, R., Allanki, S., Gentile, A., Mansingh, S., Uribe, V., Maischein, H.-M., and Stainier, D. Y. R. (2020). Tension heterogeneity directs form and fate to pattern the myocardial wall. Nature, 588(7836):130–134.

Rezakhaniha, R., Agianniotis, A., Schrauwen, J. T., Griffa, A., Sage, D., Bouten, C. V., van de Vosse, F. N., Unser, M., and Stergiopulos, N. (2011). Experimental investigation of collagen waviness and orientation in the arterial adventitia using confocal laser scanning microscopy. Biomechanics and Modeling in Mechanobiology, 11(3–4):461–473.

Salipante, P. F., Hudson, S. D., and Alimperti, S. (2022). Blood vessel-on-a-chip examines the biomechanics of microvasculature. Soft Matter, 18(1):117–125.

Schmidt, U., Weigert, M., Broaddus, C., and Myers, G. (2018). Cell detection with star-convex polygons. Medical Image Computing and Computer Assisted Intervention – MICCAI 2018, page 265–273.

Segal, S. S. (2005). Regulation of blood flow in the microcirculation. Microcirculation (New York, N.Y.: 1994), 12:33–45.

Shore, A. C. (2000). Capillaroscopy and the measurement of capillary pressure. British Journal of Clinical Pharmacology, 50:501.

Shyer, A. E., Tallinen, T., Nerurkar, N. L., Wei, Z., Gil, E. S., Kaplan, D. L., Tabin, C. J., and Mahadevan, L. (2013). Villification: How the gut gets its villi. Science, 342(6155):212–218.

Singh, A., Saha, T., Begemann, I., Ricker, A., Nüsse, H., Thorn-Seshold, O., Klingauf, J., Galic, M., and Matis, M. (2018). Polarized microtubule dynamics directs cell mechanics and coordinates forces during epithelial morphogenesis. Nature Cell Biology, 20(10):1126–1133.

Stoel, M. M. v. d., Kotini, M. P., Schoon, R. M., Affolter, M., Belting, H.-G., and Huveneers, S. (2023). Vinculin strengthens the endothelial barrier during vascular development. Vascular Biology, 5(1).

Takemasa, T., Sugimoto, K., and Yamashita, K. (1997). Amplitude-dependent stress fiber reorientation in early response to cyclic strain. Experimental Cell Research, 230:407–410.

Torres-Sánchez, A., Kerr Winter, M., and Salbreux, G. (2021). Tissue hydraulics: Physics of lumen formation and interaction. Cells Development, 168:203724.

Trubuil, E., D’Angelo, A., and Solon, J. (2021). Tissue mechanics in morphogenesis: Active control of tissue material properties to shape living organisms. Cells Development, 168:203777. Quantitative Cell and Developmental Biology.

Udan, R. S., Culver, J. C., and Dickinson, M. E. (2013). Understanding vascular development: Wire developmental biology (2012). Wiley interdisciplinary reviews. Developmental biology, 2:327.

Vazquez, K., Saraswathibhatla, A., and Notbohm, J. (2022). Effect of substrate stiffness on friction in collective cell migration. Scientific Reports 2022 12:1, 12:1–13.

Williams, S. A., Boolell, M., MacGregor, G. A., Smaje, L. H., Wasserman, S. M., and Tooke, J. E. (1990). Capillary hypertension and abnormal pressure dynamics in patients with essential hypertension. Clinical Science, 79:5–8.

Xi, W., Saw, T. B., Delacour, D., Lim, C. T., and Ladoux, B. (2018). Material approaches to active tissue mechanics. Nature Reviews Materials 2018 4:1, 4:23–44.

Yap, A. S., Duszyc, K., and Viasnoff, V. (2017). Mechanosensing and mechanotransduction at cell–cell junctions. Cold Spring Harbor Perspectives in Biology, 10(8).

