## Supplementary Information for "Interplay of actin nematodynamics and anisotropic tension controls endothelial mechanics"

March 10, 2024

In this Supplementary Information, we first describe the model of nematic tube that we propose to describe the experiments of pressure application on an endothelial tube (Part 1). We first describe how dynamic equations for the tube expansion are obtained from constitutive equations. We then introduce the fitting procedure to experimental data and discuss parameter values. We conclude by discussing model hypothesis and exploring alternative choices of constitutive equations. In the second part of the Supplementary Information, we present the isotropic nonlinear elastic Gent model, used to characterize the deformation of the endothelial tubes under applied pressure (Part 2).

### Part 1 Mechanical model of the endothelial tube at long time scales

In the following, we introduce the mechanical model of nematic active surface that we propose to describe the endothelial tubes. The model is introduced in section 1.1. The fitting procedure to experimental data, and parameter values, are discussed in section 1.2. We discuss model hypothesis in section 1.3.

#### 1.1 Mechanical model of endothelial tube

##### 1.1.1 Differential geometry of a cylindrical tube

We consider the endothelium vessel as an infinitely thin tube of radius $R$ and length $L$ (Fig.4Ai). We take $(\boldsymbol{u}_x, \boldsymbol{u}_y, \boldsymbol{u}_z)$ a cartesian basis of the space such that $\boldsymbol{u}_z$ is oriented along the axis of the cylinder. The surface of the tissue can be parametrized by two variables $\theta$ and $z$ such that the position vector of each point of the surface reads:

$$\boldsymbol{X}(\theta, z) = R\left(\cos\theta\, \boldsymbol{u}_x + \sin\theta\, \boldsymbol{u}_y\right) + z\, \boldsymbol{u}_z, \tag{1.1}$$

with $\theta \in [0, 2\pi]$ and $z \in [0, L]$. Associated to those two variables, we can define a local tangential basis $(\boldsymbol{e}_\theta, \boldsymbol{e}_z)$ at each point of the surface with:

$$\boldsymbol{e}_\theta = \partial_\theta \boldsymbol{X} = R\left(-\sin\theta\, \boldsymbol{u}_x + \cos\theta\, \boldsymbol{u}_y\right), \tag{1.2}$$

$$\boldsymbol{e}_z = \partial_z \boldsymbol{X} = \boldsymbol{u}_z, \tag{1.3}$$

and the local normal vector is:

$$\boldsymbol{n} = \cos\theta\, \boldsymbol{u}_x + \sin\theta\, \boldsymbol{u}_y. \tag{1.4}$$

In this basis, the metric, inverse metric and curvature tensors have the following expressions:

$$g_{ij} = \boldsymbol{e}_i \cdot \boldsymbol{e}_j = \begin{pmatrix} R^2 & 0 \\ 0 & 1 \end{pmatrix}, \tag{1.5}$$

$$g^{ij} = (g_{ij})^{-1} = \begin{pmatrix} 1/R^2 & 0 \\ 0 & 1 \end{pmatrix}, \tag{1.6}$$

$$C_{ij} = -(\partial_i \boldsymbol{e}_j) \cdot \boldsymbol{n} = \begin{pmatrix} R & 0 \\ 0 & 0 \end{pmatrix}, \tag{1.7}$$

and the Christoffel coefficients $\Gamma^k_{ij}$ vanish. In the following, we consider the tissue to be uniform, axisymmetric and invariant by translation, so that physical quantities do not depend on the coordinates $\theta$ and $z$; therefore their covariant derivatives vanish. We consider that the vessel remains cylindrical, with its length $L$ fixed and its radius $R$ changing in time. Thus, even if locally the endothelial cells can have tangential displacement, the mean flow at the surface of the cylinder is purely normal to the surface of the tissue, and the velocity is simply $\boldsymbol{v} = v_n \boldsymbol{n}$. The normal velocity is given by $v_n = \mathrm{d}R/\mathrm{d}t$, with the dot notation indicating a temporal derivative. The strain-rate tensor then reads:

$$v_{ij} = C_{ij} v_n = \begin{pmatrix} R\frac{\mathrm{d}R}{\mathrm{d}t} & 0 \\ 0 & 0 \end{pmatrix}, \quad v_i{}^j = \begin{pmatrix} \frac{1}{R}\frac{\mathrm{d}R}{\mathrm{d}t} & 0 \\ 0 & 0 \end{pmatrix} \tag{1.8}$$

##### 1.1.2 Dynamics of the actin fibers mean orientation tensor

Endothelial cells exhibit a network of actin stress fibers (Fig. 2Ci). The orientation of each stress fiber can be described by an unitary vector $n_i$. As we aim to build a mesoscopic theory, we construct the mean orientation tensor $n_{ij} = \langle n_i n_j \rangle$, where the operator $\langle . \rangle$ indicates the average over all the actin fibers. The tensor $n_{ij}$ describes the orientation of a set of fibers with a single mesoscopic quantity. This tensor is symmetric, with trace equal to 1. The tensor $n_{ij}$ can be decomposed into an isotropic and a traceless part as follows:

$$n_{ij} = \frac{g_{ij}}{2} + q_{ij}, \tag{1.9}$$

where $q_{ij}$, the traceless part of $n_{ij}$, measures the anisotropy of the fibers orientation.

In the following we consider that off-diagonal coefficients of $n_{ij}$ and $q_{ij}$ vanish, as experimental measurements indicate that their values are small (Fig. S4Bi,Ci). Because $q_{ij}$ has zero off-diagonal coefficients and is a traceless tensor, it has only one independent component $q = q_\theta^\theta = -q_z^z$:

$$q_{ij} = \begin{pmatrix} R^2 q & 0 \\ 0 & -q \end{pmatrix}, \; q_i{}^j = \begin{pmatrix} q & 0 \\ 0 & -q \end{pmatrix}. \tag{1.10}$$

The value of $q$ varies between $-1/2$ and $1/2$: for $q = -1/2$ all fibers are aligned with the tube axis, whereas for $q = 1/2$ all fibers are oriented circumferentially.

**Passive deformation by tissue shear.** Does the change of orientation of actin fibers occurs simply by following the tissue shear, or is it driven by additional effects? To determine the contribution of material flow to actin stress fibers reorientation, we computed the evolution of $q$ as a function of a numerically applied uniaxial strain in the circumferential direction, for different initial distributions of orientation. To generate initial distributions, we extracted a set of vectors oriented along actin stress fibers from immunostaining images of the actin filaments (Fig. S4A, Methods). We then applied a given uniaxial strain along the circumferential direction by deforming each vector individually. The order parameter $q$ was computed from the new distribution of angles of the strained vector field. We repeated this operation for strains ranging from 0 to 100%, for six different initial actin immunostaining images, to evaluate $q$ as a function of $R$ (Fig. 4B). We then compared this prediction to the experimentally measured order parameters $q$ for different radii. This comparison shows that the increase of $q$ due to deformation by the material flow explains less than 20% of actually observed actin stress fibers reorientation; indicating that other effects are driving actin stress fibers dynamics. For simplicity, we thus neglect the effect of tissue shear on actin reorientation in the model.

**Actin fibers mechanosensitivity.** Experimental observations suggest that actin stress fibers are in an ordered phase (Fig. 2C), and that their orientation is sensitive to the direction of tension (Fig. 2C, 3). We therefore choose the following equation for the dynamics of actin fibers nematic order:

$$D_t q_{ij} = -\gamma \left( \frac{1}{2} q_{kl} q^{kl} - q_0^2 \right) q_{ij} + \beta \tilde{t}^r_{ij}, \tag{1.11}$$

where $1/\gamma$ is the characteristic relaxation time of $q_{ij}$, $q_0$ is a preferred order magnitude, $\beta$ is a mechanosensitive coupling term to $\tilde{t}^r_{ij} = t^r{}_{ij} - t^{r\,k}{}_k \, g_{ij}/2$, the traceless part of the residual tension $t^r{}_{ij}$, introduced below in Eq. 1.18. The operator $D_t$ denotes the corotational derivative, with expression for a symmetric tensor $A_{ij}$:

$$D_t A_{ij} = \partial_t A_{ij} + v^k \nabla_k A_{ij} + \omega_{ik} A^k{}_j + \omega_{jk} A_i{}^k, \tag{1.12}$$

with $\omega_{ij} = 1/2(\nabla_i v_j - \nabla_j v_i)$. The first term in the right-hand side of Eq. 1.11 sets a preferred magnitude $q_0$ for actin stress fibers ordering. In the second term, actin stress fibers are assumed to respond to tension anisotropy in the tissue and to align with the direction of maximal residual tension $t^r_{ij}$. Here we exclude the contribution of tension within the actin stress fibers themselves to this feedback effect. We consider in section 1.3.5 the case where a coupling to the total tension of the tissue is replacing the coupling to $t^r_{ij}$.

##### 1.1.3 Force balance

In the following we consider that the tissue is subjected to an internal tangential tension $t_{ij}$. Since the tube dynamics operates at low Reynolds number, the force balance in the normal direction reads

$$C_{ij}t^{ij} = \Delta P, \tag{1.13}$$

where $\Delta P = P - P_{\text{gel}}$ is the difference of pressure across the endothelial monolayer, equal to the difference between the luminal pressure and the surrounding gel pressure. We consider the pressure in the gel $P_{\text{gel}}$ to be fixed and equal to 0, and therefore ignore possible contributions from the gel elasticity. Indeed, we find that the gel elastic modulus at concentrations used in experiments ($\sim 500 - 1000$Pa, [Jansen et al., 2018]) is considerably softer than the elastic modulus of the tissue ($\sim 100 - 200$kPa, Fig. 1Cii). In addition, collagen gels with similar concentrations have been shown to relax elastic stresses on the timescales of the order of an hour, a short time compared to our experiment lasting $\sim 50$ hours [Nam et al., 2016, Ban et al., 2018, Kim et al., 2017].

Since the curvature tensor has only one non-zero component $C_{\theta\theta}$ (Eq. 1.7), one obtains simply the law of Laplace:

$$t_\theta^\theta = R\Delta P. \tag{1.14}$$

In the following we assume that the pressure difference between the fluid in the tube and the surrounding gel, $\Delta P$, and all other physical quantities, are invariant along the longitudinal axis of the tube. In practice a flow has to be maintained in the tube for cell survival, which creates a gradient of pressure along the tube longitudinal direction of $\simeq 100$Pa (Methods). In the following we use average values of pressures inside the tube for our analysis.

##### 1.1.4 Constitutive equation for the tension

We discuss here the constitutive equation for the tissue tension tensor $t_{ij}$. In proposing constitutive equations for this tension tensor, we adopted the following rationale. Following previous studies [Etournay et al., 2015, Popović et al., 2017], we first reasoned that resistance to tissue expansion could generally arise from elastic resistance to cell area or to cell elongation. In addition, the strong reorientation of actin fibers observed experimentally suggested that anisotropic tension arising from the actomyosin network could play a role.

However, experimental data showed that both the average cell area and average magnitude of cell elongation drop 56 hours after the application of the increased pressure to the tube (Fig. 2Div, 4Eii). If cells had a fixed value of preferred area and elongation, this would imply that elastic stresses related to area and elongation should also decrease over time. However, the vessel radius increases continuously over the time course of the experiment, such that the tension in the circumferential direction $t_\theta^\theta$ must also increase, according to Eq. 1.14. This indicates that the tension in the tissue is not dominated by cell area and cell elongation elasticity. In addition, we note that the average cell density varies by a factor 4 in different tissues, suggesting that cell area is not strongly regulated (Fig. 4Eiii). Therefore, we propose that the tension generated by actomyosin fibers is a key player in the tissue mechanics.

**Tension generated by actomyosin fibers.** We assume that the tension within the network of actin stress fibers lays along the mean orientation of fibers:

$$t_{ij}^a = \zeta n_{ij}, \tag{1.15}$$

where $\zeta$ is the magnitude of the resulting tension. The tension has both an isotropic part (proportional to $g_{ij}/2$) and an anisotropic part (proportional to $q_{ij}$) with equal magnitudes. The simplest description of actin generated tension would be to consider that every fiber is generating the same constant tension $\zeta_f$. In this case $\zeta = \rho\zeta_f$ would be constant, if we consider the density of fibers $\rho$ to be fixed. However, we find that this would not be compatible with the dynamics of tube expansion under pressure. Indeed, $t_\theta^\theta = \zeta(q + 1/2)$ would then plateau at long time, as $q$ reaches a maximum value after 24 hours (Fig. 2Civ) and $\zeta$ is constant. As noted above, the law of Laplace implies that $t_\theta^\theta$ increases steadily in time. This suggests that actin stress fibers should generate a circumferential tension increasing in time. We therefore assume that actin stress fibers respond elastically to material deformation. If an individual actin fiber behaves as a spring of stiffness $K_a$ and orientation $n_i$ and deforms with the tissue, the tension generated by this fiber is $s_{ij} = K_a s\, n_i n_j$, with $s$ the elongational stretch of the spring. The relative velocity of the two extremities of the fiber due to the material flow is $v_i{}^k n_k$, such that its elongation rate is $D_t s = v^{kl} n_k n_l$, with $D_t s = \partial_t s + v^k \nabla_k s$. Inspired by this result for a single spring-like actin fiber, we choose the following constitutive equation for the tension generated by actin stress fibers:

$$t^a_{ij} = (\zeta_0 + K_a s)\, n_{ij}, \tag{1.16}$$

$$D_t s = v^{kl} n_{kl}. \tag{1.17}$$

where $s$ corresponds to an average level of fiber elongational stretch and $\zeta_0$ is the amplitude of the active tension generated by actin fibers.

**Additional tension generated by the tissue.** Laser ablation experiments show that a few minutes after the application of the additional pressure, the tension is strongly anisotropic, being higher along the circumferential direction (Fig. 1Bii), which is in contradiction with predictions of Eq. 1.16. Indeed, just after application of the pressure increase, actin fibers are still mainly oriented along the tube axis, implying that $t^a_{ij}$ has higher magnitude along the longitudinal than circumferential direction of the tube. We therefore propose that the tension in the tissue does not only depend on actin stress fibers, and can be decomposed into two contributions:

$$t_{ij} = t^a_{ij} + t^r_{ij}\ , \tag{1.18}$$

corresponding respectively to the tension generated by the cellular actomyosin network, and a residual tension acting within the other constituents of the tissue. Isotropic tension arising from cell area elasticity would not change the tension anisotropy. Cell elongation elasticity will also not lead to circumferentially oriented tension just after pressure step application, because cell elongation is still mainly along the tube direction. Therefore, we assume that other elements in the tissue mechanically resist its deformation. These elements could for instance correspond to microtubules, cortical actin, cell membrane or intermediate filaments. We postulate that this residual contribution arises from a Maxwell viscoelastic response to tissue shear, with short time elasticity and long time viscosity:

$$(1 + \tau D_t)\, t^r_{ij} = \mu v_{ij}, \tag{1.19}$$

where $\tau$ is the characteristic time of the Maxwell model, $\mu$ a viscosity and $D_t$ the corotational derivative defined in Eq. 1.12. We discuss this choice of constitutive relation for $t^r_{ij}$ in Sec. 1.3.3.

##### 1.1.5 Constitutive equations for cell area and cell elongation dynamics

Here we discuss the dynamics of the mean cell area $a$ and mean cell elongation $Q_{ij}$ which are coupled to the tissue expansion.

**Area dynamics.** The area dilatation rate of an infinitesimal piece of a surface is equal to the local isotropic growth rate $v^k_k$ of this surface. Cells also divide at a rate $k_d$, that we assume to be constant. Assuming that cell apoptosis or delamination can be neglected, the dynamics of the mean cell area then reads [Etournay et al., 2015, Popović et al., 2017]:

$$\frac{\partial_t a}{a} = v^k_k - k_d. \tag{1.20}$$

**Cell elongation dynamics.** We now discuss the dynamics of the cell elongation tensor $Q_{ij}$. The surface deformation contributes to cell elongation through its deviatoric part $\tilde{v}_{ij} = v_{ij} - v^k_k g_{ij}/2$. However, this purely geometric contribution is not sufficient to explain the cell elongation measured 7 and 24 hours after additional pressure increase (Fig. 4Fiii). Indeed we observe at these two times an excess of circumferential cell elongation measured experimentally relative to the value predicted by the geometrical contribution only. Moreover, we observe a strong correlation between cell elongation and actin orientation (Fig. 4Fii), suggesting that the cell elongation tensor $Q_{ij}$ follows the dynamics of the actin stress fiber nematic tensor $q_{ij}$. We therefore propose the following dynamical equation for $Q_{ij}$:

$$D_t Q_{ij} = \tilde{v}_{ij} - \lambda(Q_{ij} - \alpha q_{ij}) \tag{1.21}$$

where $1/\lambda$ is the characteristic time of alignment of cell elongation with the actin stress fibers, and $\alpha$ is a proportionality factor between the cell elongation and actin stress fibers nematic tensor. The first term in the right-hand side corresponds to cell elongation changes arising from the tissue anisotropic shear, and the last term in the right-hand side arises from cellular rearrangements in the tissue tending to relax the cell elongation tensor towards the actin fiber tensor. [Etournay et al., 2015, Popović et al., 2017].

##### 1.1.6 Dimensionless system of equations for tube radius, nematic order, cell area and elongation

To obtain dimensionless equations, we choose $R_0$, the radius of the tube at $t = 0$, before the application of the additional pressure, as length unit, $\tau$, the Maxwell viscoelastic time, as time unit, and $\zeta_0$, the constant actin tension, as unit of tension (Table 1.1).

| Physical quantity | Unit |
|---|---|
| Length | $R_0$ |
| Time | $\tau$ |
| Tension | $\zeta_0$ |

Table 1.1: Physical variables used for non-dimensionalisation.

We transform variables and parameters such as $R/R_0 \longrightarrow R$, $t/\tau \longrightarrow t$, $R_0\Delta P/\zeta_0 \longrightarrow \Delta P$, $K_a/\zeta_0 \longrightarrow K_a$, $\mu/(\tau\zeta_0) \longrightarrow \mu$, $\gamma\tau \longrightarrow \gamma$, $\tau\beta\zeta_0 \longrightarrow \beta$, $\tau k_d \longrightarrow k_d$, $\tau\lambda \longrightarrow \lambda$. We also introduce $\sigma = t^{r}{}^{\theta}_{\theta}/\zeta_0$ and $Q = Q^{\theta}_{\theta} = -Q^{z}_{z}$, as $Q_{ij}$ is a traceless tensor. We consider that off-diagonal coefficients of $Q_{ij}$ vanish, as experimental measurements indicate that their value is small (Fig. S4Bii, Cii). We then obtain the following system of dimensionless equations:

$$R\Delta P = \sigma + (1 + K_a s)\left(q + \frac{1}{2}\right), \tag{1.22}$$

$$\left(1 + \frac{\mathrm{d}}{\mathrm{d}t}\right)\sigma = \frac{\mu}{R}\frac{\mathrm{d}R}{\mathrm{d}t}, \tag{1.23}$$

$$\frac{\mathrm{d}s}{\mathrm{d}t} = \left(q + \frac{1}{2}\right)\frac{1}{R}\frac{\mathrm{d}R}{\mathrm{d}t}, \tag{1.24}$$

$$\frac{\mathrm{d}q}{\mathrm{d}t} = -\gamma\left(q^2 - q_0^2\right)q + \frac{1}{2}\beta\sigma, \tag{1.25}$$

$$\frac{1}{a}\frac{\mathrm{d}a}{\mathrm{d}t} = \frac{1}{R}\frac{\mathrm{d}R}{\mathrm{d}t} - k_d, \tag{1.26}$$

$$\frac{\mathrm{d}Q}{\mathrm{d}t} = \frac{1}{2R}\frac{\mathrm{d}R}{\mathrm{d}t} - \lambda(Q - \alpha q). \tag{1.27}$$

The dimensionless expressions of physical quantities and coefficients are given in Table 1.2.

##### 1.1.7 Initial conditions and dynamics of pressure application

We discuss here initial conditions and the dynamics of pressure $\Delta P$ which have to be specified to solve Eqs. 1.22-1.27.

In the experimental protocol, cells are seeded on the tube surface and allowed to spread on the gel and proliferate until reaching confluence. The time of confluence is defined as $t = t_c = -24$hours. After confluence, the cells form an low-permeability barrier and a small pressure difference of $\Delta P_0 \simeq 150$Pa is then established within the tube. This low pressure stems from the the minimal fluid flow in the tube necessary to maintain cells alive. This state is maintained during 24 hours to mature the monolayer. The tube radius increases slightly by $\simeq 5\%$ in 24 hours at this low pressure. At $t = 0$, a larger pressure difference $\Delta P_m$ is applied.

| | | Dimensionless expression |
|---|---|---|
| Physical variables | $R$ | $R/R_0$ |
| | $t$ | $t/\tau$ |
| | $q$ | $q$ |
| | $\Delta P$ | $R_0\Delta P/\zeta_0$ |
| | $\sigma$ | $\sigma/\zeta_0$ |
| Parameters of the model | $\tau$ | 1 |
| | $\zeta$ | 1 |
| | $K_a$ | $K_a/\zeta_0$ |
| | $\mu$ | $\mu/(\tau\zeta_0)$ |
| | $\gamma$ | $\tau\gamma$ |
| | $q_0$ | $q_0$ |
| | $\beta$ | $\tau\beta\zeta_0$ |
| | $k_d$ | $\tau k_d$ |
| | $\lambda$ | $\tau\lambda$ |
| | $\alpha$ | $\alpha$ |

Table 1.2: List of physical variables and model parameters, and corresponding dimensionless expression.

We consider that just before reaching confluence ($t = t_c^-$ with $t_c = -24$h), the assembly of disjointed cells is not impermeable to the luminal fluid and the pressure difference $\Delta P$ vanishes. There is then no circumferential tension in the tissue, and we further assume that actin fibers are unstretched, $s = \sigma = 0$. The first term in the right-hand side of Eq. 1.25 allows both $q = \pm q_0$ as a reference state. Here we consider that actin fibers in the tube initially prefer to align along the tube axis, possibly due to minimization of bending energy, thus selecting $q = -q_0$. For simplicity, we consider that the pressure difference $\Delta P$ increases from zero to $\Delta P_0$ at the establishment of confluence ($t = t_c$) on a time scale much faster than the characteristic times of the model, such that only $R$, $s$ and the elastic part of $\sigma$ respond to the pressure change and that $a$ and $Q$ jump according to the tissue deformation. Then just after confluence ($t = t_c^+$), one obtains:

$$s(t_c^+) = \left(-q_0 + \frac{1}{2}\right) \ln \frac{R(t_c^+)}{R(t_c^-)}, \tag{1.28}$$

$$\sigma(t_c^+) = \mu \ln \frac{R(t_c^+)}{R(t_c^-)}, \tag{1.29}$$

$$R(t_c^+)\Delta P_0 = \sigma(t_c^+) + (1 + K_a s(t_c^+)) \left(-q_0 + \frac{1}{2}\right), \tag{1.30}$$

$$a(t_c^+) = \frac{a(t_c^-)R(t_c^+)}{R(t_c^-)}, \tag{1.31}$$

$$Q(t_c^+) = Q(t_c^-) + \frac{1}{2} \ln \frac{R(t_c^+)}{R(t_c^-)} \tag{1.32}$$

$R(t_c^+)$ is then determined by replacing $s(t_c^+)$ and $\sigma(t_c^+)$ by their expression as a function of $R(t_c^+)$ (Eqs. 1.28 and 1.29) in Eq. 1.30 and solving the resulting equality. The value of all other quantities at $t = t_c^+$ are determined by $R(t_c^+)$. We then solve the dynamical equations Eqs. 1.22-1.27 with luminal pressure $\Delta P = \Delta P_0$, for $t_c^+ \leq t \leq 0^-$. Value of the variables $R$, $q$, $\sigma$ and $s$ at $t = 0^-$ are obtained from this solution. The normalised value of $R$ at $t = 0^-$ is imposed to be equal to 1 by rescaling the solution. We consider that the pressure increases at $t = 0$ from $\Delta P_0$ to $\Delta P_m$ much faster than the characteristic times of the model such that only $R$, $s$ and the elastic part of $\sigma$ change value to adapt to the change of pressure, and $a$ and $Q$ jump according to the tissue deformation. This gives:

$$s(0^+) = s(0^-) + \left(q(0^-) + \frac{1}{2}\right) \ln R(0^+), \tag{1.33}$$

$$\sigma(0^+) = \sigma(0^-) + \mu \ln R(0^+), \tag{1.34}$$

$$R(0^+)\Delta P_m = \sigma(0^+) + (1 + K_a s(0^+))\left(q(0^-) + \frac{1}{2}\right), \tag{1.35}$$

$$a(0^+) = a(0^-)R(0^+), \tag{1.36}$$

$$Q(0^+) = Q(0^-) + \frac{1}{2}(0^+) \tag{1.37}$$

and $q$ is continuous, $q(0^+) = q(0^-)$. As done previously, Eq. 1.35 can be used to determine the value of $R(0^+)$ that sets the value of all other quantities at $t = 0^+$. We then solve the dynamical equations 1.22-1.27 for $t \geq 0$.

#### 1.2 Model comparison to experimental data

##### 1.2.1 Fitting procedure

**Fitting of tube radius and actin order parameter dynamics.** We performed a fitting procedure to infer the parameters of the model that reproduce the measured behaviour of the dimensionless values of $R$ and $q$. We use a Nelder-Mead algorithm to minimize the following cost function:

$$C(\{p\}) = A \times \left[R(\{p\}, t_c^-) - R_c\right]^2 + \sum_{i=1}^{N_R}\left(\frac{R(\{p\}, t_i/\tau) - \bar{R}_i/R_0}{\bar{R}_i/R_0}\right)^2 + \sum_{\alpha=1}^{N_q}\left(\frac{q(\{p\}, t_\alpha/\tau) - \bar{q}_\alpha}{q^*}\right)^2, \tag{1.38}$$

where $\{p\}$ is a set of value of the parameters of the model and $R(\{p\}, t)$ (resp. $q(\{p\}, t)$) the value of the variable $R$ (resp. $q$) at time $t$ when the model is solved numerically with parameters value $\{p\}$.

Experimental data consist in two set of measurements: $\bar{R}_i$ the value of the radius measured at time $t_i$, and $\bar{q}_\alpha$ the value of $q$ measured at time $t_\alpha$. There are $N_R$ measured values of the radius $R$ and $N_q$ measured values of the order parameter $q$. $q^*$ is the mean value of measurements of $q$ performed at $t = 0^-$ and $R_c$ the experimental value of the normalized tube radius when tissue reaches confluence. To enforce a $\simeq 5\%$ radius increase between $t_c^-$ and $t = 0^-$ as observed in experiments we take $R_c = 0.96$ and $A = 2000$.

**Fitting of cell area dynamics.** The optimal value of the cell division rate $k_d$ is determined as follows, from experimental measurements of the cell mean area $a$. We use a fitting algorithm to minimize the following cost function:

$$C_a(k_d) = \sum_{i=1}^{N_a}\left(\tilde{a}\left(k_d, \frac{t_i}{\tau}\right) - \frac{\bar{a}_i}{a_0}\right)^2, \tag{1.39}$$

with $a_0$ the mean value of measured mean cell areas at $t = 0^-$. $\tilde{a}(k_d, t)$ is the value of $a/a_0$ at time $t$ computed thanks to Eq. 1.26 with a specified value of $k_d$ and the initial condition $\tilde{a}(0^-) = 1$. The measured mean cell areas are $\{\bar{a}_i\}_{i=1..N_a}$, with $t_i$ the time at which $\bar{a}_i$ has been measured and $N_a$ the number of measurements. All the parameters of the mechanical model (Eqs. 1.22-1.25) have been fixed to their optimal value determined by the previous fit on $R$ and $q$. The best fit for the evolution of the cell area as a function of time is shown in Fig. 4Eii.

**Fitting of cell elongation dynamics.** Following a similar procedure, we determine best fit values for the parameters $\lambda$ and $\alpha$ involved in the dynamics of cell elongation. The cost function we use for the fit now reads:

$$C_Q(\lambda, \alpha) = \sum_{i=1}^{N_Q}\left(Q\left(\lambda, \alpha, \frac{t_i}{\tau}\right) - \bar{Q}_i\right)^2, \tag{1.40}$$

with $Q(\lambda, \alpha, t)$ the value of $Q$ at time $t$ computed from Eq. 1.27 for the specified values of $\lambda$ and $\alpha$ with initial condition $Q(0^-) = Q_0$, where $Q_0$ is the average of experimental values of $Q$ at $t = 0^-$. The experimental values of cell elongation are $\{\bar{Q}_i\}_{i=1..N_Q}$, with $t_i$ the time at which $\bar{Q}_i$ has been measured and $N_Q$ the number of measurements. All parameters of the mechanical model (Eqs. 1.22-1.25) and $k_d$ have been fixed to their optimal value determined by previous fits. The dynamics of $Q$ with optimal values of $\lambda$ and $\alpha$ is shown in Fig. 4Fiii.

| | |
|---|---|
| Measured parameters | $R_0 = 64 \pm 3\,\mu m$ |
| | $\Delta P = 650 \pm 50\,Pa$ |
| Dimensionless parameters | $\Delta\hat{P} = 1.4$ |
| | $\hat{\mu} = 9.9$ |
| | $\hat{K}_a = 7.6$ |
| | $\hat{\gamma} = 2.9$ |
| | $q_0 = 0.26$ |
| | $\hat{\beta} = 0.38$ |
| | $\hat{k}_d = 0.04$ |
| | $\hat{\lambda} = 4.5$ |
| | $\alpha = 0.46$ |
| Dimensional parameters | $\tau = 3.4h$ |
| | $\zeta_0 = 0.03N/m$ |
| | $K_a = 0.22N/m$ |
| | $\mu = 0.98N/m.h$ |
| | $K = \mu/\tau = 0.29N/m$ |
| | $\gamma = 0.87h^{-1}$ |
| | $\beta = 3.8N^{-1}.m.h^{-1}$ |
| | $k_d = 0.011h^{-1}$ |
| | $\lambda = 1.3h^{-1}$ |

Table 1.3: Table of fitted parameters. Hatted symbols indicated dimensionless parameters according to Tab. 1.2 while unhatted symbols indicate parameters with physical dimensions.

##### 1.2.2 Discussion of best fit results

Optimal values of model parameters are given in Table 1.3.

**Dynamics of tube radius and actin order parameter.** The dynamic evolution of $R$ and $q$ is well captured using these best fit parameters (Fig. 4Di,ii). At short times after the pressure step, the residual tension $t^r_{ij}$ resists deformation elastically, which leads to circumferential reorientation of actin fibers (Fig. 4C, Div). This contribution then relaxes on a characteristic time $\tau$ due to the viscoelastic behaviour. The actin generated tension $t^a_{ij}$, now oriented circumferentially, then resists tissue expansion, such that at long time the dynamics of the tube is dominated by actin contribution (Fig. 4C, Div). To illustrate this point, we show the dynamics of the system when the elastic contribution of tension in actin fibers is removed, $K_a = 0$ (with all other parameters kept at the same value, Fig. 4Di,ii). In this case, the radius diverges in a few hours. We also highlight the importance for actin fibers to circumferentially reorient by simulating the dynamics of the system with actin-tension coupling term $\beta = 0$ (with all other parameters kept at the same value, Fig. 4Di,ii). In this case the radius also increases quickly, showing the importance of fibers aligning with the flow to resist the expansion of the tube.

**Dynamics of tissue tension.** The evolution of the components $t^\theta_\theta$ and $t^z_z$ in the model is shown in Fig. 4Diii. In the model, the longitudinal tension $t^z_z$ is initially larger than the circumferential tension $t^\theta_\theta$. The difference is larger than detected by laser ablation (Fig. 1Bii, S1A), possibly due to the weak longitudinal anisotropy of the high-density monolayers used for these experiments. At $t = 0^+$ $t^\theta_\theta$ jumps by a larger amount than $t^z_z$ such that the tension just after pressure step application is strongly anisotropic in circumferential direction, in agreement with laser ablation experiments.

**Dynamics of cell area and cell elongation.** The coupling between cell elongation and actin fiber elongation is resulting in an excess of elongation at 7 and 24 hours compared to the deformation expected from tissue shear, as observed experimentally (Fig. 4Fiii). We also show the dynamics of $Q$ for a pressure of 150Pa, starting at $t = t_c$ with the initial condition $Q(t_c^-) = \alpha q_0$. The model however does not account for a drop of elongation at 56 hours. We suspect that this drop could be due to a coupling of cell elongation to the cell area, possibly via the oriented divisions reported in experiments, as cells with small areas are observed to be preferentially isotropic.

##### 1.2.3 Discussion of the inferred parameters value

The characteristic active tension generated by actin is $\zeta_0 \approx 0.03N/m$. The characteristic tension generated by its resistive part is $\sim K_a s_{max} \approx 0.06N/m$, with $s_{max} \approx 0.25$, the maximum value of $s$ (Fig. S4E). The characteristic tension generated by the rest of the tissue is $\sim (\mu/\tau)\ln(R_0^+) \approx 0.04N/m$, with $R_0^+$ the normalized radius of the vessel just after the increase of pressure to 650Pa. Each of those three contributions are of the same order of magnitude, towards large values of the previously reported range of tensions [Duque et al., 2023, Marin-Llaurado et al., 2023]. This is broadly consistent with experimental observations that the stiffness of endothelial or epithelial tissues is divided by approximately two when actin is depolymerized [Harris et al., 2012, Pourati et al., 1998].

The characteristic time of actin reorganisation $\sim 1/\gamma \approx 70$min is of the same order of magnitude as previously reported values of 5-40min of actin or cellular tension reorientation under cycling stretching [Takemasa et al., 1997, Iba and Sumpio, 1991, Hayakawa et al., 2001, Krishnan et al., 2012]. The non-dimensional coefficient $2q_0^2\gamma\tau/(\beta\mu) \approx 0.11$ is a characteristic strain inducing actin reorientation away from its original orientation. This is comparable to observations that a 10% strain is enough to induce actin fibers reorientation in endothelial cells [Kaunas et al., 2005].

The characteristic time of cell elongation relaxation towards actin orientation is $\sim 1/\lambda \approx 45$min. This small time scale relative to the total time of the experiment (56 hours) implies that $Q$ correlates well with $q$, in accordance to the observed correlation throughout the experiment (Fig. 4Fii).

The doubling time of cell is $\sim \ln(2)/k_d \approx 60h$, in agreement with experimental measurements of the number of divisions between 0 and 7 hours (Methods), which are consistent with reported values for comparable cell densities [Gong et al., 2015, Vasiceek et al., 2021].

##### 1.2.4 Effect of the magnitude of applied pressure

To investigate the effect of the applied pressure at $t > 0$ on the tube expansion dynamics, we solve the model numerically for pressures of $\Delta P_m = 450$ Pa and 850 Pa, in addition to the value of $\Delta P_m = 650$ Pa used for fitting experimental data. The first stage of imposed pressure difference $\Delta P = 150$ Pa for $t_c < t < 0$ remains unchanged, such that for all pressures the tube is considered to be in the same state at $t = 0^-$. We show in Fig. S4G the dynamics of the system for different pressures. Comparison of these results with experimental measurements of $R$, $q$, $a$ and $Q$ for different pressures, 7 hours after the pressure step (Fig. 4G), show that the model is in good agreement with experimental values.

##### 1.2.5 Pressure arrest experiment

Here we investigate the change in tube radius following experiments where the applied pressure $\Delta P$ is reduced abruptly after 7 hours of tube expansion, from the reference value $\Delta P_m$ to the residual pressure $\Delta P_0$ (Fig. 2Bi). We assume that the reduction in pressure occurs on a time scale much smaller than characteristic times of the model. Then only $R$, $s$ and the elastic contribution of $\sigma$ can immediately respond to the change of pressure. The state of the system just after the pressure arrest is then obtained by solving the following relations:

$$s^+ = s^- + (q^- + \frac{1}{2}) \ln \frac{R^+}{R^-}, \tag{1.41}$$

$$\sigma^+ = \sigma^- + \mu \ln \frac{R^+}{R^-}, \tag{1.42}$$

$$R^+ \Delta P_0 = \sigma^+ + (1 + K_a s^+)(q^- + \frac{1}{2}), \tag{1.43}$$

where the upperscript − (resp. +) accounts for the value of the variables just before (resp. after) the pressure arrest.

We find that the pressure release leads to a relaxation of the tube radius of similar magnitudes between theory and experiment. Experimentally, the instantaneous relative change of radius following pressure variation is smaller at 7 hours than at 0 hour. In the model however we find that this change is predicted to be rather constant (Fig. S4D). This discrepancy may be linked to strain-stiffening properties of the tissue which we have not considered here for simplicity.

#### 1.3 Discussion of the model

In this section we discuss the stability of steady-state cylindrical solutions of the nematic mechanical model introduced in section 1.2. We then discuss the different model hypothesis by considering alternative constitutive equations for the actin stress fibers nematodynamics and tissue tension, summarized in Tab. 1.4.

##### 1.3.1 Stability analysis of steady-state solutions

Here we discuss steady-state solutions of the model for cylindrical tubes. At steady state, the tension $t^r_{ij}$ vanishes, and the order parameter $q = \pm q_0$ (we consider here solutions where the alignment is circumferential or longitudinal). In this subsection, we use dimensional equations. Mechanical balance then imposes that:

$$R_e \Delta P = t^\theta_\theta = (\zeta_0 + K_a s_e)\left(\frac{1}{2} \pm q_0\right) \tag{1.44}$$

with $R_e$ and $s_e$ the tube radius and fiber strain at steady-state.

To test for the stability of this solution, we consider a perturbation $R = R_s + \delta R$, $s = s_e + \delta s$, $q = q_e + \delta q$ with $q_e = \pm q_0$. At linear order in the perturbation, one obtains

$$(1 + \tau\partial_t)\partial_t \delta q = -2\gamma q_0^2 (1 + \tau\partial_t)\delta q + \frac{\beta\mu}{2}\frac{\partial_t \delta R}{R_e} \tag{1.45}$$

$$(1 + \tau\partial_t)\frac{\delta R}{R_e}\left(\frac{R_e \Delta P}{K_a} - \left(\frac{1}{2} + q_e\right)^2\right) = \frac{\mu}{K_a}\frac{\partial_t \delta R}{R_e} + \frac{\zeta_0 + K_a s_e}{K_a}(1 + \tau\partial_t)\delta q\ . \tag{1.46}$$

Performing a Laplace transform, we can ask when perturbations relax exponentially, corresponding to a stable solution. We find that pressure values satisfying

$$\Delta P < \frac{K_a\left(\frac{1}{2} + q_e\right)^2}{R_e}\ . \tag{1.47}$$

give rise to stable solutions. This also corresponds to a range of stable strain values:

$$s_e < \frac{1}{2} + q_e - \frac{\zeta_0}{K_a}. \tag{1.48}$$

The solution where fibers are circumferential ($q_e = q_0$) is therefore more stable than the solution where fibers are longitudinal ($q_e = -q_0$). With parameters in Table 1.3, this corresponds to a maximal strain $s_e \simeq 0.62$ for circumferential fibers alignment and $s_e \simeq 0.1$ for longitudinal alignment. This highlights the role of actin fibers reorientation in our model of vessel mechanics. Indeed, with longitudinal alignment stable solutions only exist for low strain until reaching a relatively low value $s_e$. As a result, when setting the tension coefficient $\beta$ coupling actin orientation to tissue tension to zero, the tube radius diverges exponentially (Fig. 4Dii) instead of simply reaching a higher steady value. However, even with circumferentially-oriented actin stress fibers, stable solutions disappear for large enough strain, indicating that the tube will not reach an equilibrium radius if the pressure crosses a threshold value. We find numerically that for pressures greater than $\simeq$ 780Pa the strain of actin fibers overcomes its critical value and the tube radius does not reach steady value (Fig. S4Gi).

##### 1.3.2 Viscoelastic response of actin stress fibers

In the constitutive equation for actin stress fibers (Eq. 1.16) we assumed that actin fibers respond elastically to deformation of the tube. This implies that actin fibers have a infinite memory of their initial reference state. Cellular processes may however lead to a loss of this memory. For instance, the actin cytoskeleton undergoes dramatic remodeling during cell division,, with the disassembly of stress fibers during mitosis (Fig. S5B). A simple way to take into account such memory loss is to consider a viscoelastic response of actin stress fibers:

$$t^a_{ij} = (\zeta_0 + \sigma_a)n_{ij}, \tag{1.49}$$

$$(1+\tau_a D_t)\sigma_a = \mu_a v^{kl} n_{kl}, \tag{1.50}$$

with $\mu_a = \tau_a K_a$, where $\tau_a$ is actin tension relaxation time.

We fitted experimental data of tube radius and actin orientation by optimizing parameters of this model through a Nelder-Mead algorithm minimizing the following cost function:

$$C(\{p\}) = A \times \left[R(\{p\}, t_c^-) - R_c\right]^2 + \sum_{i=1}^{N_{bin}} \left(\frac{R(\{p\}, t_i) - R_{bin}(t_i)}{R_{bin}(t_i)}\right)^2 + \sum_{\alpha=1}^{N_q} \left(\frac{q(\{p\}, t_\alpha/\tau) - \bar{q}_\alpha}{q^*}\right)^2, \tag{1.51}$$

where $\{p\}$ is a set of value of the constitutive equations parameters and $R(\{p\}, t)$ the value of the radius at time $t$ when its dynamics is solved numerically with parameters value $\{p\}$. $R_{bin}(t_i)$ is the average value of experimental radius value at time $t \in [t_i - 3h, t_i + 3h]$, except for $t_i = 0$ where $R_{bin}(0)$ corresponds to the average value of experimental radius values just after the pressure increase. We took $N_{bin} = 9$ and $t_i = 0, 3, 7, 10, 24, 28, 32, 47, 56$ hours.

We find that the model still accounts fairly well for experimental data, and that the tube radius does not reach a steady-state anymore (Fig. S5A). The fitted value $\tau_a \approx 134$h is close to the inverse division rate $1/k_d \approx 100$h, the characteristic time at which cell division should relax the strain memory of actin fibers. This suggests that taking actin stress fibers as behaving elastically is a reasonable simplification on a time scale of 56 hours.

##### 1.3.3 Elastic behaviour of residual tension

The constitutive equation on residual tension (Eq. 1.19) corresponds to a viscoelastic response. The short-time elastic response of $t^r_{ij}$ ensures that just after pressure application, the tissue elongation in the circumferential direction leads to higher tension in that direction, in agreement with laser ablation experiments (Fig. 1Bii). We find that these elastic stresses $t^r_{ij}$ also have to relax to explain experimental data: indeed the reorientation of actin fibers after pressure increase leads to higher circumferential tension and would lead to a decrease in the tube radius, if the residual elastic tension had not relaxed (Fig. S5Ci).

We also tested the possibility that the residual tension $t^r_{ij}$ responds elastically, while the elastic stresses in actin fibers $t^a_{ij}$ relax over time. To test this possibility, we fitted the experimental time evolution of actin elongation by an exponential function $q(t) = a + b(1 - e^{-ct})$ (Fig. S5Cii), and numerically solved the evolution of the tube radius for a purely elastic response of the residual tension:

$$t^r_{ij} = K \ln(R/R_0), \tag{1.52}$$

with $K$ an elastic modulus and $R_0$ the value of the radius before pressure increase. We also assumed that $t^a_{ij}$ follows the viscoelastic constitutive equation discussed in section 1.3.2 (Eqs. 1.49 and 1.50). We fitted the radius experimental value by finding optimal values of the parameters through a Nelder-Mead algorithm to minimize the following cost function:

$$C(\{p\}) = \sum_{i=1}^{N_{bin}} \left(\frac{R(\{p\}, t_i) - R_{bin}(t_i)}{R_{bin}(t_i)}\right)^2. \tag{1.53}$$

The model we obtain then reproduced correctly the radius dynamics (Fig. S5Ci), but also predicted higher longitudinal tension just after pressure increase (Fig. S5Ciii), in contradiction with results from laser ablation experiments (Fig. 1Bii).

##### 1.3.4 Effect of tube curvature on actin stress fibers orientation

We tested the effect of adding a curvature coupling term $-\lambda_c \tilde{C}_{ij}$ in Eq. 1.11. Such a term would correspond to actin fibers orienting along the axis of minimal curvature, possibly through bending energy minimization. This could explain why actin fibers preferentially align along the tube axis before the application of the higher pressure. Going through the fitting procedure with this additional term, we find a small dimensionless value $\hat{\lambda}_c = \tau\lambda_c/R_0 = 6.10^{-5} \ll 1$. This suggests that the effect of curvature on fiber alignment is small compared to other effects, but is sufficient to select a preferred orientation $q = -q_0$ at tissue confluence.

##### 1.3.5 Coupling of actin stress fiber orientation dynamics on total tension

Here we discuss the coupling term with tension in the dynamics of actin mean orientation (Eq. 1.11). We assumed that this coupling depends on $t^r_{ij}$, the residual tension of the tissue without actin. We discuss here an alternative choice, introducing a coupling to the total tissue tension $t_{ij}$:

$$D_t q_{ij} = -\gamma \left(\frac{1}{2} q_{kl} q^{kl} - q_0^2\right) q_{ij} + \beta \tilde{t}_{ij}, \tag{1.54}$$

with $\tilde{t}_{ij} = t_{ij} - t^k_k g_{ij}/2$. The constitutive equations for $t^a_{ij}$ and $t^r_{ij}$, Eqs. 1.16, 1.17 and 1.19, are unchanged. The dimensionless system of equations then becomes:

$$R\Delta P = \sigma + (1 + K_a s)\left(q + \frac{1}{2}\right), \tag{1.55}$$

$$\left(1 + \frac{\mathrm{d}}{\mathrm{d}t}\right)\sigma = \frac{\mu}{R}\frac{\mathrm{d}R}{\mathrm{d}t}, \tag{1.56}$$

$$\frac{\mathrm{d}s}{\mathrm{d}t} = \left(q + \frac{1}{2}\right)\frac{1}{R}\frac{\mathrm{d}R}{\mathrm{d}t}, \tag{1.57}$$

$$\frac{\mathrm{d}q}{\mathrm{d}t} = -\gamma\left(q^2 - q_0^2\right) q + \beta\left[\sigma/2 + (1 + K_a s) q\right]. \tag{1.58}$$

We then looked for the optimal set of parameters for this model to account for experimental data, by using a Nelder-Mead algorithm to minimize the cost function defined in Eq. 1.38. This version of the model still shows good agreement with experimental data (Fig. S5Di,ii). However, the radius evolution exhibits unexpected behaviour for pressures smaller than $\simeq$ 350Pa. Indeed for a small pressure increase at $t = 0$, the initial circumferential tension anisotropy is not sufficient to drive actin fibers orientation circumferentially before $t^r_{ij}$ relaxes. The residual tension then decreases, the fibers are still longitudinal and, as the total tension arises mostly from actin stress fibers, the total tension anisotropy becomes longitudinal (Fig. S5Diii). As a result, $q$ decreases (Fig. S5Dii), meaning that fibers become more longitudinal. The actin network is thus not able to resist the circumferential tube expansion, and the tube radius significantly increases (Fig. S5Di). This leads to a new increase in $t^r_{ij}$, the tension anisotropy becomes circumferential, allowing at last fibers to reorient circumferentially and the radius to stabilize to a steady value that is higher than steady-state values reached for higher pressure (Fig. S5Di).

We also observe that at intermediate pressures between 150 Pa and 650Pa, the model with mechanosensing coupling on $t^r_{ij}$ exhibits a non-monotonous tube radius expansion as a function of pressure, due to a delay in fiber reorientation. The effect is however less pronounced that for the coupling with $t_{ij}$. As we lack experimental data at these intermediate pressures, we can not conclude at this point between the two models.

##### 1.3.6 Model of the tissue as an isotropic, viscoelastic material

In the model, the tension generated inside the tissue is the combination of an active and an elastic tension, oriented along actin stress fibers, and a residual viscoelastic response (Eqs 1.16, 1.17 and 1.19). Here we ask if a model where the tissue effectively behaves as an isotropic material, and in particular where tension is not oriented along actin stress fibers, can still explain experimental measurements. Therefore we consider the tension in the tissue to be given by a Zener model (combination of a Maxwell model in parallel with a spring, Fig. S5Eiv):

$$t_{ij} = t^r_{ij}, \tag{1.59}$$

$$t^r_{ij} = t^{el}_{ij} + t^{ve}_{ij}, \tag{1.60}$$

$$D_t t^{el}_{ij} = K v_{ij}, \tag{1.61}$$

$$(1 + \tau D_t) t^{ve}_{ij} = \mu v_{ij}. \tag{1.62}$$

We assume that for $R = R_c$ the tension $t^{el}_{ij}$ vanishes, $t^{el}_{ij} = 0$. In this subsection, we use dimensional equations. This results in the following equation for the tube radius:

$$\left(1+\tau\frac{\mathrm{d}}{\mathrm{d}t}\right)(R\Delta P) = \frac{\mu+\tau K}{R}\frac{\mathrm{d}R}{\mathrm{d}t} + K\ln\left(\frac{R}{R_c}\right). \tag{1.63}$$

We looked for optimal parameters by using a Nelder-Mead algorithm to minimize the following cost function

$$C(\{p\}) = A \times \left[R(\{p\}, t_c^-) - R_c\right]^2 + \sum_{i=1}^{N_R}\left(\frac{R(\{p\}, t_i/\tau) - \bar{R}_i/R_0}{\bar{R}_i/R_0}\right)^2. \tag{1.64}$$

The time evolution of the radius obtained from this fitted procedure exhibits a different qualitative behaviour than the mean curve of radius evolution experimental points (Fig. S5Ei), showing an inverse concavity compared to experimental data.

**Additional active isotropic tension.** We now consider the same model, but adding an isotropic active tension $\zeta_0 g_{ij}$. The constitutive equations of the model become:

$$t_{ij} = t^r_{ij} + \zeta_0 g_{ij}\,, \tag{1.65}$$

$$t^r_{ij} = t^{el}_{ij} + t^{ve}_{ij}\,, \tag{1.66}$$

$$D_t t^{el}_{ij} = K v_{ij}\,, \tag{1.67}$$

$$(1+\tau D_t)t^{ve}_{ij} = \mu v_{ij}\,, \tag{1.68}$$

which leads to the following equation for the tube radius evolution:

$$\left(1+\tau\frac{\mathrm{d}}{\mathrm{d}t}\right)(R\Delta P - \zeta_0) = \frac{\mu+\tau K}{R}\frac{\mathrm{d}R}{\mathrm{d}t} + K\ln\left(\frac{R}{R_c}\right), \tag{1.69}$$

We then look for optimal parameters by using a Nelder-Mead algorithm to minimize the cost function defined in Eq. 1.64. A better agreement is found with experimental data (Fig. S5Ei), as expected from the addition of an extra parameter. However, the radius obtained from the model reaches a plateau about 24 hours after pressure increase, in contrast to experiments where the radius keeps increasing during that period. Moreover it predicts a large circumferential tension anisotropy at negative times that contradicts observation in laser ablation experiments.

##### 1.3.7 Isotropic viscoelasticity with anisotropic active tension

Here we ask whether a model where the long-time scale elastic response is dependent on tissue deformation, but not on the orientation of actin stress fibers, can still account for experimental data. We consider the same Zener model than in the previous section, with the addition of an active anisotropic tension $\zeta_0 n_{ij}$ oriented along actin fibers. The constitutive equations then become (here using dimensional parameters):

$$t_{ij} = t^a_{ij} + t^r_{ij}\,, \tag{1.70}$$

$$t^a_{ij} = \zeta_0 n_{ij}\,, \tag{1.71}$$

$$t^r_{ij} = t^{el}_{ij} + t^{ve}_{ij}\,, \tag{1.72}$$

$$D_t t^{el}_{ij} = K v_{ij}\,, \tag{1.73}$$

$$(1+\tau D_t)t^{ve}_{ij} = \mu v_{ij}\,, \tag{1.74}$$

$$D_t q_{ij} = -\gamma\left(\frac{1}{2}q_{kl}q^{kl} - q_0^2\right)q_{ij} + \beta\tilde{t}^r_{ij}\,. \tag{1.75}$$

This results in the following coupled dynamics for the tube radius and actin nematic $q$:

$$\left(1+\tau\frac{\mathrm{d}}{\mathrm{d}t}\right)\left(R\Delta P - \zeta_0\left(q+\frac{1}{2}\right)\right) = \frac{\mu+\tau K}{R}\frac{\mathrm{d}R}{\mathrm{d}t} + K\ln\left(\frac{R}{R_c}\right), \tag{1.76}$$

$$\frac{\mathrm{d}q}{\mathrm{d}t} = -\gamma\left(q^2 - q_0^2\right)q + \frac{1}{2}\beta\left(R\Delta P - \zeta_0(q + \frac{1}{2})\right), \tag{1.77}$$

We optimized the parameters of this model using a Nelder-Mead algorithm to minimize the cost function introduced in Eq. 1.38. The model is in good agreement with experimental data for radius and actin stress fiber nematic for $t > 0$ (Fig. S5Ei,ii). The tension anisotropy is however less consistent with laser ablation experiments than the model presented in main text: indeed at negative times, the tension is larger along the circumferential direction, and the longitudinal tension becomes negative at positive time (Fig. S5Eiii). In addition, the behaviour of the model at negative times is less comparable to experiments, with a large jump after confluence followed by a decrease in the tube radius.

Despite these differences, this version of the model appears still qualitatively acceptable. We conclude that an active tension oriented along actin fibers is a key aspect of the model to reproduce experimental data.

<table>
<tr><th>Actin mechanosensing</th><th>Residual tension</th><th>Actin oriented tension</th><th>Results</th></tr>
<tr><td rowspan="7">Residual tension</td><td rowspan="2">Purely elastic</td><td>Purely elastic<br>+<br>Active tension</td><td>Radius decrease after initial jump<br>(Sec. 1.3.3, Fig. S5Ci, green curve)</td></tr>
<tr><td rowspan="2">Viscoelastic<br>+<br>Active tension</td><td>Tension anisotropy after pressure increase disagrees with experiment (Sec. 1.3.3, Fig. S5Ci, red curve, Fig. S5Ciii)</td></tr>
<tr><td rowspan="2">Viscoelastic</td><td>Good agreement<br>with experimental data<br>(Sec. 1.3.2, Fig. S5A)</td></tr>
<tr><td>Purely elastic<br>+<br>Active tension</td><td>Good agreement<br>with experimental data<br>(Sec. 1.1.6, Fig. 4)</td></tr>
<tr><td>Zener model</td><td rowspan="2">None</td><td>Dynamics of radius does not<br>agree with experimental data<br>(Sec. 1.3.6, Fig. S5Ei, green curve)</td></tr>
<tr><td>Zener model<br>+<br>Isotropic active tension</td><td>Radius plateau at 24hours<br>(Sec. 1.3.6, Fig. S5Ei, dashed red curve)</td></tr>
<tr><td>Zener model</td><td>Active tension</td><td>Good agreement for radius and actin orientation dynamics but tension behaviour not in good agreement with laser ablation experiments<br>(Sec. 1.3.7, Fig. S5E)</td></tr>
<tr><td>Total tension</td><td>Viscoelastic</td><td>Purely elastic<br>+<br>Active tension</td><td>Dramatic radius increase at low pressures (Sec. 1.3.5, Fig. S5D)</td></tr>
</table>

Table 1.4: Summary of model variations. Choices corresponding to the main model are indicated in bold.

### Part 2 Mechanical model of the endothelial tube instantaneous strain-stiffening

In the following, we present the isotropic nonlinear elastic Gent model, that we then use to characterize the deformation of the endothelial tubes under applied pressure. The contribution of the deformed collagen material surrounding the tube is accounted for, to determine the pressure differential across the cell layer. The experimental data are fit to this model with an orthogonal distance regression, to determine elastic moduli and maximum extension of the nonlinear tissue. The effect of the collagen concentration on the tissue properties are investigated, and the contribution of the actin cytoskeleton and cell-cell junctions are discussed.

#### 2.1 Model of the endothelial tube

##### 2.1.1 Gent model

The cell monolayer is modeled as a nonlinear elastic cylindrical shell, using the Gent model, surrounded by a linear elastic collagen medium [Gent, 1996, Salipante et al., 2022]. The Gent model is an empirical model for hyperelasticity, that captures the quasi-linear response at small strains and strain hardening behavior at larger strain. The model relates the pressure drop across the cell layer as a function of the extension of the vessel radius, defined as $\lambda = \frac{R}{R_0}$:

$$\Delta P = -\frac{hJ_m\left(\lambda^4 - 1\right)\mu}{R_0\lambda^2(1-(2+J_m)\lambda^2+\lambda^4)} \tag{2.1}$$

where $h$ is the monolayer thickness, $\mu$ the shear elastic modulus and $J_m$ the extensibility parameter. Assuming the material is isotropic, the shear modulus can then be used to compute the Young's modulus by

$$E = 2\mu(1+\nu), \tag{2.2}$$

where $\nu$ is the Poisson's ratio of the endothelial tissue, assumed to be 0.5; while the extensibility can be used to define a maximum extension by

$$\lambda_{max} = \sqrt{(2+J_m+\sqrt{J_m}\sqrt{4+J_m})/2}. \tag{2.3}$$

##### 2.1.2 Contribution of the surrounding collagen gel

In our experiments, both the cell monolayer and the surrounding collagen resist deformation induced by the applied luminal pressure, $P_{lum}$. The total pressure drop between the lumen and the outside applied pressure equals the sum of the pressure drop across the cell layer, $\Delta P$, and across the collagen matrix, $\Delta P_g$. We are interested in the elastic response of the cell layer, which depends on the pressure drop across the layer. The pressure drop across the cell layer is then $\Delta P = P_{lum} - \Delta P_{gel}$.

To determine the pressure at the interface between the cell layer and collagen, we need to account for the deformation of the collagen. We model the collagen as a linear elastic material surrounding a cylindrical inclusion. The circumferential strain of the cylinder results in a pressure at surface of the surrounding collagen, that decays to a zero reference pressure far from the cylinder. The pressure exerted by the surrounding collagen is determined using the following equation:

$$\Delta P_c = \frac{E_{gel}\epsilon}{(1+\nu_{gel})} \tag{2.4}$$

where $E_{gel}$ is the Young's modulus of the collagen and the strain is $\epsilon = \lambda - 1$. The Poisson's ratio of the collagen gel $\nu_{gel}$ is assumed to be 0.5.

The elastic modulus of collagen varies significantly with collagen concentration, following a power law [Jansen et al., 2018]. The modulus of the 2 mg/mL collagen has been measured to be around 50 Pa and the modulus of the 6 mg/mL collagen around 650 Pa. The contribution of collagen to the pressure drop is calculated from Eq. 2.4 using the measured strain, allowing us to determine the pressure drop across the cell layer. The contribution of the collagen gel to the pressure drop is largest at small deformation, but is still small compared to the cell layer [Salipante et al., 2022]. Assuming a linear elastic response of the cell layer, the relative contribution of the collagen layer to the pressure drop is approximately $\Delta P_{gel}/P_{lum} \sim 10$ %.

##### 2.1.3 Fitting procedure

The thickness of the monolayer $h$ is observed on fluorescent images of actin immunostainings to vary within the tissue, being maximal above cell nuclei and minimal in between nuclei. By delineating manually the monolayer contour on several images, the average thickness is measured to be $h = 3.6 \pm 0.5\mu m$.

Using the continuous recording of the circumferential strain as a function of pressure (see Methods), the shear modulus $\mu$, extensibility parameter $J_m$, and reference state $R_0$, are determined by fitting individual strain-stress curve with orthogonal distance regression (ODR) using the Scipy package in Python [Boggs and Rogers, 1990]. The fitted curves are shown in Fig. 1Ci, Di, and the corresponding parameters are shown in Tables in the following section. The uncertainty is reported with the standard error on the fitted parameters.

#### 2.2 Results

##### 2.2.1 Substrate-dependent properties of endothelial tubes

We first sought to measure the mechanical properties of our endothelial tubes, and to check for potential substrate dependencies due to cellular mechanosensing. To that end, we templated cellular monolayers inside two different collagen concentrations, 2 mg/mL and 6 mg/mL, thereby tuning the substrate ridigity.

**Reference state** $R_0$ The experimental data from monolayers on both collagen concentrations displays a significant portion of quasi-linear deformation, whose fit can be extrapolated to infer the reference state $R_0$ corresponding to the radius at $\Delta P = 0$. This value is found to be slightly above the needle radius (60$\mu$m) used to fabricate the channels, $R_0 = 62.5\mu$m, matching the diameter of $125\mu m$ of bare channels reported in our previous work [Dessalles et al., 2021].

**Young's modulus** $E$ The elastic moduli characterize the linear mechanical response of the cell layer at low applied pressures. We find that the inferred elastic modulus changes significantly with surrounding collagen concentration, a signature of cellular mechanosensitivity. The measured modulus of the cell layer templated in 6 mg/mL collagen is 128 ± 24 kPa, significantly greater than that of the layer templated in 2 mg/mL collagen 28 ± 3 kPa. This difference does not come from the different collagen stiffnesses, as the contribution of the gel to the response is accounted for in the calculation. We hypothesize that the difference is due to reinforcement of the actin network into thick bundles of stress fibers anchored at large focal adhesions, triggered by mechanosensing of the collagen properties (see Main text, Fig. 1C-D).

**Maximal extension** $\lambda_{max}$ The strain stiffening response is determined by the extensibility parameter $J_m$ of the Gent model in Eq. 2.1, but is most easily understood by the asymptotic extension value of the maximum extension, $\lambda_{max}$. We find that the inferred maximum extension are $1.36 \pm 0.06$ and $1.23 \pm 0.1$ for the tissues on the 2 mg/mL and 6 mg/mL collagen respectively. The two maximum extensions are not significantly different and could be defined by subcellular components which show no or a weaker adaptation to substrate properties.

##### 2.2.2 Subcellular determinants of endothelium properties.

To probe the role of the actin cytoskeleton and cell-cell junctions in the tissue properties, we treated the monolayers on 2 mg/mL collagen with cytochalasinD, to depolymerize actin, and EDTA, to perturb adherens

| Device # | Radius $R_0$ ($\mu$m) | Shear modulus $\mu$ (kPa) | Young's modulus $E$ (kPa) | Extensibility Parameter $J_m$ | Maximum extension $\lambda_{max}$ |
|---|---|---|---|---|---|
| 1 | 63.2 ± 0.3 | 10.6 ± 0.5 | 31.8 ± 1.6 | 0.273 ± 0.008 | 1.295 ± 0.005 |
| 2 | 64.2 ± 0.3 | 9.3 ± 0.4 | 27.9 ± 1.0 | 0.412 ± 0.008 | 1.371 ± 0.004 |
| 3 | 62.4 ± 0.3 | 8.7 ± 0.4 | 26.3 ± 1.1 | 0.532 ± 0.013 | 1.429 ± 0.006 |

Table 2.1: Fitted parameters for endothelial tubes templated in 2 mg/mL collagen. Uncertainties represent standard error on the parameter estimates from ODR.

| Device # | Radius $R_0$ ($\mu$m) | Shear modulus $\mu$ (kPa) | Young's modulus $E$ (kPa) | Extensibility Parameter $J_m$ | Maximum extension $\lambda_{max}$ |
|---|---|---|---|---|---|
| 1 | 62.6 ± 0.2 | 50.9 ± 3.4 | 152.8 ± 10.1 | 0.151 ± 0.025 | 1.213 ± 0.019 |
| 2 | 60.4 ± 0.2 | 41.9 ± 3.4 | 125.9 ± 10.1 | 0.064 ± 0.006 | 1.134 ± 0.006 |
| 3 | 62.5 ± 0.2 | 34.6 ± 0.8 | 103.9 ± 10.4 | 0.398 ± 0.052 | 1.363 ± 0.027 |

Table 2.2: Fitted parameters for endothelial tubes templated in 6 mg/mL collagen. Uncertainties represent standard error on the parameter estimates from ODR.

junctions, prior to the stress-strain curve measurement. Because both treatments significantly softened the tissue (see below), the circumferential strain was increased and the initial low pressure of 150 Pa was sufficient to exceed the linear regime (Fig. 1E). Therefore, the reference state $R_0 = 62.5\mu m$ determined above was used as a fixed parameter in the fits.

**Actin depolymerization** We find the Young's modulus of cytochalasinD-treated tissues to be 11 ± 2 kPa, significantly smaller than that of the untreated tissues, 28 ± 3 kPa. This confirms that the actin network is a major contributor to tissue elasticity. Interestingly, the maximum extension is inferred to be 1.33 ± 0.02, similar to the untreated tissues at 1.36 ± 0.06. This suggests that the actin network contribution to the maximum extension is minimal, in line with the observation that the maximum extension is not substrate-sensitive (see above). Other subcellular components are thus driving the strain-stiffening properties of tissues found at larger pressures, with intermediate filaments being a likely candidate [Latorre et al., 2018, Singh et al., 2018].

| Device # | Shear modulus $\mu$ (kPa) | Young's modulus $E$ (kPa) | Extensibility Parameter $J_m$ | Maximum extension $\lambda_{max}$ |
|---|---|---|---|---|
| 1 | 3.1 ± 0.1 | 9.4 ± 0.3 | 0.374 ± 0.005 | 1.351 ± 0.003 |
| 2 | 3.8 ± 0.1 | 11.4 ± 0.4 | 0.386 ± 0.006 | 1.358 ± 0.003 |
| 3 | 4.2 ± 0.2 | 12.6 ± 0.5 | 0.313 ± 0.006 | 1.318 ± 0.003 |

Table 2.3: Fitted parameters for for monolayers on 2 mg/mL collagen and treated with cytochalasinD. An initial radius of $R_0 = 62.5$ $\mu$m is assumed for all data. Uncertainties represent standard error on the parameter estimates from ODR.

**Junctions perturbation** We find the Young's modulus of EDTA-treated tissues to be 12±4 kPa, significantly smaller than that of the untreated tissues, 28 ± 3 kPa. This could be due to softer cells or to the stretch of the soft collagen between the now-disjointed cells. The maximum extension is inferred to be 1.53 ± 0.06, significantly larger than that of the control tissues, at 1.36 ± 0.06. Similarly, this could be due to a modification of the cell properties or to the stretch of the collagen gel.

| Device # | Shear modulus $\mu$ (kPa) | Young's modulus $E$ (kPa) | Extensibility Parameter $J_m$ | Maximum extension $\lambda_{max}$ |
|---|---|---|---|---|
| 1 | 3.1 ± 0.1 | 9.3 ± 0.2 | 0.908 ± 0.011 | 1.584 ± 0.004 |
| 2 | 5.0 ± 0.1 | 14.9± 0.4 | 0.673 ± 0.013 | 1.491 ± 0.005 |

Table 2.4: Fitted parameters for monolayers on 2 mg/mL collagen and treated with EDTA. An initial radius of $R_0 = 62.5\,\mu$m is assumed for all data.

### Bibliography

[Ban et al., 2018] Ban, E., Franklin, J. M., Nam, S., Smith, L. R., Wang, H., Wells, R. G., Chaudhuri, O., Liphardt, J. T., and Shenoy, V. B. (2018). Mechanisms of plastic deformation in collagen networks induced by cellular forces. *Biophysical Journal*, 114:450–461.

[Boggs and Rogers, 1990] Boggs, P. T. and Rogers, J. E. (1990). Orthogonal distance regression. *Contemporary mathematics*, 112:183–194.

[Dessalles et al., 2021] Dessalles, C. A., Ramon-Lozano, C., Babataheri, A., and Barakat, A. I. (2021). Luminal flow actuation generates coupled shear and strain in a microvessel-on-chip. *Biofabrication*, 14(1):015003.

[Duque et al., 2023] Duque, J., Bonfanti, A., Fouchard, J., Baldauf, L., Azenha, S. R., Ferber, E., Harris, A., Barriga, E., Kabla, A., and Charras, G. (2023). Rupture strength of living cell monolayers. *bioRxiv*.

[Etournay et al., 2015] Etournay, R., Popović, M., Merkel, M., Nandi, A., Blasse, C., Aigouy, B., Brandl, H., Myers, G., Salbreux, G., Jülicher, F., et al. (2015). Interplay of cell dynamics and epithelial tension during morphogenesis of the drosophila pupal wing. *Elife*, 4:e07090.

[Gent, 1996] Gent, A. (1996). A new constitutive relation for rubber. *Rubber chemistry and technology*, 69(1):59–61.

[Gong et al., 2015] Gong, M., Yang, H., Zhang, S., Yang, Y., Zhang, D., Qi, Y., and Zou, L. (2015). Superparamagnetic core/shell goldmag nanoparticles: Size-, concentration- and time-dependent cellular nanotoxicity on human umbilical vein endothelial cells and the suitable conditions for magnetic resonance imaging. *Journal of Nanobiotechnology*, 13.

[Harris et al., 2012] Harris, A. R., Peter, L., Bellis, J., Baum, B., Kabla, A. J., and Charras, G. T. (2012). Characterizing the mechanics of cultured cell monolayers. *Proceedings of the National Academy of Sciences*, 109(41):16449â16454.

[Hayakawa et al., 2001] Hayakawa, K., Sato, N., and Obinata, T. (2001). Dynamic reorientation of cultured cells and stress fibers under mechanical stress from periodic stretching. *Experimental Cell Research*, 268:104–114.

[Iba and Sumpio, 1991] Iba, T. and Sumpio, B. E. (1991). Morphological response of human endothelial cells subjected to cyclic strain in vitro. *Microvascular Research*, 42:245–254.

[Jansen et al., 2018] Jansen, K. A., Licup, A. J., Sharma, A., Rens, R., MacKintosh, F. C., and Koenderink, G. H. (2018). The role of network architecture in collagen mechanics. *Biophysical Journal*, 114:2665.

[Kaunas et al., 2005] Kaunas, R., Nguyen, P., Usami, S., and Chien, S. (2005). Cooperative effects of rho and mechanical stretch on stress fiber organization. *Proceedings of the National Academy of Sciences of the United States of America*, 102:15895–15900.

[Kim et al., 2017] Kim, J., Feng, J., Jones, C. A., Mao, X., Sander, L. M., Levine, H., and Sun, B. (2017). Stress-induced plasticity of dynamic collagen networks. *Nature Communications 2017 8:1*, 8:1–7.

[Krishnan et al., 2012] Krishnan, R., Canović, E. P., Iordan, A. L., Rajendran, K., Manomohan, G., Pirentis, A. P., Smith, M. L., Butler, J. P., Fredberg, J. J., and Stamenović, D. (2012). Fluidization, resolidification, and reorientation of the endothelial cell in response to slow tidal stretches. *American Journal of Physiology-Cell Physiology*, 303(4):C368–C375. PMID: 22700796.

[Latorre et al., 2018] Latorre, E., Kale, S., Casares, L., GÃ$^{3}mez - Gonzlez, M., Uroz, M., Valon, L., Nair, R.V., Garreta, E., Montserrat, N., delCampo, A., andet al.(2018).Actives dimensionalepitheliaofcontrolledshape.Nature, 563(7730) : 203208.$

[Marin-Llaurado et al., 2023] Marin-Llaurado, A., Kale, S., Ouzeri, A., Golde, T., Sunyer, R., Torres-Sanchez, A., Latorre, E., Gomez-GonzÃ¡lez, M., Roca-Cusachs, P., Arroyo, M., and Trepat, X. (2023). Mapping mechanical stress in curved epithelia of designed size and shape. *Nature Communications 2023 14:1*, 14:1–11.

[Nam et al., 2016] Nam, S., Lee, J., Brownfield, D. G., and Chaudhuri, O. (2016). Viscoplasticity enables mechanical remodeling ofÂ matrix by cells. *Biophysical Journal*, 111:2296–2308.

[Popović et al., 2017] Popović, M., Nandi, A., Merkel, M., Etournay, R., Eaton, S., Jülicher, F., and Salbreux, G. (2017). Active dynamics of tissue shear flow. *New Journal of Physics*, 19(3):033006.

[Pourati et al., 1998] Pourati, J., Maniotis, A., Spiegel, D., Schaffer, J. L., Butler, J. P., Fredberg, J. J., Ingber, D. E., Stamenovic, D., and Wang, N. (1998). Is cytoskeletal tension a major determinant of cell deformability in adherent endothelial cells? *American Journal of Physiology - Cell Physiology*, 274.

[Salipante et al., 2022] Salipante, P. F., Hudson, S. D., and Alimperti, S. (2022). Blood vessel-on-a-chip examines the biomechanics of microvasculature. *Soft Matter*, 18(1):117â125.

[Singh et al., 2018] Singh, A., Saha, T., Begemann, I., Ricker, A., NÃ¼sse, H., Thorn-Seshold, O., Klingauf, J., Galic, M., and Matis, M. (2018). Polarized microtubule dynamics directs cell mechanics and coordinates forces during epithelial morphogenesis. *Nature Cell Biology*, 20(10):1126â1133.

[Takemasa et al., 1997] Takemasa, T., Sugimoto, K., and Yamashita, K. (1997). Amplitude-dependent stress fiber reorientation in early response to cyclic strain. *Experimental Cell Research*, 230:407–410.

[Vasiceek et al., 2021] Vasiceek, J., Balazi, A., Bauer, M., Svoradova, A., Tirpakova, M., Tomka, M., and Chrenek, P. (2021). Molecular profiling and gene banking of rabbit epcs derived from two biological sources. *Genes*, 12:1–27.
