## Supplementary figures and images for "Interplay of actin nematodynamics and anisotropic tension controls endothelial mechanics"

### SV1

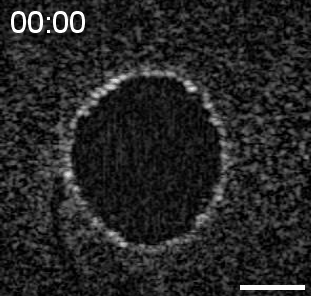

### SV2

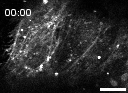

### SV3

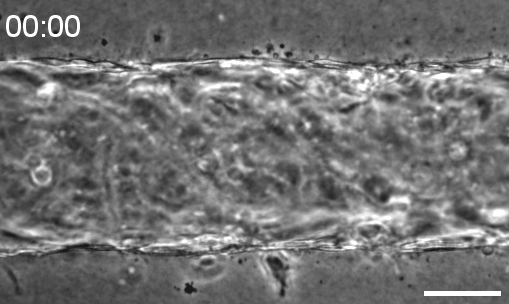

### SV4

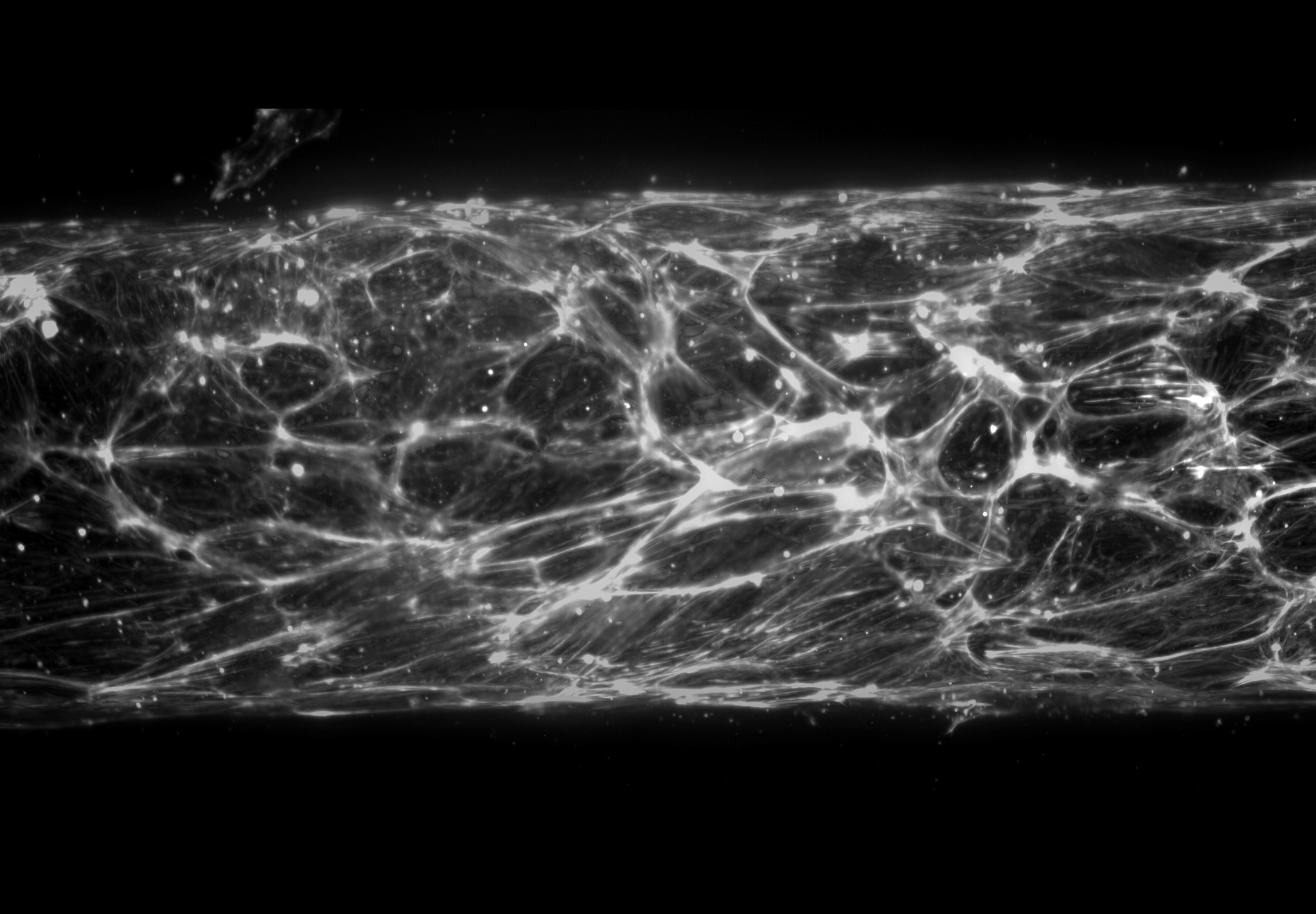

### SV5

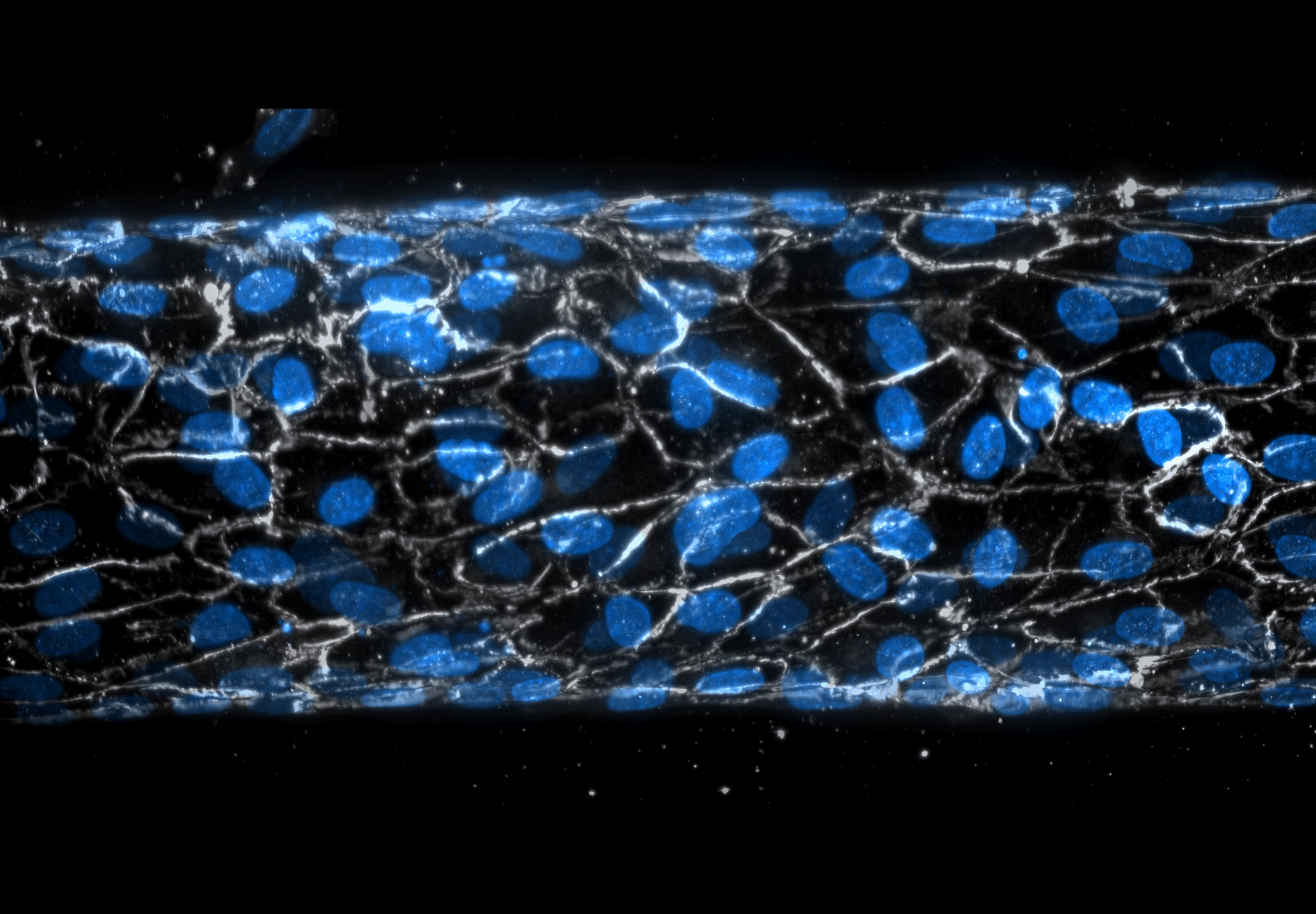
